# Cell of origin alters myeloid-mediated immunosuppression in lung adenocarcinoma

**DOI:** 10.1101/2024.06.19.599651

**Authors:** Minxiao Yang, Noah Shulkin, Edgar Gonzalez, Jonathan Castillo, Chunli Yan, Keqiang Zhang, Leonidas Arvanitis, Zea Borok, W. Dean Wallace, Dan Raz, Evanthia T. Roussos Torres, Crystal N. Marconett

## Abstract

Solid carcinomas are often highly heterogenous cancers, arising from multiple epithelial cells of origin. Yet, how the cell of origin influences the response of the tumor microenvironment is poorly understood. Lung adenocarcinoma (LUAD) arises in the distal alveolar epithelium which is populated primarily by alveolar epithelial type I (AT1) and type II (AT2) cells. It has been previously reported that *Gramd2*^+^ AT1 cells can give rise to a histologically-defined LUAD that is distinct in pathology and transcriptomic identity from that arising from *Sftpc^+^* AT2 cells^1,2^. To determine how cells of origin influence the tumor immune microenvironment (TIME) landscape, we comprehensively characterized transcriptomic, molecular, and cellular states within the TIME of *Gramd2*^+^ AT1 and *Sftpc*^+^ AT2-derived LUAD using KRAS^G12D^ oncogenic driver mouse models. Myeloid cells within the *Gramd2^+^*AT1-derived LUAD TIME were increased, specifically, immunoreactive monocytes and tumor associated macrophages (TAMs). In contrast, the *Sftpc*^+^ AT2 LUAD TIME was enriched for Arginase-1^+^ myeloid derived suppressor cells (MDSC) and TAMs expressing profiles suggestive of immunosuppressive function. Validation of immune infiltration was performed using flow cytometry, and intercellular interaction analysis between the cells of origin and major myeloid cell populations indicated that cell-type specific markers SFTPD in AT2 cells and CAV1 in AT1 cells mediated unique interactions with myeloid cells of the differential immunosuppressive states within each cell of origin mouse model. Taken together, *Gramd2*^+^ AT1-derived LUAD presents with an anti-tumor, immunoreactive TIME, while the TIME of *Sftpc*^+^ AT2-derived LUAD has hallmarks of immunosuppression. This study suggests that LUAD cell of origin influences the composition and suppression status of the TIME landscape and may hold critical implications for patient response to immunotherapy.

## INTRODUCTION

Lung cancer has been the leading cause of cancer-associated death for decades in the U.S.^1^ and worldwide^2^. Lung adenocarcinoma (LUAD) is the most common subtype of lung cancer^3^, accounting for approximately 40% of cases^4,5^. LUAD is highly heterogenous in multiple aspects, including epigenetic landscape^6^, transcriptomic signatures^7^, histologic presentation^3^, and critically, patient response to immunotherapy^8^.

LUAD originates from distal airways and alveoli where multiple types of epithelial cell reside, including AT2, AT1, club, and bronchioalveolar stem cells (BASCs). These cell types are distinct in function and in cell marker expression both in vivo and in vitro. Previous research has demonstrated that mouse models driving expression of oncogenic Kristen RAt Sarcoma viral oncogene homolog (KRAS)-G12D amino acid mutation in *Sftpc*^+^ AT2 cells give rise to lesions histologically similar to human LUAD^9–11^. In addition, recent studies have established that *Gramd2*^+^ and *Hopx*^+^ AT1 cells can also give rise to KRAS^G12D^-driven LUAD^12,13^. However, how cell of origin influences the immune landscape within the tumor immune microenvironment (TIME) in LUAD is poorly understood.

Precision medicine has revolutionized cancer treatment in the past decade by specifically targeting genomic alterations in a patients’ tumor to improve survival. A major success in precision treatment for LUAD is widespread adoption of immune checkpoint inhibition (ICI) therapy, which has significantly altered the therapeutic paradigm, emerging as a first-line treatment option for LUAD patients over the past ten years^14^ with a dramatically improved patient response^15,16^. However, the collective clinical response rate to ICI therapy remains relatively modest, with only an estimated ∼20% of LUAD patients experiencing therapeutic benefits from ICI interventions^17–19^. Although the mechanisms underlying the observed differences in ICI response remain largely unknown, the dynamics of tumor immune microenvironment (TIME) around and within LUADs can be inferred as a significant contributor to the effectiveness of response^20–22^. Robust biomarkers predicting failure of ICI therapy is understudied^23,24^, and the underlying molecular basis for why some LUADs foster an immunoreactive TIME remains largely unknown.

Certain subsets of LUAD are known to have TIME enriched with natural killer (NK) cells, cytotoxic T cells, macrophages, and neutrophils, resulting in enhanced interferon, interleukin, and chemokine signaling^25–27^ that promote anti-tumor immune activity. In contrast, other subsets of LUADs adopt an immunosuppressive environment, characterized by an increase in immature dendritic cells (DCs), myeloid-derived suppressor cells (MDSCs), regulatory T cells (Tregs), and pro-tumoral tumor-associated macrophages (TAMs)^28–30^. The mechanisms that promote immune suppression and favor tumor growth are an active area of investigation in the field.

The myeloid immune cell population plays an instrumental role in response to ICI therapy within the TIME of lung adenocarcinoma. MDSCs and TAMs have distinct yet interrelated functions that contribute to a suppressive immune landscape, aiding tumor escape from immune surveillance. MDSCs and TAMs exert their suppressive mechanisms primarily through the production of inhibitory cytokines, such as interleukin-10 (IL-10) and through the expression of enzymes like arginase-1 (ARG1) and inducible nitric oxide synthase (iNOS)^33^. These enzymes deplete local supplies of amino acids essential for T cell function, such as arginine, and produce reactive oxygen species, which impair T cell receptor signaling and promote T cell apoptosis^34^. By suppressing the activation and proliferation of cytotoxic T cells via PD-1/PD-L1 pathways^35^, MDSCs and TAMs could also enable tumor cells to evade immune surveillance and even impact the efficacy of immunotherapy^36^. Therefore, understanding the mechanisms behind myeloid reprogramming within the immunosuppressive TIME is essential for bolstering immunotherapeutic efficacy for LUAD patients.

KRAS mutation can also regulate the TIME via recruitment of immature suppressive immune cells such as MDSCs^37–39^ through activation of the KRAS-IRF2-CXCL3-CXCR2 signaling axis. Strategies exploiting this relationship are currently under development as adjuvant ICI therapies^40^. Additionally, KRAS mutations are known to affect the immunosuppressive state of resident macrophages within the TIME of LUAD^41,42^, a process closely linked to the cancer’s cell of origin^42^. This inflammatory response is intricately linked to the interaction between alveolar macrophages and AT2 cells mediated by Connexin-43, which is highly expressed on AT2 cells of patient and murine KRAS^mutant^ LUADs and could facilitate crucial intercellular communication to promote immune tolerance^42,43^. KRAS is also involved in repair and differentiation conditions in alveolar cells. Previous studies have suggested that alveolar cell-type specific regulation of KRAS function, activation of endogenous, non-mutated KRAS in AT2 cells, facilitates stem cell niche renewal but non-mutated KRAS activation in AT1 cells can lead to apoptosis^44^.

Overall, alveolar cell of origin of LUAD may play an important role in how KRAS-mediated oncogenic transformation influences the immune phenotype of the TIME, which is critical to ICI response. Therefore, this study was designed to decipher the mechanisms by which alveolar cell of origin affects immune cell states within KRAS^mutant^ LUAD.

## RESULTS

### Single-nucleus RNA sequencing of *Sftpc^+^* AT2- and *Gramd2*^+^ AT1-derived KRAS^G12D^ mouse lung adenocarcinoma models reveal differential immune composition

To comprehensively characterize how cellular composition of the tumor microenvironment differs within LUAD based on the cell of origin in which the tumor arises, we performed single-nucleus RNA sequencing (snRNA-Seq) using the 10X Genomics Chromium platform on mouse models designed to activate the KRAS^G12D^ oncogenic driver in distinct alveolar epithelial cell populations (Figure 1a). Quality control measures to ensure only high-quality cells were included in downstream analysis were performed, which required filtration of samples with low gene counts (<150), low feature numbers (<100) and high mitochondrial gene percentage (>20%). Data processing included principal component analysis (PCA), and reduction of batch effects using harmony integration^45^ (Supplemental Figure 1). In total, 152,000 nuclei from fourteen mouse lung samples (Table 1) were profiled, including four negative controls, two vehicle controls for *Sftpc^CreERT2:^* KRAS^LSL-G12D^, two vehicle controls for *Gramd2^CreERT2:^* KRAS^LSL-G12D^, three *Sftpc^+^* (AT2-driven) KRAS^G12D^ LUAD samples, and three *Gramd2^+^*(AT1-driven) KRAS^G12D^ LUAD samples. Uniform manifold approximation and projection (UMAP) clustering of single-nucleus transcriptomes identified 16 major clusters of unique cellular identities (Figure 1b). The disease status, batch ID and individual mouse ID of each sample are included in Figure 1c and Supplemental Figure 1.

**Fig. 1.**
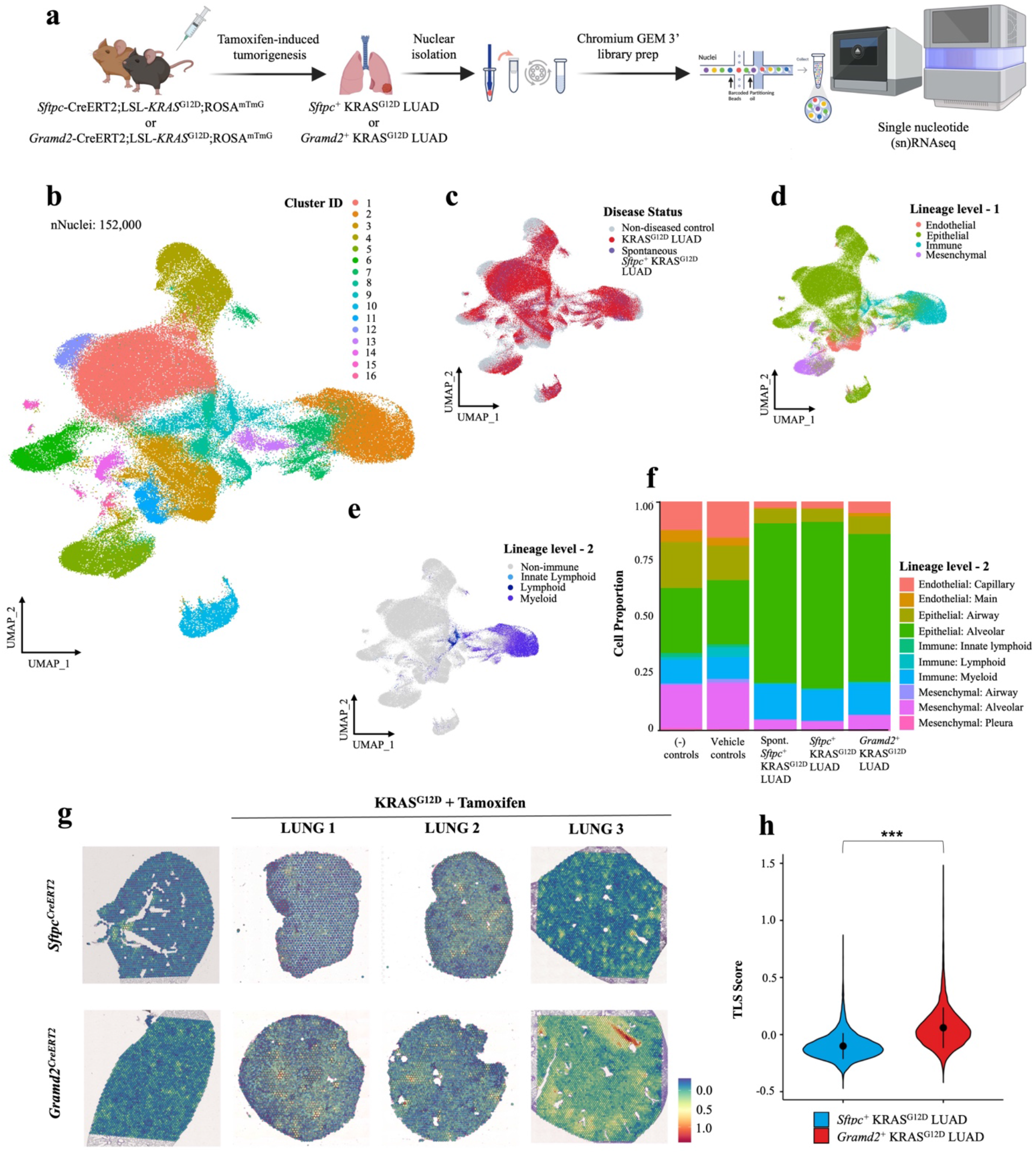
Single-nucleus RNA sequencing of *Sftpc^+^* AT2- and *Gramd2*^+^ AT1-derived KRAS^G12D^ mouse lung adenocarcinoma models reveal differential immune composition. a, Schematic representation of the overall workflow. *Sftpc*-CreERT2;LSL-KRAS^G12D^;ROSA^mTmG^ mouse model was used to established AT2 cell-derived lung adenocarcinoma (*Sftpc^+^* KRAS^G12D^ LUAD) and *Gramd2*-CreERT2;LSL-KRAS^G12D^;ROSA^mTmG^ mouse model was used to established AT1 cell-derived lung adenocarcinoma (*Gramd2^+^* KRAS^G12D^ LUAD) upon tamoxifen induction. Created with <u>BioRender.com</u>. b, Uniform manifold approximation and projection (UMAP) plot of single nucleus transcriptomic expression data. Cells colored by Seurat cluster ID (dims = 1:16; resolution = 0.3). c, UMAP plot of single nucleus transcriptomic expression data colored by disease status. Grey = healthy controls; red = LUAD samples; purple = spontaneous *Sftpc^+^*KRAS^G12D^ LUAD. d, UMAP plot of single nucleus transcriptomic expression data colored by LungMAP cell lineages (level1) predicted using Azimuth^47^. g, UMAP as plot of single nucleus transcriptomic expression data colored by LungMAP cell sub-lineages (level2) as predicted by Azimuth. The immune cell sub-lineages are indicated as follows; light blue = innate lymphoid cells, dark blue = lymphoid cells, and purple = myeloid cells. Non-immune cell sub-lineages are shown in grey. f, Overall cell composition of each experimental group is shown as a proportion of total profiled cells. Salmon = endothelial: capillary, gold = endothelial: main, peridot = epithelial: airway, green = epithelial: alveolar, seafoam = immune: innate lymphoid, teal = immune: lymphoid, blue = immune: myeloid, purple = mesenchymal: airway, magenta = mesenchymal: alveolar, and pink = mesenchymal: pleura. g, Spatial transcriptomic profiles of previously generated dataset^13^ overlaid with tertiary lymphoid structure (TLS) composite score. Red = high TLS score, blue = low TLS score. h, Violin plots of TLS scores between spatial transcriptomic profiling samples shown in (g) from *Sftpc^+^* KRAS^G12D^ LUAD (light blue) and *Gramd2^+^* KRAS^G12D^ LUAD models (red) across LUAD array samples. Wilcoxon rank-sum test was applied to test the significance of difference (p value < 2.2e-16). *p < 0.05, **p < 0.01, ***p < 0.001.

**Table 1:** Mouse experimental groups included in snRNAseq analysis. WT = wilt type, mTmG = Gt(ROSA)26Sor^tm4(ACTB-tdTomato,-EGFP)Luo^/J, TAM = Tamoxifen-treated, (-) = negative.

| Genotype | Sample size (n) | Tamoxifen Treatment | Mouse ID | Group ID |
| --- | --- | --- | --- | --- |
| WT | 1 | 200 mg/kg of TAM for 3 times every other day | WT 001 | (-) controls |
| mTmG | 1 | 200 mg/kg of TAM for 3 times every other day | mTmG 001 | (-) controls |
| *Sftpc*<sup>CreERT2</sup>:mTmG | 1 | 100 mg/kg of TAM for 2 times every other day | S:TG 001 | (-) controls |
| *Gramd2*<sup>CreERT2</sup>:mTmG | 1 | 200 mg/kg of TAM for 3 times every other day | G:TG 001 | (-) controls |
| *Sftpc*<sup>CreERT2</sup>:KRAS<sup>G12D</sup>:mTmG | 2 | 100 mg/kg of corn oil for 2 times every other day | S:K:TG 003, 009 | Spon. *Sftpc*<sup>+</sup> KRAS<sup>G12D</sup> LUAD |
| *Gramd2*<sup>CreERT2</sup>:KRAS<sup>G12D</sup>:mTmG | 2 | 200 mg/kg of corn oil for 3 times every other day | G:K:TG 007, 014 | Vehicle controls |
| *Sftpc*<sup>CreERT2</sup>:KRAS<sup>G12D</sup>:mTmG | 3 | 100 mg/kg of TAM for 2 times every other day | S:K:TG 005, 008, 010 | *Sftpc*<sup>+</sup> KRAS<sup>G12D</sup> LUAD |
| *Gramd2*<sup>CreERT2</sup>:KRAS<sup>G12D</sup>:mTmG | 3 | 200 mg/kg of TAM for 3 times every other day | G:K:TG 004, 005, 011 | *Gramd2*<sup>+</sup> KRAS<sup>G12D</sup> LUAD |

We observed that LUAD lesions formed in vehicle controls for *Sftpc^+^*KRAS^G12D^ LUAD. There was a high degree of overlap in the distribution of cell cluster association in the UMAP between the spontaneously formed *Sftpc^+^* KRAS^G12D^ LUAD (abbreviated as Spon*. Sftpc^+^* KRAS^G12D^ LUAD) and tamoxifen-induced LUAD. This may be due to the previously reported leaky expression of *Sftpc^CreERT2^* in AT2 cells^46^. To determine the identities of each cluster, the mouse LungMAP^47^ reference database was used to assign identity for each cell profiled to major lung lineages (level 1, Figure 1d) as well as the sub-lineages (level 2, Figure 1e). The proportion of these ten cell sub-lineages was then compared between experimental groups (Figure 1f, Supplemental Table 1); revealing that the observed proportions of alveolar epithelial cells (AECs, *p* < 2.2e-16) and myeloid cells (*p* < 2.2e-16) were significantly higher in the three LUAD groups compared to the healthy controls. Within the LUAD experimental groups, *Sftpc*^+^ KRAS^G12D^ LUAD showed significantly increased proportions of alveolar epithelial cells (*p* < 2.2e-16) as compared to *Gramd2*^+^ KRAS^G12D^ LUAD.

Further investigating this difference in myeloid populations, we utilized a previously published spatial transcriptomic (ST-seq) dataset on an independent cohort of *Sftpc^+^* KRAS^G12D^ LUAD, *Gramd2^+^* KRAS^G12D^ LUAD^13^, and controls to determine the frequency of macrophage-rich tertiary lymphoid structure (TLS), which emerge in non-lymphoid tissues under chronic inflammatory conditions, such as cancer^48^ and play a crucial role in immune surveillance by facilitating the local activation of immune responses against tumor cells^49^ with associated responsiveness to immunotherapies, such as immune checkpoint inhibitors^50–53^. TLS were calculated using a composite score expression for multiple immune cell types, including B cells: *Igha*, *Ighg1*, *Ighg3*, *Ighm*, *Igkc*, *Iglc1*, *Iglc2*, *Iglc3*, *Jchain*, *Cd79a*, *Fcrl5*, *Mzb1*, *Ssr4*, *Xbp1*; T cells: *Trbc2*, *Il7r*; for fibroblasts and stroma: *Cxcl12*, *Lum*; and for macrophages: *C1qa*, *C7*, *Cd52*, *Apoe*, *Pim2*, *Derl3*, *Timd4*, *Cd163*, and *Folr2*. Intensity mapping of composite TLS score onto each sample’s transcriptomic array highlighted high TLS signatures, indicative of immune-cell enriched pockets within the tumor landscape (Figure 1g). Significantly elevated TLS scores were observed within *Gramd2^+^* KRAS^G12D^ LUAD compared to *Sftpc^+^*KRAS^G12D^ LUAD (Figure 1h). The striking difference in TLS occurrence between tumors of different cells of origin suggested that major shifts were occurring in the TIME that merited further investigation.

### Myeloid cells show distinct differential trajectories between AT2- and AT1-derived LUAD

To fully characterize myeloid subpopulation differences between cell of origin models of LUAD, cells annotated with LungMAP myeloid sub-lineage identity were subset out from the overall dataset. Myeloid cells (n = 19,092) were then segregated into higher-resolution clusters to identify unique myeloid populations, the stability of which was validated using “clustree” (Supplemental Figure 2a). Of the 22 myleoid-lineage sub-clusters, 8 previously described cell types were identified (Figure 2a) with marker gene expression patterns consistent with multiple previously published scRNA-seq studies^47,54–58^ (Supplemental Figure 2b-d). These included: hematopoietic stem and progenitor cells (HSPCs; *Kit*/CD117^+^, *Ly6a*/SCA-1 ^+^, *Cd34* ^+^), inflammatory monocytes (iMONs; *Csf1r* ^+^, *Itgam/*CD11B ^+^, *Cx3cr1* ^+^), proliferating myeloid cells (*Neil* ^+^*, Kif14* ^+^ *, Esco2* ^+^*, Mki67* ^+^), myeloid-derived suppressor cells (MDSCs; *Fcer1g* ^+^, *Tyrobp* ^+^, *Spi1* ^+^*, Ly6c1* ^+^), tumor associated macrophages (TAMs; *Mmp8* ^+^*, Mmp12* ^+^*, Arg1* ^+^*, Chil3* ^+^*, Chil4* ^+^*, Cd274/*PD-L1 ^+^*, Cd33/*SIGLEC3 ^+^), macrophages (*Ptprc/*CD45 ^+^*, Cd33* ^+^*, Siglecf* ^+^*, Mertk* ^+^*, Itgax/*CD11C) ^+^, dendritic cells (DCs;*H2-Aa* ^+^*, H2-Ab1* ^+^*, H2-Eb1* ^+^*, Cst3* ^+^) and granulocytes (*Csf3r* ^+^*, Cd24a* ^+^*, Trem1* ^+^*, Ltf* ^+^) (Figure 2b). While 8 previously known myeloid cell identities were observed in all five experimental groups, the proportion of these cells showed significant differences in occurrence, especially between *Sftpc^+^*KRAS^G12D^ LUAD and *Gramd2^+^* KRAS^G12D^ LUAD (Figure 2c, Supplemental Table 2). Specifically, the enrichments of iMONs (*p* < 0.01) and macrophages (*p* < 2e-16) were significantly higher in the *Gramd2^+^* KRAS^G12D^ LUAD model, whereas the cell proportions of MDSCs (*p* < 2e-16) and TAMs (*p* < 2e-16) were significantly higher in *Sftpc^+^* KRAS^G12D^ LUAD (Figure 2d, Supplemental Table 3). These findings were independently replicated in the previously published ST-seq dataset (Supplemental Figure 3).

**Fig. 2:**
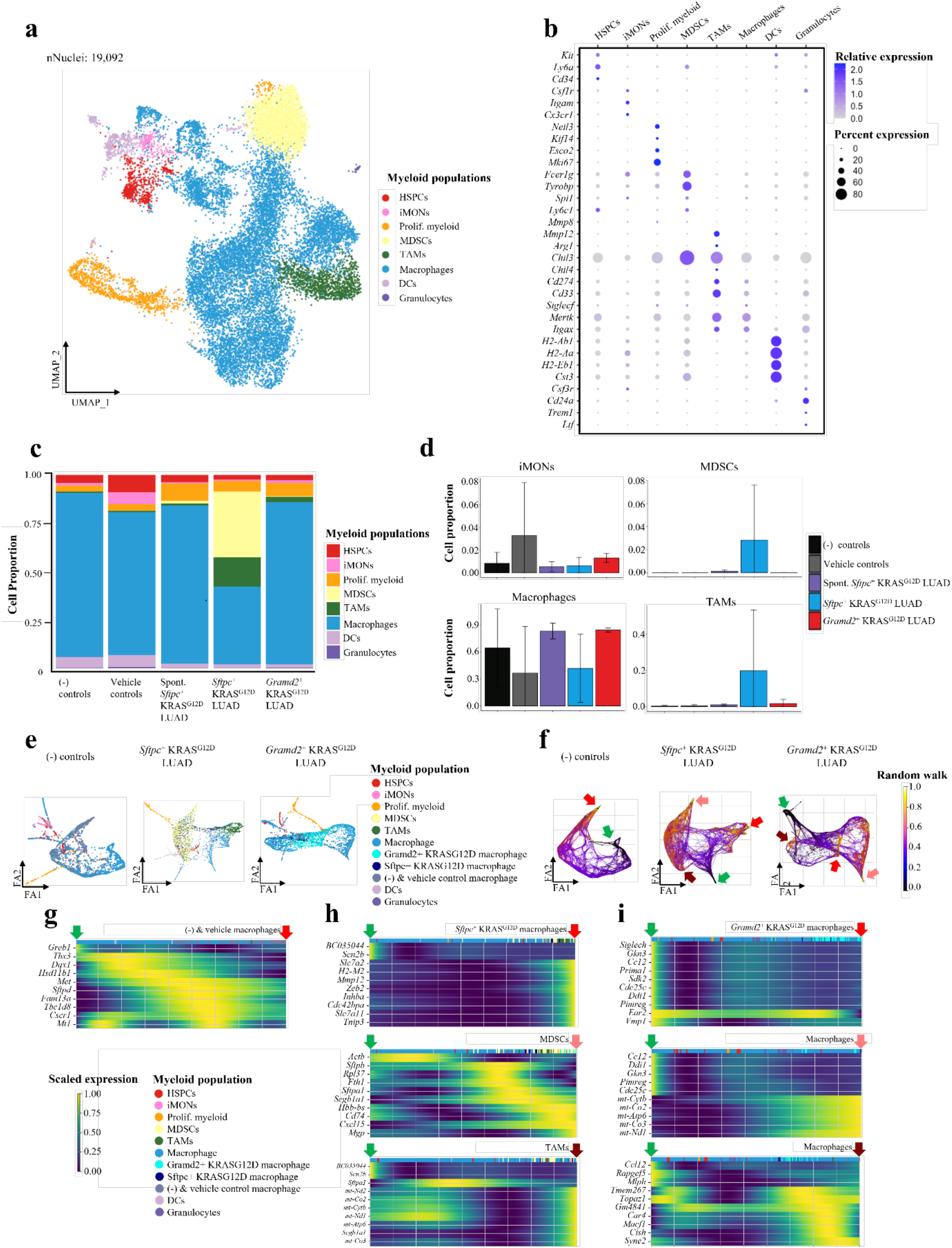
Myeloid cells show distinct differential trajectories between AT2- and AT1-derived LUAD. a, UMAP plot of LungMAP-annotated myeloid sub-lineage cells. Cells colored by myeloid cell-state identity. Red = Hematopoietic stem and progenitor cells (HSPCs), orange = inflammatory monocytes (iMONs) pink= proliferating myeloid cells (prolif. myeloid cells), yellow = myeloid-derived suppressor cells (MDSCs), green = tumor-associated macrophages (TAMs), blue = alveolar macrophages (macrophages), lavender = dendritic cells (DCs), and purple = granulocytes. b, Dot plot of gene expression of marker genes for myeloid annotation. Dot colors were based on normalized data and used to indicate the relative expression levels, blue = high expression, grey = low expression. Dot sizes indicate the percentage of cells expressing the indicated gene in the subpopulation. c, Bargraph indicating the proportion of myeloid cell subpopulations within each sample group. Cells colored as in (a). d, The average proportion of different types of myeloid cells within each sample group. Error bars (standard deviation) are shown in Barplots. e, Force-directed graph layout of cells from indicated sample groups. Cells are color coded as in (a) with the additional splitting of ‘macrophage’ populations by experimental type, cyan = *Gramd2^+^* KRAS^G12D^ macrophages, navy blue = *Sftpc^+^* KRAS^G12D^ macrophages, charcoal = (-) control and vehicle control macrophages. f, Simulated random walk of the diffusion psuedotime transition matrix, project on the force-directed graph layout. Random walk start point is denoted with a green arrow,and end points are labeled with red arrows, with different shades corresponding to different terminal points. Line color indicates transition from starting population (black) to terminal population (yellow). g, Heatmap of temporal gene activation within specific lineages present in the negative (-) and vehicle control macrophage group. Cells are sorted by diffusion pseudotime value. Green arrow = computed starting point for indicated macrophage lineage from (f), shaded red arrows = terminal end points from (f). Myeloid subpopulation classification as in (e). Heatmap colors are scaled for gene enrichment, blue = low expression, yellow = high expression. h, Heatmap of temporal gene activation within specific lineages present in the *Sftpc^+^* KRAS^G12D^ macrophagessample group. Cells are sorted by diffusion pseudotime value. Arrows correspond to terminal end points from (f). Myeloid subpopulation classification as in (e). Heatmap colors are scaled for gene enrichment, blue = low expression, yellow = high expression. i, Heatmap of temporal gene activation within specific lineages present in the *Gramd2^+^* KRAS^G12D^ macrophagessample group. Cells are sorted by diffusion pseudotime value. Arrows correspond to terminal end point from (f). Myeloid subpopulation classification as in (e). Heatmap colors are scaled for gene enrichment, blue = low expression, yellow = high expression.

To further study the dynamics of myeloid differentiation in response to LUAD, we performed trajectory analysis on the entire myeloid lineage population within each experimental condition and re-computed the nearest neighbors distance matrix. To visualize the global structure of macrophage cellular relationships and how they progress between different phases force-directed graph drawing in ForceAtlas2 was performed (Figure 2e). To identify dynamic gene changes within unique populations of myeloid cells, diffusion pseudotime was applied to the common macrophage subclusters present between experimental groups. Random walk projection using CellRank2 identified the *Cd33^+^* population of macrophages as the starting point, shown in black, as well as the projected end points, shown in yellow (Figure 2f). The low threshold connectivity branches from the partition-based graph abstraction (PAGA) plot were removed from pseudotime analysis to focus on the main myeloid lineages (Supplemental Figure 4). Genes with the highest correlation towards a terminal macrostate were identified with CellRank’s estimator model (Figure 2g-i). Only one terminal macrostate was observed in the negative and vehicle controls (Figure 2g), whereas three unique terminal macrostates of myeloid cells were identified in both the *Sftpc^+^* KRAS^G12D^ LUAD (Figure 2h) and *Gramd2^+^* KRAS^G12D^ LUAD experimental groups (Figure 2i). In the *Sftpc^+^* KRAS^G12D^ LUAD model, myeloid terminal macrostates included: lung-resident macrophages, MDSCs, and immunosuppressive TAMs. In the *Gramd2^+^* KRAS^G12D^ LUAD model, all macrophage lineages were classified as types of differentiated macrophages that varied in their activation state for metabolic and/or inflammatory-associated genes.

Interestingly, the *Gramd2^+^* KRAS^G12D^ LUAD contained one unique lineage of macrophages with altered expression of mitochondrial genes such as *mt-Cytb*, *mt-Co2*, and *mt-Atp6* (Figure 2i). Previous work has shown that enhanced mitochondrial metabolism corresponds to macrophage activation^59,60^, suggesting immunoreactive macrophages are present in *Gramd2^+^* KRAS^G12D^ LUAD. In contrast, the *Sftpc^+^*KRAS^G12D^ LUAD model contained two distinct tumor associated macrophage lineages as well as a MDSC lineage (Figure 2h). Taken together, *Gramd2^+^*KRAS^G12D^ LUAD and *Sftpc^+^* KRAS^G12D^ LUAD exhibit transcriptionally distinct myeloid cell lineages which may result in distinct macrophage subpopulations within their TIME.

### Immunoreactive macrophages are enriched within AT1-LUAD

To further understand the unique myeloid lineages present within each cell-of-origin model (Figure 3a), and how they may influence TIME states within LUAD, Nebulosa was used to establish an enrichment score for immunoreactivity (Figure 3b) based on kernel density estimation of combined *Cd86*, *Il1a*, *Cxcl2*, *Ccl4*, and *Tnf* expression^61^. In addition, a joint macrophage reactivity score including a weighted combination of 16 known markers for immunoreactive macrophages, cytokines, and chemokines, including: *Csf2, Isg15, Il1a, Il1b, Il6, Il8, Il22, Il23a, Cxcl3, Cxcl9, Cxcl10, Ccl4, Cd80, Cd86, Il2,* and *Il18* was used to determine immunoreactive state^62–70^ across all experimental groups (Figure 3c). *Gramd2^+^* AT1 KRAS^G12D^ LUAD samples exhibited statistically significantly higher enrichment for immunoreactive/anti-tumoral states of macrophages (*p* value < 2e^-16^) than *Sftpc^+^* AT2 KRAS^G12D^ LUAD. Immunoreactive macrophages were enriched in the *Gramd2^+^* AT1 KRAS^G12D^ LUAD samples relative to both vehicle controls and *Sftpc^+^* AT2 KRAS^G12D^ LUAD samples (Figure 3c). Validation of expression levels for the immunoreactive macrophage gene signature in Figure 3b was also performed on the previously published ST-seq dataset^13^ between *Sftpc^+^* AT2 KRAS^G12D^ LUAD and *Gramd2^+^* AT1 KRAS^G12D^ LUAD sections. The immunoreactive score was found to be statistically significantly elevated in *Gramd2^+^* AT1 KRAS^G12D^ LUAD (*p* value < 1.4e^-14^) compared to *Sftpc^+^* AT2 KRAS^G12D^ LUAD (Figure 3d).

**Fig. 3:**
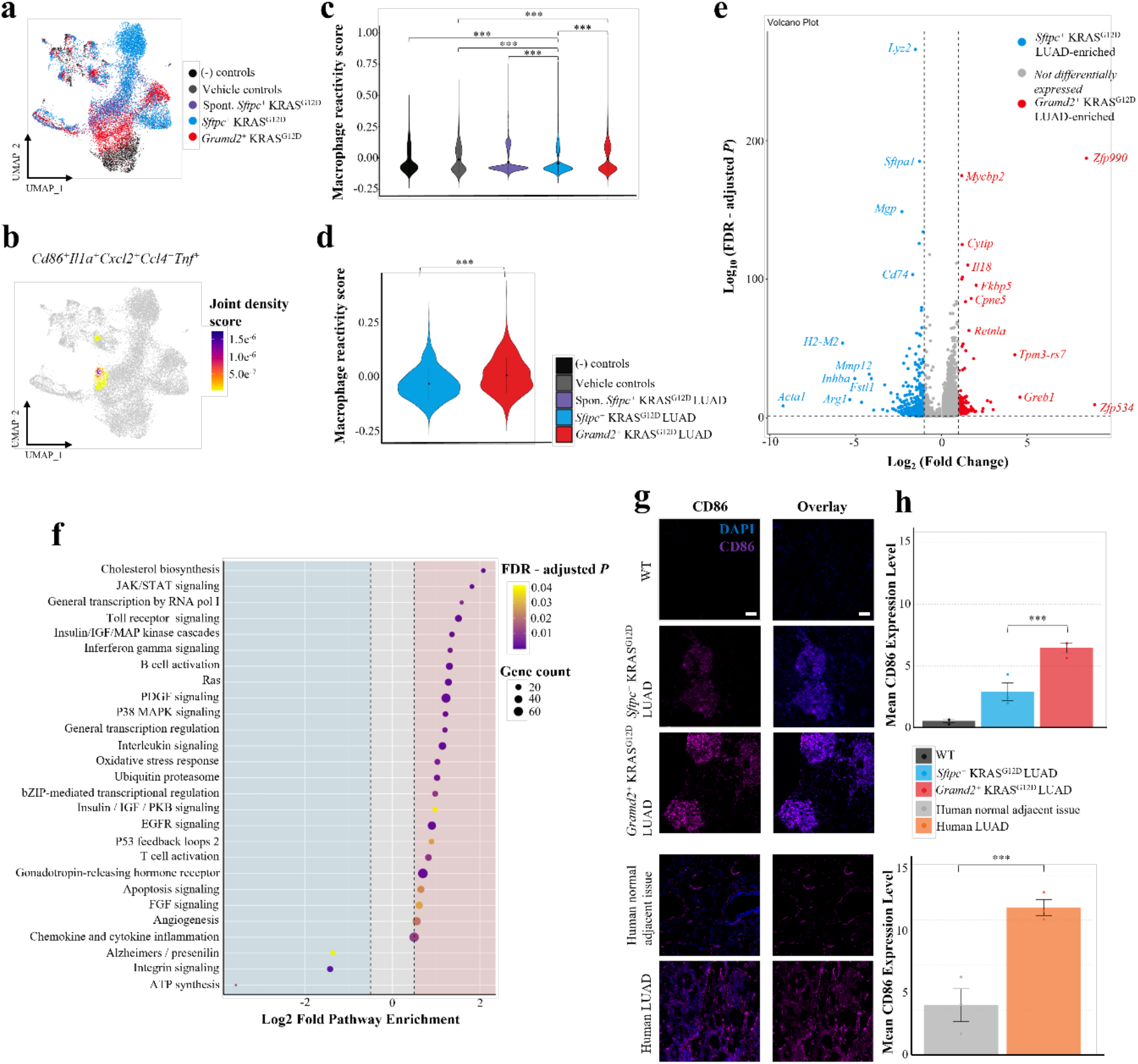
Immunoreactive macrophages are enriched within AT1-LUAD. a, UMAP plot of LungMAP-annotated myeloid sub-lineage cells. Cells colored by experimental group, black = negative (-) controls, grey = vehicle controls, purple = spontaneous *Sftpc^+^* KRAS^G12D^ LUAD, blue = *Sftpc^+^* KRAS^G12D^ LUAD, and red = *Gramd2^+^* KRAS^G12D^ LUAD. b, UMAP of Nebulosa joint kernel density plot highlighting known immune-active macrophage features (*Cd86^+^Il1a^+^Cxcl2^+^Ccl4^+^Tnf^+^).* Color indicates joint density score, low = grey, high = purple. c, Violin plot of macrophage reactivity score between experimental groups. Colors as in (a). Macrophage reactivity score was measured based on joint expression of known immuno-active gene features: *Csf2, Isg15, Il1a, Il1b, Il6, Il8, Il22, Il23a, Cxcl3, Cxcl9, Cxcl10, Ccl4, Cd80, Cd86, Il2, Il18.* A Kruskal Wallis test was used to test the significance of difference of mean macrophage reactivity score overall (*p* value < 2.2e^-16^), and a pairwise Wilcoxon test was used to determine significance of differences between individual experimental groups, *p* value < 2e^-16^). d, Violin plot of macrophage reactivity score within previously published ST-seq dataset^13^, using the joint expression of the same genes as (c). Colors as in (a). Wilcoxon rank-sum test was applied to test the significance of differences of mean macrophage reactivity score between *Sftpc^+^* KRAS^G12D^ LUAD and *Gramd2^+^*KRAS^G12D^ LUAD. P value < 1.4e^-14^). e, Volcano plot of differentially expressed genes (DEGs) in cells assigned “macrophage” identity within the myleoid lineage as calculated between *Gramd2^+^* KRAS^G12D^ LUAD and *Sftpc^+^* KRAS^G12D^ LUAD samples. X-axis = log2 fold change difference in gene expression between *Gramd2^+^* KRAS^G12D^ LUAD macrophage (+) and *Sftpc^+^* KRAS^G12D^ LUAD (-) macrophage population. Y-axis = significant expression differences were adjusted through false discovery rate (FDR) correction, adjusted p value cutoff = 0.05; Log2FC cutoff = 1. Red = genes enriched in the *Gramd2^+^* KRAS^G12D^ LUAD macrophage population relative to *Sftpc^+^* KRAS^G12D^ LUAD macrophages. Blue = genes enriched in *Sftpc^+^* KRAS^G12D^ LUAD macrophages relative to the *Gramd2^+^* KRAS^G12D^ LUAD macrophage population. f, Bubble plot of significant PANTHER pathways predicted based on the differentially expressed genes of macrophages between *Gramd2^+^* KRAS^G12D^ LUAD macrophages and *Sftpc^+^* KRAS^G12D^ LUAD macrophages. X-axis = log2 fold enrichment values for *Gramd2^+^*KRAS^G12D^ LUAD macrophages (+) as compared to *Sftpc^+^*KRAS^G12D^ LUAD macrophages (-). Dot color = FDR-corrected p value for pathway enrichment in PANTHER, purple = highly significant, yellow = approaching threshold of significance cut-off (0.05). Dot size = gene counts per PANTHER pathway. g, Representative immunofluorescent staining images of CD86 in mouse and human FFPE tissue sections. Top = Mouse FFPE lung sections with indicated genotypes, bottom = human LUAD tissue microarray (TMA) sections. Purple = CD86, blue = DAPI + CD86 merged images. Scale bar = 100 μm. h, Quantification of CD86 expression in (g). Mean intensity values within experimental groups are shown. Error bars represent standard deviation between genotype biological replicates. An unpaired T test was used to assess significance of observed differences between indicated groups. p < 0.05 = *, p < 0.01 = **, p < 0.001 = ***.

To understand more broadly the transcriptomic differences in macrophage populations that varied between tumors with different alveolar epithelial cell of origin, differentially expressed genes (DEGs) between *Sftpc^+^* AT2 KRAS^G12D^ LUAD and *Gramd2^+^* AT1 KRAS^G12D^ LUAD models were calculated using the FindMarkers function in Seurat (Figure 3e, Supplemental Table 4). A total of 5481 significant gene expression alterations were detected within macrophages in *Sftpc^+^* AT2 KRAS^G12D^ LUAD compared to *Gramd2^+^* AT1 KRAS^G12D^ LUAD. Of note, zinc finger protein 990 (*Zfp990*, fold change = 8.46, FDR-corrected p-value = 4.78e^-188^), which is involved in macrophage activation and inflammatory processes^71^, showed significantly higher expression in *Gramd2^+^* AT1 KRAS^G12D^ LUAD macrophages. Interleukin 18 (*Il18*, fold change = 1.54, FDR-p value = 5.92 e^-111^), FKBP prolyl isomerase 5 (*Fkbp5*, fold change = 2.03, FDR-p value = 2.5e^-96^), and MYC Binding Protein 2 (*Mycbp2*, fold change 1.19, FDR-p value 1.56e^-175^), all upregulated in the macrophage population of *Gramd2^+^* AT1 KRAS^G12D^ LUAD, are implicated in recruiting various immune cells, activating pro-inflammatory pathways and thus facilitating immune response^72–74^. In contrast, genes elevated in *Sftpc^+^* AT2 KRAS^G12D^ LUAD macrophages included Arginase 1 (*Arg1*, fold change = -5.38, FDR p value = 8.48e^-14^) which is associated with immunosuppressive activities in tumor-associated macrophages and MDSCs. *Sftpc^+^* AT2 KRAS^G12D^ LUAD specific expression of Matrix metallopeptidase 12 (*Mmp12*, fold change = 4.20, FDR p value = 3.35e^-32^) whose role in extracellular matrix degradation is well established^75^, was also observed. Several genes known to be associated with alveolar identity were also identified, including surfactant protein A1 (*Sftpa1)* in *Sftpc^+^*KRAS^G12D^ LUAD macrophages, which has been observed previously ^76^.

To determine the functional impact these DEGs within macrophages may have on anti-cancer immunity, we performed gene set enrichment analysis (GSEA) by leveraging Protein ANalysis THrough Evolutionary Relationships (PANTHER)^77,78^. Pathways significantly enriched in *Gramd2^+^* AT1 KRAS^G12D^ LUAD included interferon-gamma signaling (fold enrichment = 1.32, FDR p-value = 2.4e^-3^), interleukin signaling (fold enrichment = 1.14, FDR p-value = 2.8e^-6^), oxidative stress response (fold enrichment = 1.02, FDR p-value = 4.4e^-3^), and inflammation signaling (fold enrichment = 0.49, FDR p-value = 1.1e^-2^), suggesting that macrophages in *Gramd2^+^* AT1 KRAS^G12D^ LUAD model possess a more active, inflamed immune state than macrophages present in *Sftpc^+^* AT2 KRAS^G12D^ LUAD (Figure 3f, Supplemental Table 5). These findings were further supported using fgsea, an R-package for fast pre-ranked GSEA^79^. In total, ninety (90) significantly enriched pathways were detected in macrophages between *Gramd2^+^* AT1 KRAS^G12D^ LUAD and *Sftpc^+^* AT2 KRAS^G12D^ LUAD (Supplemental Figure 5).

To validate enrichment for immunoreactive macrophages in the *Gramd2^+^* AT1 KRAS^G12D^ LUAD model, we performed immunofluorescence staining on FFPE-embedded lung sections from all sample groups using cluster of differentiation antigen 86 (*Cd86*), a cell surface marker of inflammation on antigen-presenting cells, including macrophages^80,81^ (Figure 3g). CD86 expression was significantly elevated in both *Gramd2^+^* AT1 KRAS^G12D^ LUAD and *Sftpc^+^* AT1 KRAS^G12D^ LUAD models relative to controls (ANOVA p value 1.36e^-3^); however CD86 levels were higher in the *Gramd2^+^* AT1 KRAS^G12D^ LUAD model (p value < 1.24e^-2^) (Figure 3h). To determine if these results in mouse models bear relevance to human LUAD, we performed CD86 staining on twenty-six (26) tumor microarray (TMA) sections derived from 9 individual LUAD samples of primary human FFPE LUAD, which also displayed significantly elevated CD86 levels (Figure 3h). Taken together, these results indicate that macrophages in *Gramd2^+^*AT1 KRAS^G12D^ LUAD adopt an immunoreactive, anti-tumoral state.

### Immunosuppressive myeloid cells are enriched within AT2-LUAD

While *Gramd2^+^* AT1 KRAS^G12D^ LUAD macrophages exhibited enrichment for immuno-active, proinflammatory macrophages, *Sftpc^+^* AT2 KRAS^G12D^ LUAD myeloid cells showed an increased expression level of *Arg1*, which is associated with an immunosuppressive TIME^82–84^. To characterize the types of immunosuppressive myeloid-lineage cells present in the TIME of the *Sftpc^+^* AT2 KRAS^G12D^ LUAD model (Figure 4a), the enrichment of known *Arg1^+^* myeloid populations (TAMs; *Msr1^+^Mrc1^+^Chil3^+^Arg1^+^Cd274*^+^, and MDSCs; *S100a8^+^S100a9^+^Wfdc17^+^Cd36^+^Cd38^+^Cd84^+^*) was plotted within the larger myeloid lineage cell population (Figure 4b). Enrichment for TAMs and MDSCs overlapped with clusters composed primarily of myeloid cells derived from the *Sftpc^+^* AT2 KRAS^G12D^ LUAD samples (Figures 4a-b). A panel of nine genes with known immunosuppressive activity (*Slc7a2*, *Slc7a11*, *Mmp12*, *Arg1*, *Msr1*, *Inhba*, *H2-M2*, *Chil4*, and *Mafb*)^82,85–94^ was also examined for their distribution within the myeloid population amongst experimental groups. All immunosuppressive markers showed statistically significant elevation in the *Sftpc^+^* AT2 KRAS^G12D^ LUAD myeloid cells relative to the *Gramd2^+^* AT1 KRAS^G12D^ LUAD myeloid population (Figure 4c). A joint macrophage suppression score including a weighted combination of 28 known markers for TAMs, cytokines, and chemokines, including: *CD163*, *Spp1*, *Apoe*, *Gpnmb*, *Trem2*, *Il10*, *Il13*, *Il33*, *Vegf*, *Csf1*, *Ccl2*, *Ccl5*, *Ccl17*, *Ccl22*, *Tgfb*, *Rgf*, *Pdgf*, *Arg1*, *Fibrin*, *Wnt*, *MMPs*, *Vcan*, *Cxcl8*, *Csf1*, *Csf3*, *Cd47*, *Enpp1*, *Cgas*, and *Sting*^62–64,66–70,95–98^ was then used to determine immunosuppression state across all experimental groups (Figure 4d). *Sftpc^+^* AT2 KRAS^G12D^ LUAD samples were again observed to be statistically significantly enriched for immunosuppressive myeloid cell states (*p* value < 2e-16).

**Fig. 4:**
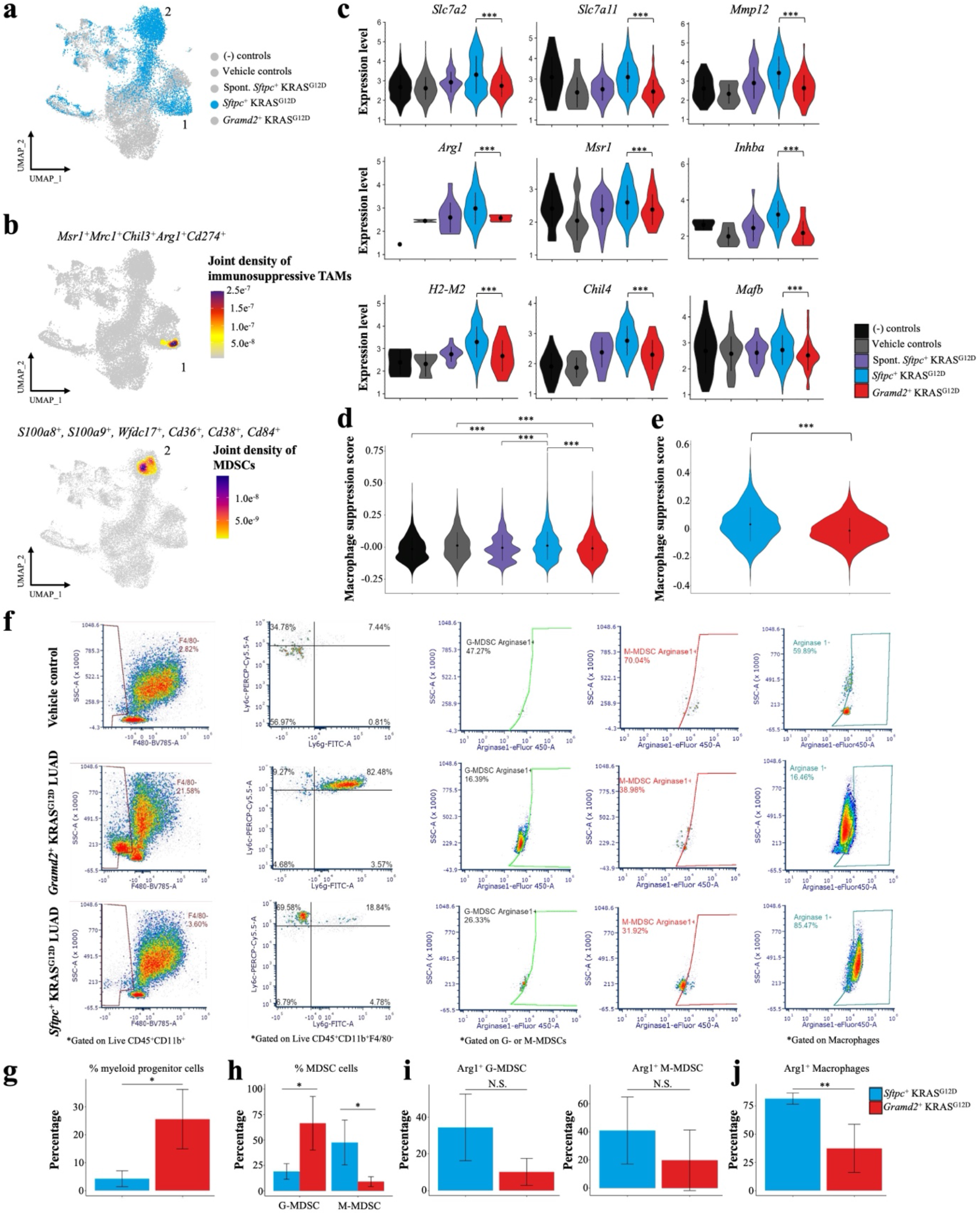
Immunosuppressive myeloid cells are enriched within AT2-LUAD. a, UMAP plot of LungMAP-annotated myeloid sub-lineage cells. Blue = *Sftpc^+^* KRAS^G12D^ LUAD myleoid-lineage cells, grey = myleoid-lineage cells from all other experimental groups. b, UMAP of Nebulosa joint kernel density plot highlighting known gene signatures of immunosuppressive TAMs(*Msr1^+^Mrc1^+^Chil3^+^Arg1^+^Cd274*^+^) and MDSCs (*S100a8^+^, S100a9^+^, Wfdc17^+^, Cd36^+^, Cd38^+^, Cd84^+^*). Color indicates joint density score, low = grey, red = purple c, Violin plots of expression levels for the indicated known immunosuppressive myeloid cell markers within each experimental group based on snRNAseq dataset. Black = negative (-) controls, grey = vehicle controls, purple = spontaneous *Sftpc^+^*KRAS^G12D^ LUAD, blue = *Sftpc^+^* KRAS^G12D^ LUAD, and red = *Gramd2^+^* KRAS^G12D^ LUAD. A Kruskal Wallis test was used to test the significance of the difference of mean macrophage reactivity score overall (*p* value < 2.2e-16), and a pairwise Wilcoxon test was used to determine significance of differences between individual experimental groups, *p* value < 2e-^16^). *p < 0.05, **p < 0.01, ***p < 0.001. d, Violin plot of aggregate myeloid suppression score across all experimental groups based on the previously published ST-seq dataset^13^. Experimental groups are colored as in (c). A Kruskal-Wallis test was applied to test the significance of mean macrophage suppression score across five groups (*p* value < 2.2e^-16^). A Wilcoxon rank-sum test was applied to test the significance of differences in mean macrophage suppression score between *Sftpc^+^*KRAS^G12D^ LUAD and *Gramd2^+^* KRAS^G12D^ LUAD. *p* value < 2e^-16^). *p < 0.05, **p < 0.01, ***p < 0.001. e, Violin plot of macrophage suppression score from (d) as measured within the previously published ST-seq dataset between *Sftpc^+^* KRAS^G12D^ LUAD and *Gramd2^+^*KRAS^G12D^ LUAD samples. Wilcoxon rank-sum test was applied to test the significance of observed differences (p value < 4.8e^-15^). f, Representative flow cytometric plots: Four panels from left to right are gated on: expression of F4/80 on CD45+CD11b+ myeloid cells to identify myeloid progenitor cells within mouse lung tissue; expression of LY6G and LY6C on CD45^+^CD11b^+^F4/80^-^ for MDSC classification; ARG1 positivity on CD45^+^CD11b^+^F4/80^-^Ly6G^+^Ly6C^low^ G-MDSCs and CD45^+^CD11b^+^F4/80^-^Ly6G^-^Ly6C^high^ M-MDSCs; and ARG1 positivity on CD45^+^CD11b^+^Ly6g^-^F4/80^+^MHCII^+^ macrophages. g, Bar plot of myeloid precursor cell percent occurrence within each LUAD cell-of-origin model. Y-axis. = percentage of myeloid precursor cells within each tamoxifen-induced experimental group. Color of bars as in (e). Error bars represent means with standard deviation. (N = 3). An unpaired two-tailed Student’s t-test was used to determine significance. *p < 0.05, **p < 0.01, ***p < 0.001. h, Bar plot of MDSC cell percent occurrence within each LUAD cell-of-origin model. Y-axis. = percentage of MDSCs within each tamoxifen-induced experimental group. Color of bars as in (e). Error bars represent means with standard deviation. (N = 3). An unpaired two-tailed Student’s t-test was used to determine significance. *p < 0.05, **p < 0.01, ***p < 0.001. i, Bar plots of Arg1^+^ G-MDSC (left) and Arg1^+^ M-MDSC (right) occurrence within each LUAD cell-of-origin model. Y-axis = percentage of Arg1^+^ MDSC subpopulation within each tamoxifen-induced experimental group. Color of bars as in (e). Error bars represent means with standard deviation. (N = 3). An unpaired two-tailed Student’s t-test was used to determine significance. *p < 0.05, **p < 0.01, ***p < 0.001. j, Bar plot of Arg1^+^ macrophage cell percent occurrence within each LUAD cell-of-origin model. Y-axis. = percentage of Arg1^+^ macrophage within each tamoxifen-induced experimental group. Color of bars as in (e). Error bars represent means with standard deviation. (N = 3). An unpaired two-tailed Student’s t-test was used to determine significance. *p < 0.05, **p < 0.01, ***p < 0.001.

Validation of the observed immunosuppressive state enrichment for myeloid-lineage cells in the *Sftpc^+^* AT2 KRAS^G12D^ LUAD model was then performed on an independent, previously published ST-seq profiling dataset of identical genotypes and treatment course^13^. The same joint immunosuppressive score testing used in the snRNAseq dataset was applied to the ST-seq dataset (Figure 4e) and was used to determine immunosuppression state across each spot within all KRAS^G12D^ LUAD samples. The *Sftpc^+^* KRAS^G12D^ LUAD samples exhibited a significantly higher joint immunosuppressive score than that present in the *Gramd2^+^* KRAS^G12D^ LUAD model (*p* value < 4.8e^-15^), indicating that the TIME macrophage populations in AT2-derived LUAD samples were dominated by an anti-tumor signature (Figure 4e). Taken together, both snRNA-Seq and ST-seq analysis on an independent cohort of samples suggest that *Sftpc^+^* KRAS^G12D^ LUAD is enriched for immunosuppressive TAMs and MDSCs compared to the *Gramd2^+^*KRAS^G12D^ LUAD TIME.

Further validation of immunosuppressive myeloid population enrichment in the TIME of *Sftpc^+^* KRAS^G12D^ LUAD was performed through flow cytometry on single cell suspensions prepared from dissociated whole mouse lung tissue (Figure 4f). Macrophages and MDSCs were identified in tamoxifen-induced *Sftpc^+^* KRAS^G12D^ LUAD, *Gramd2^+^* KRAS^G12D^ LUAD, and corn oil vehicle controls from matching genotypes by gating on live CD45^+^CD11b^+^F4/80^+^MHCII^+^ and live CD45^+^CD11b^+^F4/80^-^ cells, respectively. MDSCs were further classified into granulocytic (G-MDSC) and monocytic (M-MDSC) subtypes by splitting the general MDSC population into Ly6C^high^Ly6G^-^ M-MDSC and Ly6G^+^Ly6C^low^ G-MDSC subpopulations. A statistically significant difference in the percentage of myeloid progenitor cells (live CD45^+^CD11b^+^F4/80^-^) was observed between the *Gramd2^+^* KRAS^G12D^ LUAD and *Sftpc^+^* KRAS^G12D^ LUAD models (*p* value < 0.05), with the *Gramd2^+^* KRAS^G12D^ LUAD showing elevated myeloid progenitor cell levels (Figure 4g). Within the MDSC population, a significant difference in the percentage of Ly6c^+^Ly6g^-^ M-MDSC cells and G-MDSC subpopulation was observed between the *Gramd2^+^* KRAS^G12D^ LUAD and *Sftpc^+^* KRAS^G12D^ (Figure 4h, *p* value < 0.05), with M-MDSC elevated in *Sftpc^+^* KRAS^G12D^ LUAD and G-MDSC elevated in *Gramd2^+^* KRAS^G12D^ LUAD. Testing Arg1 positivity of both the M-MDSCs and G-MDSCs populations showed that there was a trend toward a higher mean percentage of Arg1^+^ MDSCs in the *Sftpc^+^*KRAS^G12D^ LUAD model for both G- and M-subpopulations, when compared to *Gramd2^+^* KRAS^G12D^ LUAD, though the relatively small sample size (N=3) resulted in a lack of statistical significance (Figure 4i). However, Arg1^+^ cells among all macrophage populations were significantly elevated in *Sftpc^+^* KRAS^G12D^ LUAD relative to *Gramd2^+^* KRAS^G12D^ LUAD (Figure 4j, *p* value < 0.01). Taken together, flow cytometry analysis confirmed that elevated Arg1^+^ MDSC and TAM populations were increased in the *Sftpc^+^* KRAS^G12D^ LUAD compared to the *Gramd2^+^* KRAS^G12D^ LUAD model.

### AT1- and AT2-derived LUADs exhibit heterogenous cell-cell communication patterns

To unravel how cell of origin could influence epithelial-myleoid interactions to foster distinctly different immunomodulatory states within the TIME, LIANA^99^ was used to integrate cell-cell communication (CCC) interaction prediction between malignant LUAD epithelial and myeloid-lineage cell populations. First, malignant AECs that from both *Sftpc^+^* KRAS^G12D^ LUAD and *Gramd2^+^* KRAS^G12D^ LUAD experimental groups were identified using InferCNV^100^, with a cutoff of greater than 95 percentile (>95%) copy number variation (CNV) score relative to control cells to screen out non-KRASG12D recombined AEC. Second, cells annotated within the myeloid lineage from both *Sftpc^+^* KRAS^G12D^ LUAD and *Gramd2^+^* KRAS^G12D^ LUAD experimental groups were subset from the main dataset and re-clustered with the InferCNV-defined malignant epithelial cells. CCC interaction calculation was then performed between the malignant epithelial and myeloid cell populations within the two LUAD cell-of-origin models.

Frequency of interaction was compared between the distinct malignant epithelial and myeloid cell populations in both the *Sftpc^+^*KRAS^G12D^ LUAD and *Gramd2^+^* KRAS^G12D^ LUAD experimental groups (Figure 5a), with enrichment for interactions higher in the *Sftpc^+^* KRAS^G12D^ LUAD scaled in blue, and enrichment for interactions higher in the *Gramd2^+^* KRAS^G12D^ LUAD scaled in red. When comparing CCC interaction between malignant AECs as the ligand/source cells and myeloid populations as the receptor/target cells, the *Gramd2^+^* KRAS^G12D^ LUAD model had elevated levels of interaction between AT1 cells and granulocytes, macrophages, iMONs, and interestingly, AT2 cells. In contrast, the *Sftpc^+^* KRAS^G12D^ LUAD model had elevated numbers of interactions between AT2 cells and TAMs. Examining the reverse relationship, where CCC interactions originated in myeloid populations (ligand/source) to malignant AECs (target/receptor), *Gramd2^+^* KRAS^G12D^ LUAD showed increased levels of interaction from iMONs and macrophages towards AT1 cells; however, within *Sftpc^+^*KRAS^G12D^ LUAD model, although the TAMs had lower interaction activities towards AT2 cells, interactions were increased AT1 cells and AT1/AT2 cells. Interestingly, and in line with other recent studies showing transition of AT1 and/or AT2 cells into an intermediate AT1/AT2 transitional cell state^101,102^, *Sftpc^+^* KRAS^G12D^ LUAD was enriched for higher interactions between AT1/AT2 transitional cells and TAMs, regardless of which cell acted as the ligand/source cell or receptor/target cell.

**Fig. 5.**
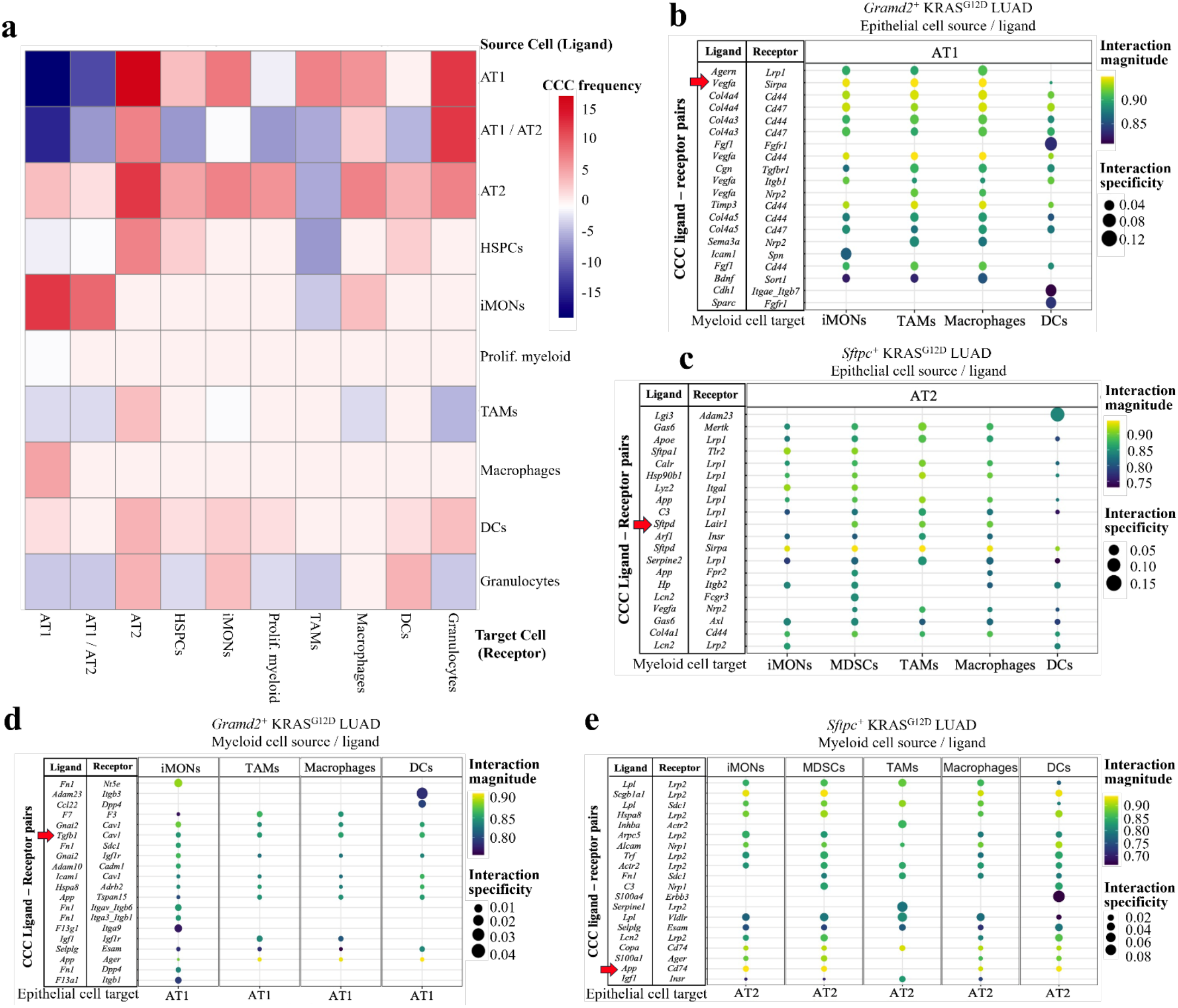
AT1- and AT2-derived LUADs exhibit heterogenous cell-cell communication patterns. a, Correlation matrix displaying enrichment for cell-cell communication (CCC) frequency of ligand receptor-pairs between distinct cell-of-origin LUAD models. Enrichment for CCC interactions in *Gramd2^+^* KRAS^G12D^ LUAD over *Sftpc^+^* KRAS^G12D^ LUAD is shown in increasingly dark shades of red, enrichment for CCC interactions in *Sftpc^+^* KRAS^G12D^ LUAD relative to those in *Gramd2^+^* KRAS^G12D^ LUAD is shown in increasingly dark shades of blue. Rows = cells predicted by LIANA^99^ to be the source of the ligand in the ligand-receptor pairs. Columns = cells predicted by LIANA^99^ to be the target and possess the receptor in the ligand-receptor pairs predicted. b, Dot plot of ligand receptor-pairs predicted within the *Gramd2^+^* KRAS^G12D^ LUAD model between malignant AT1 cells (source / ligand) and myeloid-lineage cell populations (target / receptor). Color of dot = interaction magnitude, highly expressed ligand-receptor pairs = yellow, low expression ligand-receptor pairs = blue. Size of dot = interaction specificity, highly specific ligand-receptor interaction for the indicated cell types = large dot, non-specific interactions for the indicated cell types = small dot. c, Dot plot of ligand receptor-pairs predicted within the *Sftpc^+^* KRAS^G12D^ LUAD model between malignant AT2 cells (source / ligand) and myeloid-lineage cell populations (target / receptor). Color and size of dots as in (b). d, Dot plot of ligand receptor-pairs predicted within the *Gramd2^+^* KRAS^G12D^ LUAD model between myeloid-lineage cell populations (source / ligand) and malignant AT1 cells. Color and size of dots as in (b). e, Dot plot of ligand receptor-pairs predicted within the *Sftpc^+^* KRAS^G12D^ LUAD model between myeloid-lineage cell populations (source / ligand) and malignant AT2 cells. Color and size of dots as in (b).

To understand the nature of the molecular interactions occurring between malignant epithelial and myeloid-lineage populations within our cell of origin LUAD models, the top 20 significantly enriched ligand-receptor pairs for each type of cell-cell communication were extracted from the larger interaction matrix generated by LIANA. An interesting pattern emerged, in that known cell-type specific markers for distinct AEC populations were among the top predicted ligand-receptor interaction pairs between cell types. Specifically, within the *Gramd2^+^* KRAS^G12D^ LUAD model, malignant AT1 cells were predicted to associate with iMONs, anti-tumoral TAMs, and macrophages by secreting vascular endothelial growth factor alpha (VEGFA), which has been previously reported as enriched within the AT1 cell population^103^, and interacting with signal regulatory protein alpha (SIRPA) on these specific myeloid-lineage cell states (Figure 5b, red arrow). In contrast, malignant AT2 cells within the *Sftpc^+^* KRAS^G12D^ LUAD model were predicted to interact with MDSCs, pro-tumoral TAMs, and macrophages by releasing surfactant protein-D (SFTPD), a crucial component of the pulmonary surfactant system, primarily synthesized and secreted by AT2 cells, to interact with leukocyte-associated immunoglobulin-like receptor 1 (LAIR1) on these immunosuppressive myeloid cell surfaces (Figure 5c, red arrow).

Predicted crosstalk between myeloid populations interacting with known cell markers on malignant epithelial cells was also observed. Of note, multiple myeloid populations (iMONs, TAMs, macrophages, and DCs) within the *Gramd2^+^* KRAS^G12D^ LUAD model were predicted to secrete transforming growth factor beta-1 (TGFϕ31), which can interact with the AT1-cell type specific marker Caveolin 1 (CAV1)^104^ (Figure 5d, red arrow). Similarly, amyloid precursor protein (APP) proteolysis on the surface of myeloid-lineage cells produces ligands that are known to interact with cell-differentiation marker 74 (CD74), which is known to be enriched on the surface of AT2 cells^105^ and are known to play a role in immunosuppression^106–108^. Taken together, these results suggest that the intrinsic cell-type specific expression of cell surface receptors and secreted ligands from unique AEC populations may contribute to establishment and maintenance of the immunosuppressive state within the TIME.

## Discussion

This study was designed to discover how cell of origin both influences and is influenced by the tumor immune microenvironment. Results from this study revealed insights into how alveolar cell of origin influences myeloid cell immune states in KRAS-driven LUAD, with AT1-derived *Gramd2^+^* KRAS^G12D^ LUAD promoting a pro-inflammatory TIME with enrichment for iMON and anti-tumoral macrophage populations in snRNAseq populations, elevated CD86 expression in FFPE lung sections, elevated immunoreactive gene signatures in previously published ST-seq analyses, and predicted specificity in cell-cell interactions fostering an inflammatory TIME. In contrast, AT2-driven *Sftpc^+^*KRAS^G12D^ LUAD fostered an immunosuppressive TIME with enrichment for MDSC, pro-tumoral TAMs, and DCs in snRNAseq populations, flow cytometry validated enrichment of MDSC and Arg1^+^ TAM macrophage populations, and predicted AT2-cell marker specific interactions with immunosuppressive pathway genes on myeloid lineage cells.

That *Sftpc^+^* AT2-derived LUADs foster an immunosuppressive TIME with elevated Arg1^+^ MDSCs and TAMs is of high clinical relevance, as the presence of these cell populations in LUAD are known to facilitate tumor growth and inhibit effective antitumor immune responses, including response to immune checkpoint inhibition (ICI) therapy. The predicted interactions between SFTPD and SIRPA and between APP and CD74, identified using LIANA, is of particular importance. Immunotherapy is effective in only ∼20% of LUAD patients, and the response failure rate to ICI therapy based on CTLA-4/PD-L1 IHC staining remains stubbornly high. Additional disruptions to AT2-immunosuppressive myeloid interactions may hold novel promise in boosting response to ICI therapy by blocking the multiple ways in which malignant AT2 cells interact with the TIME.

While this study provides the first comprehensive characterization of the influence of cell of origin on the immune landscape of LUAD, this study was restricted to the myeloid cell lineage, which does not encompass the entirety of TIME composition and dynamics. Lymphoid, stromal, and endothelial cells also play pivotal roles in TIME, immune surveillance, and suppressive states. Building on this initial study, future endeavors should incorporate a broader spectrum of cell types, while investigating the phenotypic, cellular and genetic changes that influence their interactions with cancer cells to affect suppressive states. Moreover, this study was performed at a single point in tumor development. Longitudinal studies may elicit a deeper understanding of how the TIME evolves alongside tumor progression. Translating these findings to human KRAS^mutant^ tumor samples, including those from diverse demographic and genetic backgrounds, will be critical for translation of this work into the clinical setting.

Taken together, this study demonstrates that the alveolar cell of origin in LUAD significantly influences the immune landscape of the TIME, implying response to immunotherapy may be in part dependent on the cell in which the cancer arose. This finding could be beneficial in optimization of the clinical criteria for immunotherapeutic and combination therapeutic strategies, potentially leading to better patient stratification and improved clinical outcomes. Numerous clinical and preclinical studies in other tumor types demonstrate that use of successful drug combinations that affect myeloid suppression can improve response to checkpoint inhibition^109–115^. Our data suggest a subset of patients with lung adenocarcinomas, ostensibly patients whose tumors originated in AT2 cells, could benefit from this treatment strategy as well. A recent study in which a mathematical model that accounts for the effect of MDSC suppression on tumor progression was validated on a cohort of lung cancer patients, further supporting the importance of suppression on tumor progression^116,117^. This body of work, among others^13,118,119^, provides supporting evidence that LUAD heterogeneity may arise in part from multiple alveolar cell state origins, and design strategies for the next generation of therapeutic strategies in LUAD must begin to incorporate multiple cellular origins into their modeling to improve accuracy of testing and development..

## MATERIALS and METHODS

### Transgenic Mouse models

Detailed descriptions of mouse strains were described recently^13^. In brief, *Sftpc^CreERT2^* mice were a gift from Hal Chapman to Zea Borok, and maintained on a C57BL/6 (WT genotype) background. *Gramd2^CreERT2^*mice were generated by Applied Stem Cell, Inc. (Milpitas, CA) maintained on a C57BL/6 background. LSL-KRAS^G12D^ mice were purchased from JAX labs (B6.129S4-Krastm4Tyj/J; Jackson Laboratory strain #:008179) and maintained on a C57BL/6 background. mTmG mice were purchased from JAX (B6.129(Cg)-Gt(ROSA)26Sortm4(ACTB-tdTomato,-EGFP)Luo/J; Jackson Laboratory strain #:007676) and maintained on a C57BL/6 background. Triple transgenic *Stfpc*:Kras^G12D^:mTmG and *Gramd2*:Kras^G12D^:mTmG mice were generated by crossing either *Stfpc*:Kras^G12D^ or *Gramd2*:Kras^G12D^ heterozygous mice with the mTmG transgenic mouse line. The *Sftpc^CreERT2^* and *Gramd2^CreERT2^* mouse lines have been used previously to track AT2 and AT1 cell fate, respectively^9,13^.

### Mouse genotyping

All mouse procedures underwent ethical review by the Institutional Animal Care and Use Committee (IACUC) at USC prior to experimentation (IACUC Protocol #20963). Mouse tail samples (0.1cm) were dissolved with 200 µL 50 mM NaOH and placed in an oven at 95 °C for 20 min. NaOH was then neutralized with 50 µL 1.0 M Tris-Buffer, and crude DNA was used at a concentration of 50 ng/ul as a template for subsequent genotyping PCR reactions. PCR was performed using Sapphire Amp PCR Mastermix (Takara Bio, RR350B). For *Sftpc^CreERT2^*, reactions consisted of 10 µL PCR master mix, 8 µL ddH2O, 1 µL DNA sample, and 0.5 µL of each primer (requires forward and reverse primers for both mutant and wild-type genes) for a total volume of 20 µL. For the *Gramd2^CreERT2^*, mTmG, and KRAS-LSL-G12D transgenes, an identical PCR recipe was used with the substitution of forward primer pairs to 0.5 µL of a common forward primer used for the mutant and wild-type genes. Primer sequences were as follows: *Gramd2^CreERT2^* common forward: 5’- CTA GTC CTG TCC TCG TCC TAT C-3’, *Gramd2^CreERT2^* mutant allele reverse: 5’- GGG AAA CCA TTT CCG GTT ATT C-3’ *Gramd2^CreERT2^* WT allele reverse: 5’- CAC ATC CCA GCC TTC TCA AA-3’; *Sftpc^CreERT2^* WT allele forward: 5’- TGG TTC CGA GTC CGA TTC TTC-3’, *Sftpc^CreERT2^* WT allele reverse: 5’- CCT TTT GCT CTG TTC CCC ATT A-3’, *Sftpc^CreERT2^* mutant allele forward: 5’- TGA GGT TCG CAA GAA CCT GAT GGA-3’, *Sftpc^CreERT2^* mutant allele reverse: 5’- ACC AGC TTG CAT GAT CTC CGG TAT-3’. Kras^LSL-G12D^ common reverse: 5’- CTG CAT AGT ACG CTA TAC CCT GT-3’, Kras^LSL-G12D^ mutant forward: 5’- GCA GGT CGA GGG ACC TAA TA-3’, Kras^LSL-G12D^ WT forward: 5’- TGT CTT TCC CCA GCA CAG T-3’. mTmG RFP/GFP reporter mice (aka Gt(ROSA)26Sor^tm4(ACTB-tdTomato, -EGFP)Luo^/J^97;^ <u>mTmG</u> WT forward: 5’ CCG GAT TGA TGG TAG TGG TC 3’, mTmG WT reverse: GGC TTA AAG GCT AAC CTG ATG TG 3’, mTmG transgene forward: 5’ AAT CCA TCT TGT TCA ATG CGG GAT C 3’, mTmG transgene reverse: GGA GCG GGA GAA ATG GAT ATG 3’. Next, PCR was performed via using ProFlex PCR System (Thermo Fisher Scientific Inc). *Sftpc^CreERT2^* gene PCR cycling was performed for 3 minutes at 94 °C, 35 cycles of 30 seconds at 94 °C, 30 seconds at 58 °C then 40 seconds at 72 °C, and a final 2-minute product extension at 72 °C. PCR product was then stored at 4 °C. For the *Gramd2^CreERT2^*transgene, the PCR reaction underwent amplification for 2 minutes at 94 °C, 35 cycles of 20 seconds at 94 °C, 30 seconds at 58 °C, 90 seconds at 72 °C, and a final extension of 10 minutes at 72 °C before being stored at 4 °C. For KRAS-LSL-G12D transgene, the PCR reaction was incubated for 3 min at 94 °C, 10 cycles of 30 seconds at 94 °C, 30 seconds at 65 °C (decreasing 0.5 °C per cycle), 90 seconds at 68 °C, 28 cycles of 30 seconds at 94 °C, 30 seconds at 60 °C, 90 seconds at 72 °C, and a final extension of 7 minutes at 72 °C prior to being stored at 10 °C. For the mTmG transgene, the reaction was incubated for 3 min at 94 °C, 10 cycles of 30 seconds at 94 °C, 30 seconds at 65 °C (decreasing 0.5 °C per cycle beginning at cycle 2), 90 seconds at 68 °C, 25 cycles of 30 seconds at 94 °C, 30 seconds at 60 °C, 90 seconds at 72 °C, and a final extension of 7 minutes at 72 °C prior to being stored at 4 °C.

### Gel electrophoresis

20 µL sample volume of PCR product was loaded on to a 2% agarose gel (BioPioneer, C0009) with GelRed (BioTium, 41003-T), combined with 6 µL dyed DNA ladder, made up of 1 uL DNA Ladder (New England Biolabs, N3231L), 1 µL Purple Dye (New England Biolabs, B7025S), and 4 uL ddH20. Genotyping gels were run at 100V for 40 minutes prior to being imaged on a ChemiDoc Imaging System (Bio-Rad).

### Single nucleus RNA sequencing

Mice were euthanized with 150-200 mg/kg Euthasol (ANADA #200-071, Virbac) via intraperitoneal (IP) injection, followed by secondary cervical dislocation. Mouse lungs were perfused with 20 mL phosphate buffered saline (PBS). Lungs were snap frozen in liquid nitrogen and subjected to nuclear isolation using the 10X Genomics Nuclei Isolation Kit (10X, 1000494), using the standard protocol with modification only to the lysis reaction time, which was adjusted to 8 minutes. Sequencing libraries were generated using the 10X Genomics Next GEM Single Cell 3’ Gene Expression protocol and sequenced on a Novaseq6000 S2 flow cell run at 28×90 cycles in the USC Molecular Genomics Core.

### Single nucleus resolved transcriptomic data analyses

#### snRNA-seq data quality control and preprocessing

Raw FASTQ sequencing files in the snRNA seq dataset were aligned to the GRCm39 v109 genome that had been modified into a custom file to include the *mTmG* transgene^120^ using the 10x Genomics Cell Ranger pipeline (Count v7.1). Cells were then filtered to include only those with 100–10,000 genes detected, 90,000 Unique molecular identifiers (UMIs) counted, and their fraction of mitochondrial reads <= 20%. 152,000 high-quality cells from 14 mouse lung samples were selected for further analysis. The resulting gene expression matrices were normalized to total unique molecular identifier (UMI) counts for each cell and subsequently transformed using the natural logarithm scale. To correct for technical and biological variations and enhance the accuracy of cell type identification, Harmony was used to integrate multiple biological replicates and minimize technical batch effects. Next, the merged and integrated Seurat object underwent data normalization via the LogNormalize global scaling function in Seurat^121^. The number of principal components (PCs) for different cell types was selectively chosen based on the knee point of the scree plot for each cell type, using the ElbowPlot function in Seurat. This approach accommodates varying population complexities. For downstream analysis, the top 20 PCs were used to run the FindNeighbors and RunUMAP functions. Cell clustering was performed using the FindClusters function, with resolutions ranging from 0.1 to 0.8 explored for optimal clustering. Specifically, a resolution of 0.3 was used for the overall cells. The two-dimensional Uniform Manifold Approximation and Projection (UMAP) method was employed to visualize cell clusters.

#### Cell type annotation

Automated cell type prediction for myeloid cells was performed using LungMAP^47^. For LungMAP annotation, 10X CellRanger hdf5 files were uploaded to the LungMAP Azimuth^47^. Mouse Lung CellRef online application, and cells were mapped to Reference Metadata of lineage_level1, lineage_level2, celltype_level1, celltype_level2, and celltype_level3. The predicted cell types and scores were downloaded and appended to each sample in R. Annotation results were refined by incorporating annotation results from the singleR^122^ and the Immgen Datasets in The Immunological Genome Project (ImmGen)^123^. The final processed Seurat object is available as “Supplemental File 1_Myeloid_Annotated.RDS”.

#### GSEA and pathway analysis

Differentially expressed genes (DEGs) were detected using the Seurat function FindAllMarkers with FDR-corrected p-values. Gene ontology (GO) analysis^124^ and immunologic signatures collection (ImmuneSigDB^125^) were performed with the full list of significant (FDR p <0.05) DEGs using the gene set enrichment analysis (GSEA) function implemented in fgsea package, as well as PANTHER^77,78^, then plotted in R using ggplot2^126^. R code used in this analysis is available as Supplemental File 2.

#### Trajectory analysis

CellRank2^127^ was used to establish the transition matrix based on diffusion pseudotime (DPT)^128^. The *Cd33^+^*population of macrophages was selected as the starting point for the DPT analysis. The transition matrix was visualized by plotting random walks which iteratively choose the next cell based on the current cell’s transition probability. The Generalized Perron Cluster Cluster Analysis (GPCCA) estimator was used to compute macrostates, fate probabilities and gene expression trends from the transition matrix from the transition matrix generated in CellRank’s PseudotimeKernel^129^. Code used for macrophage trajectory analyses is available in Supplemental File 3.

#### Cell-cell communication analysis

InferCNV^100^ was used to identify malignant epithelial cells by first using the raw expression counts and the gene-ordering file, gencode_v21_gen_pos.complete.txt (https://data.broadinstitute.org/Trinity/CTAT/cnv/). Then, copy number variation (CNV) was estimated using infercnv::run function by mapping genes to their chromosomal positions and applying a moving average to their relative expression levels. Subsequently, CNV score for each cell was obtained from “infercnv.observation.txt”, one of output files from the prescribed step, and a cutoff of CNV score was determined at the 95 percentiles of the CNV score of normal cells. All epithelial cells within LUAD samples that exceed the cutoff value were identified as malignant LUAD cells, and therefore isolated out and integrated with myeloid cells from LUAD samples for downstream CCC analysis. The new integrated Seurat objects were saved as “Supplemental File 4_CCCObj_in_AT1LUAD.RDS” and “Supplemental File 5_CCCObj_in_AT2LUAD.RDS”. CCC analysis was performed via using LIANA package (v0.1.12)^99^ with standard parameters (https://saezlab.github.io/liana/articles/liana_tutorial.html). LIANA R code is listed Supplemental File 6. MouseConsensus was applied among CCC resources with the liana_wrap function. Correlation matrix plotting of cell-cell interactions was performed using interactions in the *Gramd2^+^* KRAS^G12D^ LUAD model by subtracting corresponding interactions in the *Sftpc^+^* KRAS^G12D^ LUAD model. Red = higher number of interactions in the *Gramd2^+^* KRAS^G12D^ LUAD model; blue: higher number of interactions in the *Sftpc^+^* KRAS^G12D^ LUAD model. R code used for the CCC analysis is available as Supplemental File 6.

### Spatially resolved transcriptomic (ST-seq) analyses

The ST-seq dataset and analysis pipeline used throughout this study were previously published^13^.

#### Integration of the snRNA and the ST-seq datasets

Integration of snRNA-seq and ST-seq dataset was performed using Seurat V5. Initially, transfer anchors were identified between the snRNA-seq dataset (Supplemental File 1) and the ST-seq dataset (Supplemental File 7) using the FindTransferAnchors function. SCTransform (SCT) normalization was applied to both datasets prior to integration to ensure data compatability. Anchors facilitated accurate label transfer, performed using the TransferData function. Cell type annotations for myeloid cells from the snRNA-seq dataset (reference) were then mapped onto the ST-seq dataset (query). The result of this transfer process was stored in a new assay, termed “Integration,” within the Seurat object. To ensure that the integrated data was used for downstream analyses, the default assay was set to “Integration.” For the indicated queries.

#### Tumor lymphocyte signature (TLS) scoring

TLS scoring was performed on the ST-seq dataset set to SCTransform (SCT) normalization using Seurat V5. The AddModuleScore function in Seurat was used to calculate a module score for each cell for composite TLS gene signature per default program parameters. The resulting TLS score was integrated as a unique metadata column into the ST-seq Seurat object. Visualization of the TLS scores was performed using UMAP as well as spatial plots to depict TLS distribution across the dataset. Violin plots were generated to compare TLS scores across different experimental groups, with normality determined using an Anderson-Darling post-test and statistical significance determined using a Wilcoxon test. R code describing the analysis of the ST-seq dataset is available as Supplemental File 8.

### Flow cytometry

Mouse whole lung tissue was dissociated into a single-cell suspension using the Miltenyi Biotec Tumor Dissociation Kit, mouse (#130-096-730) and subsequently passed through a 70 µm strainer (Miltenyi Biotec, #130-098-462). Red blood cells were then removed by resuspending cells in 2 mL lysis buffer (Gibco, A1049201) for 2 minutes at room temperature. The lysis reaction was quenched using 15 mL TIL media (RPMI-1640 with 10% FBS, 1% penicillin/streptomycin, and 0.5% L-glutamine) before being resuspended in 1 mL TIL media. Cells were plated at 1.0 x 10^6^ cells/well in a 96-well u-bottom plate. Cells were washed with 200 µL PBS, resuspended in 100 µL Live/Dead Fixable Aqua stain (Invitrogen, L34966) at 1:1000 dilution in PBS, and incubated at 4°C for 30 minutes in the absence of light. Cells were then washed three times with 200 µL FACS buffer (DPBS with 2% FBS). After adding 100 µL FcR blocking solution (1:50, 101302, Biolegend) to samples for 10 minutes at room temperature, cells were stained for cell-surface markers of the Macrophage and MDSC panels. Surface staining of the Macrophage panel consisted of the following primary antibodies diluted in FACS buffer (2% FBS in DPBS) with anti-mouse CD45-Brilliant Violet 650 (1:200, 103151, Biolegend), anti-mouse F4/80-Brilliant Violet 785 (1:200, 123141, Biolegend), anti-mouse Ly6g-FITC (1:300, 127605, Biolegend), anti-mouse/human CD11b-PE/Dazzle 594 (1:120, 101256, Biolegend), and anti-mouse MHCII APC/Cyanine7 (1:120, 107628, Biolegend). The following isotype control antibodies were diluted in FACS Buffer to be used in the macrophage surface staining panel: Brilliant Violet 650 Rat IgG2b, k (400651, Biolegend), Brilliant Violet 785 Rat IgG2a, k (400546, Biolegend), FITC Rat IgG2a, k (400505, Biolegend), PE/Dazzle 595 Rat IgG2b, k (400660, Biolegend), APC/Cyanine7 Rat IgG2b, k (400624, Biolegend). Surface staining of the MDSC panel consisted of the following primary antibodies diluted in FACS buffer: anti-mouse CD45-Brilliant Violet 650 (1:200, 103151, Biolegend), anti-mouse F4/80-Brilliant Violet 785 (1:200, 123141, Biolegend), anti-mouse Ly6g-FITC (1:300, 127605, Biolegend), anti-mouse Ly6c-PerCP/Cyanine5.5 (1:100, 128012, Biolegend), and anti-mouse/human CD11b-PE (1:200, 101207, Biolegend). The following isotype antibodies were diluted in FACS Buffer to be used in the MDSC surface staining panel: Brilliant Violet 650 Rat IgG2b, k (400651, Biolegend), Brilliant Violet 785 Rat IgG2a, k (400546, Biolegend), FITC Rat IgG2a, k (400505, Biolegend), PerCP/Cyanine5.5 Rat IgG2c, k (400723, Biolegend), Rat IgG2b kappa, PE (12-4031-82, Invitrogen). 50 µL of antibody mastermix solutions were added to each well and incubated at 4°C for 1 hour in the absence of light. Cells were washed three times with 200 µL FACS buffer. Fixation and permeabilization was then performed for 1 hour at 4°C using the CytoFast Fix/Perm Set (426803, Biolegend). After fixation/permeabilization, samples were washed with 200 µL CytoFast wash buffer, underwent a second Fc blocking procedure, and then stained for the following intracellular markers. The Macrophage panel’s intracellular primary antibodies consisted of anti-mouse/human Arginase 1-eFluor 450 (1:50, 48-3697-82, Invitrogen) and anti-mouse Ki67-PE/Cyanine7 (1:150, 25-5698-82, Invitrogen), while the MDSC panel’s intracellular primary antibodies consisted of anti-mouse/human Arginase 1-eFluor 450 (1:300, 48-3697-82, Invitrogen). The Macrophage panel’s intracellular isotype antibodies consisted of Rat IgG2a kappa, eFluor 450 (48-4321-82, Invitrogen) and PE/Cyanine7 Rat IgG2a, k (400521, Biolegend), while the MDSC panel’s intracellular isotype antibodies consisted of Rat IgG2a kappa, eFluor 450 (48-4321-82, Invitrogen). Primary antibodies for intracellular staining were diluted in CytoFast wash buffer. 50 µL of intracellular antibody master mix were added to each sample and incubated at 4°C for 1 hour in the absence of light. Samples were washed twice with 200 µL CytoFast wash buffer and washed once with 200 µL of FACS buffer before being resuspended in 200 µL FACS buffer. Samples were characterized using an Attune NxT Flow Cytometer (A24858, ThermoFisher Scientific). Single-color compensation for each channel was performed with UltraComp Compensation Beads (01-222-41, Invitrogen) stained with isotype antibodies for every fluorophore. MLE-15 cells were used as unstained and Live/Dead Fixable Aqua single-color compensation in the MDSC panel, while freshly dissociated whole mouse lung cells were used for unstained and Live/Dead Fixable Aqua single-color compensation in the Macrophage panel. Data was analyzed using FCS Express 7 (Version 7.22.006) Software (De Novo).

### Cell lines

MLE-15 cell line was established by Dr. Jeffrey A. Whitsett (Children’s Hospital Medical Center, Cincinnati, OH)^130^, and kindly gifted by our collaborator, Z. Beiyun lab, University of Southern California. MLE-15 cells were cultivated in HITES medium (RPMI-1640 medium supplemented with 10 nM hydrocortisone, 5 μg/ml insulin, 5 μg/ml human transferrin, 10 nM β-estradiol, 5 μg/ml selenium, 2 mM L-glutamine, 10 mM HEPES, 100 U/ ml penicillin, 100 μg/ml streptomycin and 4% FBS (Atlanta Biologicals, Flowery Branch, GA).

### Data accessibility

Visium 10X ST-seq profiling was previously published^13^ and is available via the NCBI Gene Expression Omnibus (GEO) as GSE215858. SnRNAseq generated as part of this study is also available through GEO as GSE267731.

## ACKNOWLEDGEMENTS

The authors would like to acknowledge Dr. Beiyun Zhou for her gift of the MLE-15 cell line, originally generated in the laboratory of Dr. Whitsett at the University of Cincinnati. The authors would also like to thank Stephanie Tring and the USC Molecular Genomics Core for generation of the sequencing data used in this study, the USC Stem Cell flow cytometry facility for assistance performing FACS analysis, and the USC Norris Tissue Pathology Core for their assistance in embedding and sectioning of lung tissues. All cores were supported by the USC Norris Comprehensive Cancer Center Core grant from NCI (P30 CA014089). The authors would also like to thank colleagues from the Marconett, Roussos Torres, and Raz labs for critical discussion of the study. Funding for this study was provided by the NCI to Dr. Marconett (R01 CA262258), and to Dr. Roussos Torres (R01 CA283169). In addition, Dr. Marconett was funded in part by an American Cancer Society Research Scholar Grant (RMC-RSG-20-135-01); Dr. Borok was supported by the NHLBI (R35 HL135747); Dr. Raz was supported by the Department of Defense (W81XWH-22-1-0306). This study was supported in part by the Department of Translational Genomics at USC as well as the Department of Integrative Translational Sciences at the Beckman Research Institute, City of Hope.

## Supplementary

**Supplemental Figure 1:**
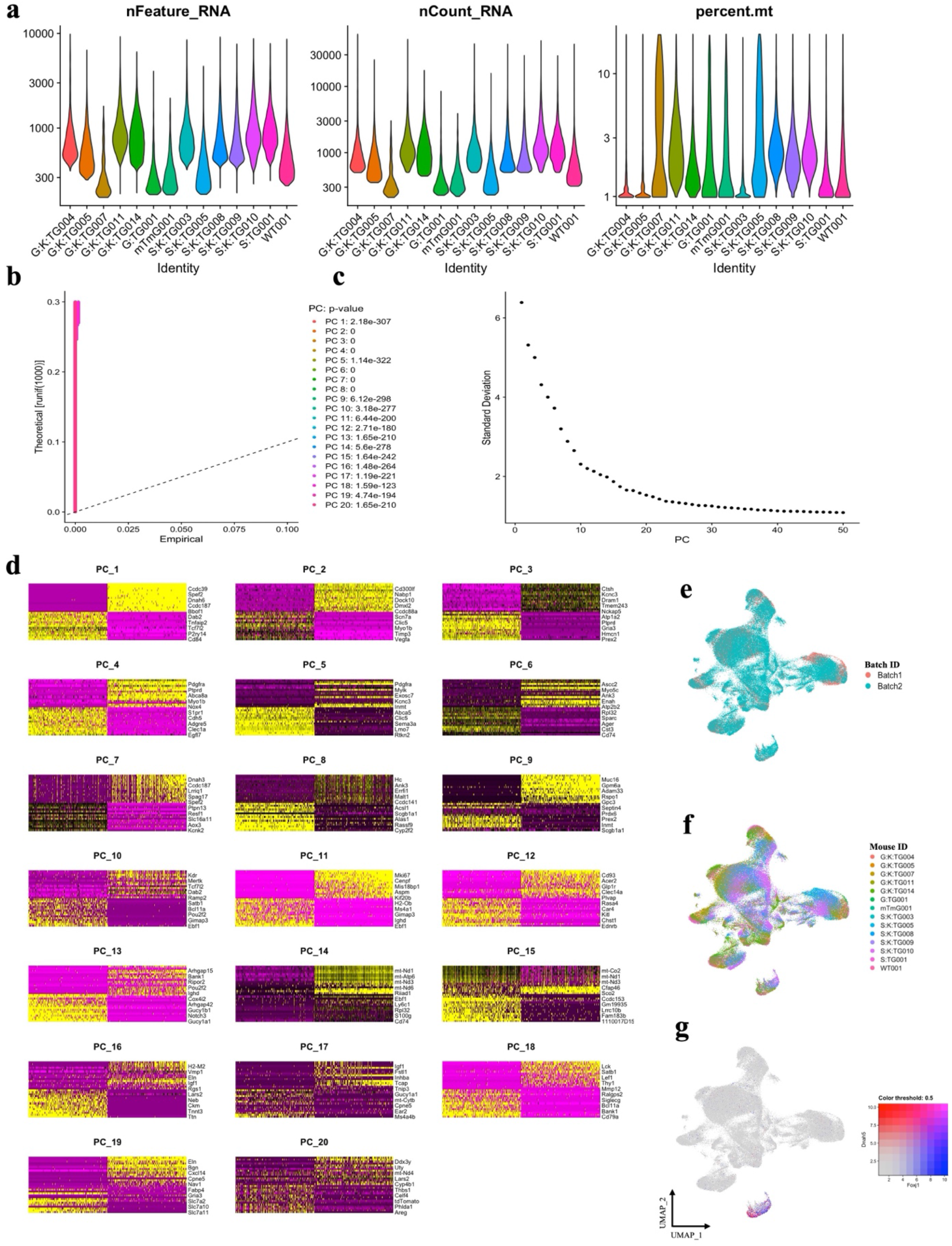
Quality control and processing metrics for mouse snRNAseq dataset. a, Quality control metrics for raw data in SeuratObject, all experimental groups are notated as in Table 1. b, Jack Straw plot for significance of principal components in the mouse snRNAseq data cohort. c, Elbow plot showing significance of each PCA within the snRNAseq object. d, Heatmaps showing differential gene expression contributing to each of the top 20 principal components within the snRNAseq dataset. Purple = little to no expression, yellow = high expression. e, UMAP of snRNAseq dataset. Colors indicate batch sample(s) were sequenced in. Salmon = batch 1, teal = batch 2. f, UMAP of snRNAseq dataset. Colors indicate individual mice cells were isolated from. Red = *Gramd2^+^* KRAS^G12D^ LUAD-004 (G:K:TG004), orange = *Gramd2^+^* KRAS^G12D^ LUAD-005 (G:K:TG005), gold = *Gramd2^CreERT2:^* KRAS^-^LSL-G12D, vehicle control-007 (G:K:TG007), peridot = *Gramd2^+^* KRAS^G12D^ LUAD-011 (G:K:TG011), forest green = *Gramd2^CreERT2:^* KRAS^-^LSL-G12D vehicle control-014 (G:K:TG014), green = *Gramd2^CreERT2^*; mTmG control-001 (G:TG001), seafoam = mTmG control-001 (mTmG001), teal = *Sftpc^CreERT2^*KAS-LSL-G12D vehicle control-003 (S:K:TG003), blue = *Sftpc^+^* KRAS^G12D^ LUAD-005 (S:K:TG005), dark blue = *Sftpc^+^* KRAS^G12D^ LUAD-008 (S:K:TG008), purple = *Sftpc^CreERT2^* KAS-LSL-G12D vehicle control-009 (S:K:TG009), violet = *Sftpc^+^* KRAS^G12D^ LUAD-010 (S:K:TG010), maroon = *Sftpc^CreERT2^* mTmG control-001 (S:TG001), and pink = wild type-001 (WT001). g, UMAP comparing *Dnah5* (Red) and *Foxj1* (blue) to identify ciliated cells.

**Supplemental Figure 2:**
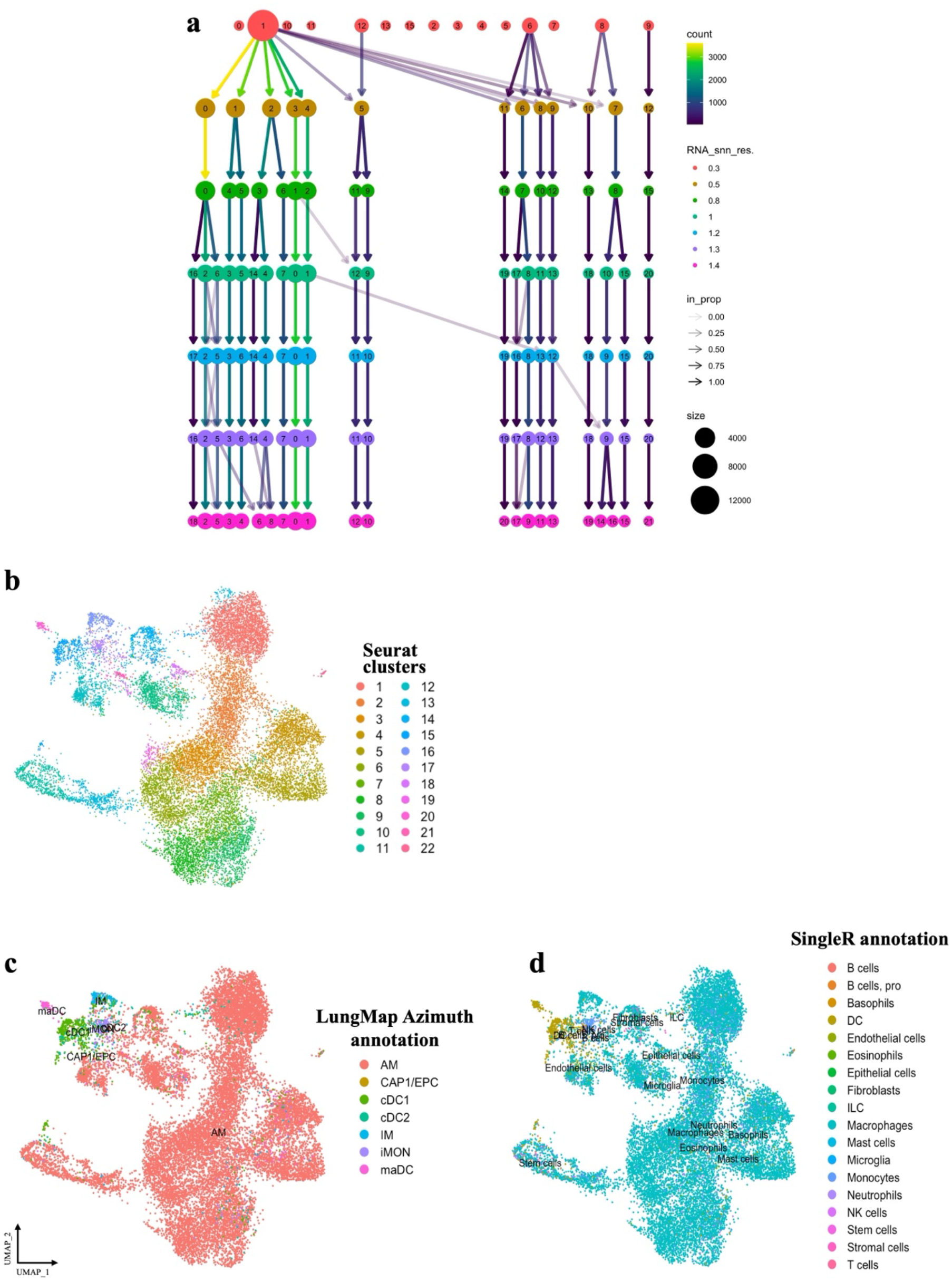
Optimization and visualization of snRNAseq clustering. a, Tree diagram of mouse snRNAseq clustering results containing myeloid cells from all experimental groups. Nodes are colored according to the value of RNA_snn_resolution and sized according to the number of nuclei they represent. Edges are colored according to the number of samples (from blue representing few to yellow representing many). The transparency is adjusted according to the in-proportion, with stronger lines showing edges that are more important to the higher-resolution cluster. Cluster labels are randomly assigned by the k-means algorithm. b, UMAP plot of single nucleus transcriptomic expression data of myeloid cells colored by Seurat clusters (dims = 1:21; resolution = 1.4). c,UMAP plot of single nucleus transcriptomic expression data of myeloid cells colored by myeloid cell types annotated in Azimuth^47^ according to LungMAP mouse reference (predicted cell type level2). d,UMAP plot of single nucleus transcriptomic expression data of myeloid cells colored by myeloid cell types annotated by SingleR.

**Supplemental Figure 3:**
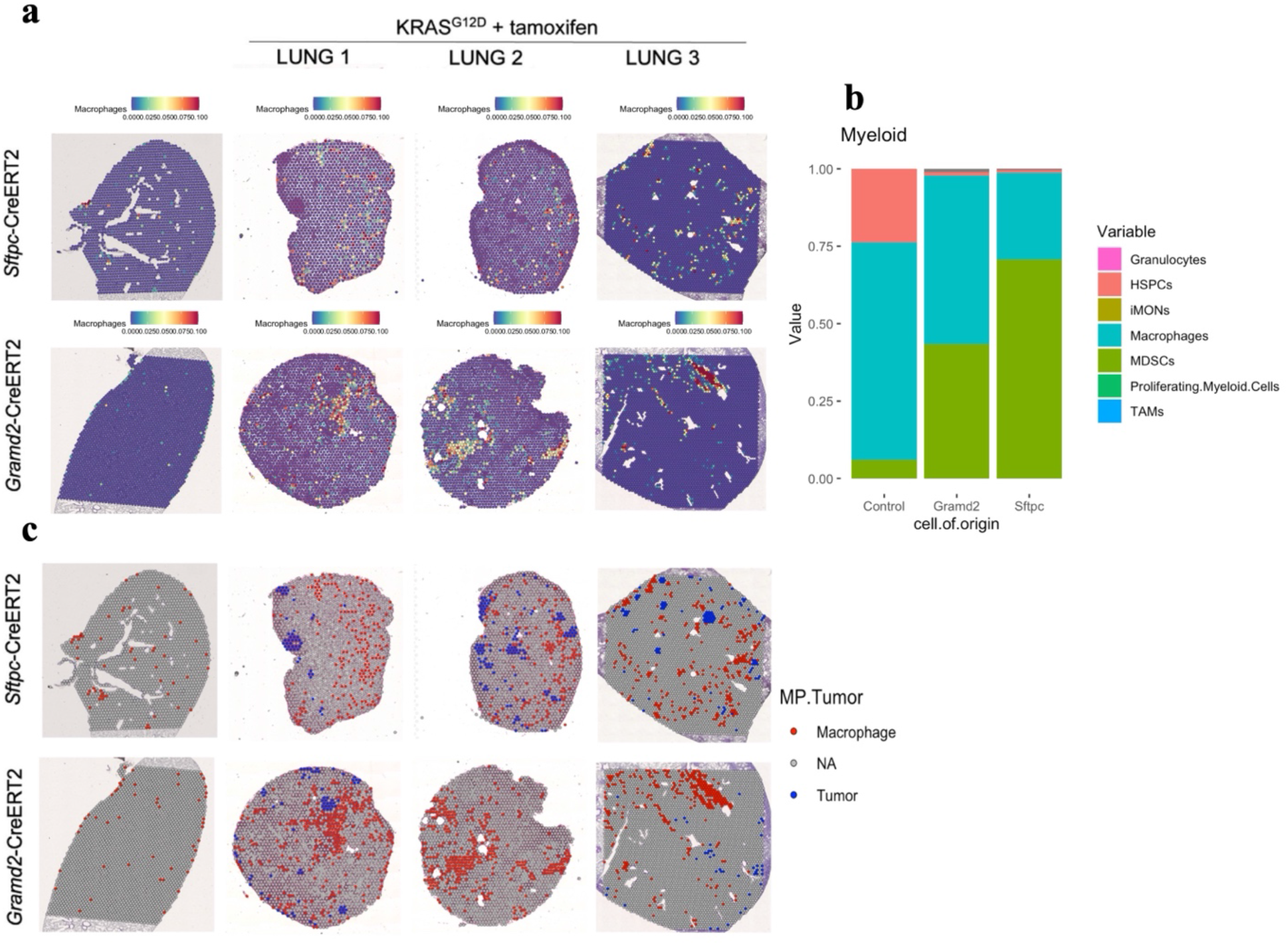
ST-seq analysis of mouse experimental groups for myleoid cell composition. a, SnRNAseq integration onto ST-Seq data for composite macrophage signature. Low macrophage score = blue, high macrophage score = red. b, Stacked bar chart of myeloid subpopulation proportions based on values generated from snRNAseq integration onto ST-Seq data, separated by cell of origin and CreERT2 controls. Pink = granulocytes, salmon = HSPCs, peridot = iMONs, teal = macrophages, green = MDSCs, seafoam = proliferating myeloid cells, blue = TAMs. c, Spatial representation of tumor- and macrophage-containing spots to visualize proximity of these cell types. Macrophage-containing spots were defined as spots that have a Seurat snRNAseq integration value greater than zero, indicating that a macrophage signature comprises some composition of any given spot. Tumor-containing spots were defined as spots that have a Tumor Purity Score greater than 0.84, as designated by the R package, ESTIMATE. The Tumor Purity threshold value of 0.84 excluded 99% of the spots from the CreERT2 control samples. Red = macrophages, blue = LUAD tumor, grey = neither.

**Supplemental Figure 4:**
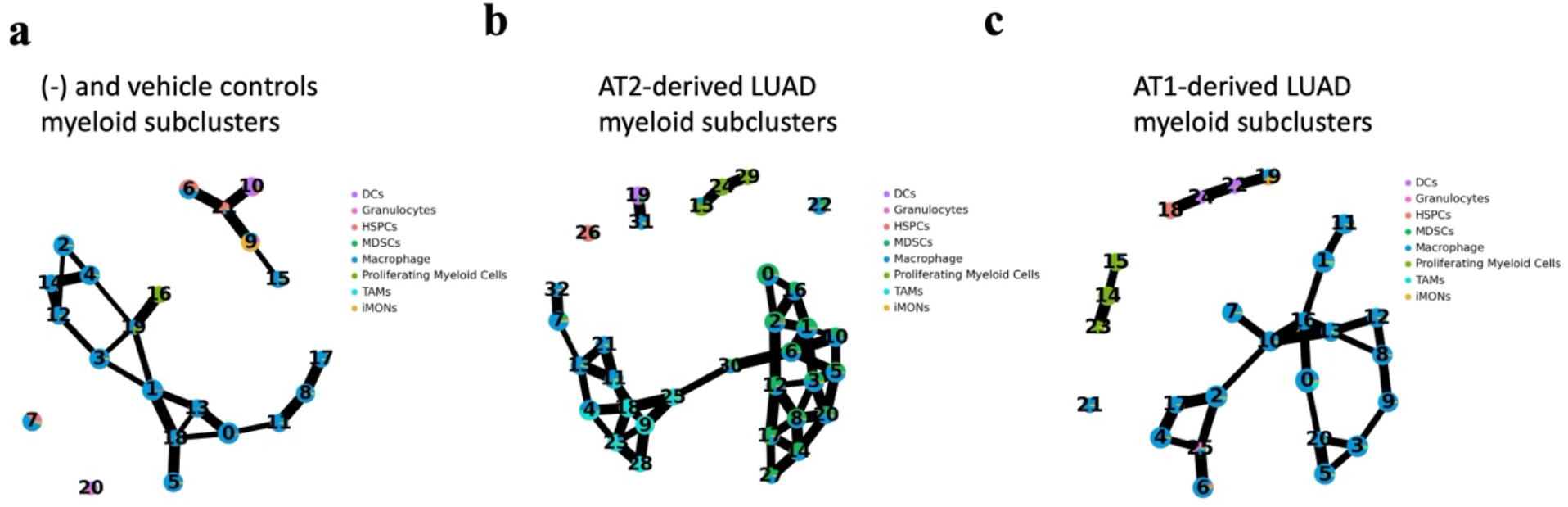
Trajectory analysis PAGA diagrams split by experimental group. a, PAGA analysis of combined negative (-) and vehicle controls for myeloid lineage trajectory analysis. Colors indicate cell type identity. Purple = DCs, pink = granulocytes, red = HSPCs, teal = MDSCs, blue = macrophages, green = proliferating myeloid, light blue = TAMs, gold = iMONs. b, PAGA analysis of *Sftpc^+^* KRAS^G12D^ LUAD for myeloid lineage trajectory analysis. Colors indicate cell type identity. Colors as in (a). c, PAGA analysis of *Gramd2^+^* KRAS^G12D^ LUAD for myeloid lineage trajectory analysis. Colors indicate cell type identity. Colors as in (a).

**Supplemental Figure 5:**
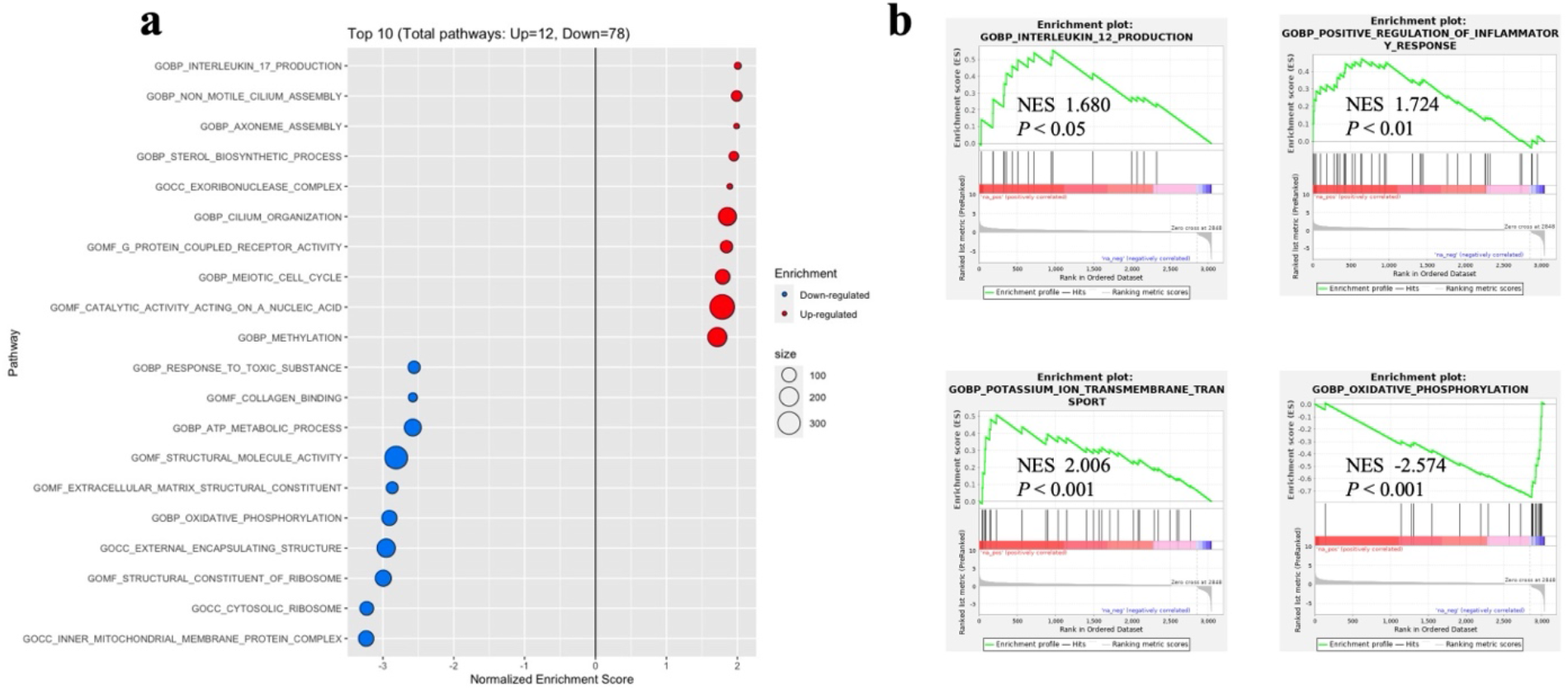
GSEA pathways enrichment analysis between *Sftpc^+^* KRAS^G12D^ LUAD and *Gramd2^+^* KRAS^G12D^ LUAD macrophages. a, GSEA for clusters of experimental group against mouse ontology gene sets. Top 10 significantly enriched pathways are displayed. Red = enriched in *Gramd2^+^* KRAS^G12D^ LUAD macrophages, blue = enriched in *Sftpc^+^* KRAS^G12D^ LUAD macrophages. Size of dot indicates gene count assigned to each pathway from GSEA, small dot = ∼100 genes in pathway, large dot = ∼300 genes in pathway. X-axis = normalized enrichment score. NES = enrichment score normalized to mean enrichment of random samples of the same size. b, Gene sets for leukocyte chemotaxis and migration were significantly enriched. The y-axis represents enrichment score (NES) and on the x-axis are genes (vertical black lines) represented in gene sets. The green line connects points of ES and genes. ES is the maximum deviation from zero as calculated for each gene going down the ranked list and represents the degree of over-representation of a gene set at the top or the bottom of the ranked gene list. The colored band at the bottom represents the degree of correlation of genes with the asthma phenotype (red for positive and blue for negative correlation). Significance threshold set at FDR < 0.05.

