## Supplemental File 2 for "Cell of origin alters myeloid-mediated immunosuppression in lung adenocarcinoma": Supplemental File 3_r code for snRNA-seq analyses.R

```
############################################################################################################################################################
############################################################## FIGURE 1 & Supplemental FIGURE1############################################################## 
############################################################################################################################################################
install.packages('EnsDb.Mmusculus.v79')
library(EnsDb.Mmusculus.v79) # for analysing mouse data
library(patchwork)
library(Seurat)
library(GenomeInfoDb)
library(ggplot2)
library(patchwork)
library(dplyr)
library(biovizBase)
library(hdf5r)
library(harmony)
library(tidyverse)
library(fgsea)
library(infercnv)
library(dendextend)
set.seed(1234)


#Create Seueat Objects for 8 sampels in Batch 1
Batch1_Sample1 <- Read10X_h5("your/path/here/filtered_feature_bc_matrix (1).h5", use.names = TRUE, unique.features = TRUE)
Batch1_Sample2 <- Read10X_h5("your/path/here/filtered_feature_bc_matrix (2).h5", use.names = TRUE, unique.features = TRUE)
Batch1_Sample3 <- Read10X_h5("your/path/here/filtered_feature_bc_matrix (3).h5", use.names = TRUE, unique.features = TRUE)
Batch1_Sample4 <- Read10X_h5("your/path/here/filtered_feature_bc_matrix (4).h5", use.names = TRUE, unique.features = TRUE)
Batch1_Sample5 <- Read10X_h5("your/path/here/filtered_feature_bc_matrix (5).h5", use.names = TRUE, unique.features = TRUE)
Batch1_Sample6 <- Read10X_h5("your/path/here/filtered_feature_bc_matrix (6).h5", use.names = TRUE, unique.features = TRUE)
Batch1_Sample7 <- Read10X_h5("your/path/here/filtered_feature_bc_matrix (7).h5", use.names = TRUE, unique.features = TRUE)
Batch1_Sample8 <- Read10X_h5("your/path/here/filtered_feature_bc_matrix (8).h5", use.names = TRUE, unique.features = TRUE)

#add metadata columns/additional information to each Seurat object
Batch1_Sample1 <- CreateSeuratObject(counts = Batch1_Sample1, project = "AT1-LUAD", min.cells = 3, min.features = 200)
Batch1_Sample1$Genotype <- c("Sftpc:Kras:mTmG")
Batch1_Sample1$Disease_Status <- c("LUAD")
Batch1_Sample1$Cell_of_Origin <- c("AT2-derived LUAD")
Batch1_Sample1$Mouse_ID <- c("S:K:TG005")
Batch1_Sample1$Batch <- c("Batch1")
prediction11 <- read.delim("your/path/here/azimuth_pred (1).tsv", row.names = 1)
Batch1_Sample1 <- AddMetaData(object = Batch1_Sample1, metadata = prediction11)

Batch1_Sample2 <- CreateSeuratObject(counts = Batch1_Sample2, project = "AT1-LUAD", min.cells = 3, min.features = 200)
Batch1_Sample2$Genotype <- c("Sftpc:Kras:mTmG")
Batch1_Sample2$Disease_Status <- c("Sp_LUAD")
Batch1_Sample2$Cell_of_Origin <- c("Spontaneous_AT2-derived LUAD")
Batch1_Sample2$Mouse_ID <- c("S:K:TG003")
Batch1_Sample2$Batch <- c("Batch1")
prediction12 <- read.delim("your/path/here/azimuth_pred (2).tsv", row.names = 1)
Batch1_Sample2 <- AddMetaData(object = Batch1_Sample2, metadata = prediction12)

Batch1_Sample3 <- CreateSeuratObject(counts = Batch1_Sample3, project = "AT1-LUAD", min.cells = 3, min.features = 200)
Batch1_Sample3$Genotype <- c("Gramd2:Kras:mTmG")
Batch1_Sample3$Disease_Status <- c("LUAD")
Batch1_Sample3$Cell_of_Origin <- c("AT1-derived LUAD")
Batch1_Sample3$Mouse_ID <- c("G:K:TG004")
Batch1_Sample3$Batch <- c("Batch1")
prediction13 <- read.delim("your/path/here/azimuth_pred(3).tsv", row.names = 1)
Batch1_Sample3 <- AddMetaData(object = Batch1_Sample3, metadata = prediction13)

Batch1_Sample4 <- CreateSeuratObject(counts = Batch1_Sample4, project = "AT1-LUAD", min.cells = 3, min.features = 200)
Batch1_Sample4$Genotype <- c("Gramd2:Kras:mTmG")
Batch1_Sample4$Disease_Status <- c("LUAD")
Batch1_Sample4$Cell_of_Origin <- c("AT1-derived LUAD")
Batch1_Sample4$Mouse_ID <- c("G:K:TG005")
Batch1_Sample4$Batch <- c("Batch1")
prediction14 <- read.delim("your/path/here/azimuth_pred(4).tsv", row.names = 1)
Batch1_Sample4 <- AddMetaData(object = Batch1_Sample4, metadata = prediction14)

Batch1_Sample5 <- CreateSeuratObject(counts = Batch1_Sample5, project = "AT1-LUAD", min.cells = 3, min.features = 200)
Batch1_Sample5$Genotype <- c("Gramd2:Kras:mTmG")
Batch1_Sample5$Disease_Status <- c("Healthy")
Batch1_Sample5$Cell_of_Origin <- c("Vehicle_Controls")
Batch1_Sample5$Mouse_ID <- c("G:K:TG007")
Batch1_Sample5$Batch <- c("Batch1")
prediction15 <- read.delim("your/path/here/azimuth_pred(5).tsv", row.names = 1)
Batch1_Sample5 <- AddMetaData(object = Batch1_Sample5, metadata = prediction15)

Batch1_Sample6 <- CreateSeuratObject(counts = Batch1_Sample6, project = "AT1-LUAD", min.cells = 3, min.features = 200)
Batch1_Sample6$Genotype <- c("Gramd2:mTmG")
Batch1_Sample6$Disease_Status <- c("Healthy")
Batch1_Sample6$Cell_of_Origin <- c("Neg_Controls")
Batch1_Sample6$Mouse_ID <- c("G:TG001")
Batch1_Sample6$Batch <- c("Batch1")
prediction16 <- read.delim("your/path/here/azimuth_pred(6).tsv", row.names = 1)
Batch1_Sample6 <- AddMetaData(object = Batch1_Sample6, metadata = prediction16)

Batch1_Sample7 <- CreateSeuratObject(counts = Batch1_Sample7, project = "AT1-LUAD", min.cells = 3, min.features = 200)
Batch1_Sample7$Genotype <- c("mTmG")
Batch1_Sample7$Disease_Status <- c("Healthy")
Batch1_Sample7$Cell_of_Origin <- c("Neg_Controls")
Batch1_Sample7$Mouse_ID <- c("mTmG001")
Batch1_Sample7$Batch <- c("Batch1")
prediction17 <- read.delim("your/path/here/azimuth_pred(7).tsv", row.names = 1)
Batch1_Sample7 <- AddMetaData(object = Batch1_Sample7, metadata = prediction17)

Batch1_Sample8 <- CreateSeuratObject(counts = Batch1_Sample8, project = "AT1-LUAD", min.cells = 3, min.features = 200)
Batch1_Sample8$Genotype <- c("WT")
Batch1_Sample8$Disease_Status <- c("Healthy")
Batch1_Sample8$Cell_of_Origin <- c("Neg_Controls")
Batch1_Sample8$Mouse_ID <- c("WT001")
Batch1_Sample8$Batch <- c("Batch1")
prediction18 <- read.delim("your/path/here/azimuth_pred(8).tsv", row.names = 1)
Batch1_Sample8 <- AddMetaData(object = Batch1_Sample8, metadata = prediction18)

#merge 8 Seurat objects into one
AT1LUAD_MergedBatch1 <- merge(Batch1_Sample1, y = c(Batch1_Sample2, Batch1_Sample3, Batch1_Sample4,Batch1_Sample5, Batch1_Sample6, Batch1_Sample7, Batch1_Sample8),
                              add.cell.ids = ls()[1:8],
                              project = "AT1-LUAD")

#save
saveRDS(AT1LUAD_MergedBatch1, file = "your/path/here/AT1LUAD_MergedBatch1.RDS")


#Create Seurat Objects for 6 samples in Batch 2
Batch2_Sample1 <- Read10X_h5("your/path/here/STG001_filtered_feature_bc_matrix.h5", use.names = TRUE, unique.features = TRUE)
Batch2_Sample2 <- Read10X_h5("your/path/here/SKTG009_filtered_feature_bc_matrix.h5", use.names = TRUE, unique.features = TRUE)
Batch2_Sample3 <- Read10X_h5("your/path/here/SKTG008_filtered_feature_bc_matrix.h5", use.names = TRUE, unique.features = TRUE)
Batch2_Sample4 <- Read10X_h5("your/path/here/SKTG010_filtered_feature_bc_matrix.h5", use.names = TRUE, unique.features = TRUE)
Batch2_Sample5 <- Read10X_h5("your/path/here/GKTG014_filtered_feature_bc_matrix.h5", use.names = TRUE, unique.features = TRUE)
Batch2_Sample6 <- Read10X_h5("your/path/here/GKTG011_filtered_feature_bc_matrix.h5", use.names = TRUE, unique.features = TRUE)

#add metadata columns/additional information to each Seurat object
Batch2_Sample1 <- CreateSeuratObject(counts = Batch2_Sample1, project = "AT1-LUAD", min.cells = 3, min.features = 200)
Batch2_Sample1$Genotype <- c("Sftpc:mTmG")
Batch2_Sample1$Disease_Status <- c("Healthy")
Batch2_Sample1$Cell_of_Origin <- c("Neg_Controls")
Batch2_Sample1$Mouse_ID <- c("S:TG001")
Batch2_Sample1$Batch <- c("Batch2")
prediction21 <- read.delim("your/path/here/azimuth_pred (1).tsv", row.names = 1)
Batch2_Sample1 <- AddMetaData(object = Batch2_Sample1, metadata = prediction21)

Batch2_Sample2 <- CreateSeuratObject(counts = Batch2_Sample2, project = "AT1-LUAD", min.cells = 3, min.features = 200)
Batch2_Sample2$Genotype <- c("Sftpc:Kras:mTmG")
Batch2_Sample2$Disease_Status <- c("Sp_LUAD")
Batch2_Sample2$Cell_of_Origin <- c("Spontaneous_AT2-derived LUAD")
Batch2_Sample2$Mouse_ID <- c("S:K:TG009")
Batch2_Sample2$Batch <- c("Batch2")
prediction22 <- read.delim("your/path/here/azimuth_pred (2).tsv", row.names = 1)
Batch2_Sample2 <- AddMetaData(object = Batch2_Sample2, metadata = prediction22)

Batch2_Sample3 <- CreateSeuratObject(counts = Batch2_Sample3, project = "AT1-LUAD", min.cells = 3, min.features = 200)
Batch2_Sample3$Genotype <- c("Sftpc:Kras:mTmG")
Batch2_Sample3$Disease_Status <- c("LUAD")
Batch2_Sample3$Cell_of_Origin <- c("AT2-derived LUAD")
Batch2_Sample3$Mouse_ID <- c("S:K:TG008")
Batch2_Sample3$Batch <- c("Batch2")
prediction23 <- read.delim("your/path/here/azimuth_pred (3).tsv", row.names = 1)
Batch2_Sample3 <- AddMetaData(object = Batch2_Sample3, metadata = prediction23)

Batch2_Sample4 <- CreateSeuratObject(counts = Batch2_Sample4, project = "AT1-LUAD", min.cells = 3, min.features = 200)
Batch2_Sample4$Genotype <- c("Sftpc:Kras:mTmG")
Batch2_Sample4$Disease_Status <- c("LUAD")
Batch2_Sample4$Cell_of_Origin <- c("AT2-derived LUAD")
Batch2_Sample4$Mouse_ID <- c("S:K:TG010")
Batch2_Sample4$Batch <- c("Batch2")
prediction24 <- read.delim("your/path/here/azimuth_pred (4).tsv", row.names = 1)
Batch2_Sample4 <- AddMetaData(object = Batch2_Sample4, metadata = prediction24)

Batch2_Sample5 <- CreateSeuratObject(counts = Batch2_Sample5, project = "AT1-LUAD", min.cells = 3, min.features = 200)
Batch2_Sample5$Genotype <- c("Gramd2:Kras:mTmG")
Batch2_Sample5$Disease_Status <- c("Healthy")
Batch2_Sample5$Cell_of_Origin <- c("Vehicle_Controls")
Batch2_Sample5$Mouse_ID <- c("G:K:TG014")
Batch2_Sample5$Batch <- c("Batch2")
prediction25 <- read.delim("your/path/here/azimuth_pred (5).tsv", row.names = 1)
Batch2_Sample5 <- AddMetaData(object = Batch2_Sample5, metadata = prediction25)

Batch2_Sample6 <- CreateSeuratObject(counts = Batch2_Sample6, project = "AT1-LUAD", min.cells = 3, min.features = 200)
Batch2_Sample6$Genotype <- c("Gramd2:Kras:mTmG")
Batch2_Sample6$Disease_Status <- c("LUAD")
Batch2_Sample6$Cell_of_Origin <- c("AT1-derived LUAD")
Batch2_Sample6$Mouse_ID <- c("G:K:TG011")
Batch2_Sample6$Batch <- c("Batch2")
prediction26 <- read.delim("your/path/here/azimuth_pred (6).tsv", row.names = 1)
Batch2_Sample6 <- AddMetaData(object = Batch2_Sample6, metadata = prediction26)

#merge 6 Seurat objects into one
AT1LUAD_MergedBatch2 <- merge(Batch2_Sample1, y = c(Batch2_Sample2, Batch2_Sample3, Batch2_Sample4, Batch2_Sample5, Batch2_Sample6),
                              add.cell.ids = ls()[9:14],
                              project = "AT1-LUAD")

#save
saveRDS(AT1LUAD_MergedBatch2, file = "your/path/here/AT1LUAD_MergedBatch2.RDS")


################### Standard data processing for BATCH1 ################### 
#Standard pre-processing workflow:
#1. Quality control and selecting cells for further analysis
#2. Data normalization (NormalizeData)
#3. Identification of highly variable features (feature selection: FindVariableFeatures)
#4. Data scaling (ScaleData)
##1.1. view QC metrics
AT1LUAD_MergedBatch1 <- readRDS("your/path/here/AT1LUAD_MergedBatch1.RDS")
#inspection
View(AT1LUAD_MergedBatch1@meta.data)
range(AT1LUAD_MergedBatch1$nFeature_RNA)
range(AT1LUAD_MergedBatch1$nCount_RNA)
#store mitochondrial percentage in object meta data
AT1LUAD_MergedBatch1 <- PercentageFeatureSet(AT1LUAD_MergedBatch1, pattern = "^mt-", col.name = "percent.mt")
range(AT1LUAD_MergedBatch1$percent.mt)

##1.2. selecting cells for further analysis
# visualize QC metrics as a violin plot
VlnPlot(AT1LUAD_MergedBatch1, features = c("nFeature_RNA", "nCount_RNA","percent.mt"), group.by = c("Disease_Status"), ncol = 3,
        log = TRUE, pt.size = 2) + NoLegend()
VlnPlot(AT1LUAD_MergedBatch1, features = c("nFeature_RNA", "nCount_RNA","percent.mt"), group.by = c("Genotype"), ncol = 3,
        log = TRUE, pt.size = 0) + NoLegend()
VlnPlot(AT1LUAD_MergedBatch1, features = c("nFeature_RNA", "nCount_RNA","percent.mt"), group.by = c("Mouse_ID"), ncol = 3,
        log = TRUE, pt.size = 0) + NoLegend()
# remove low-quality samples
AT1LUAD_MergedBatch1 <- subset(AT1LUAD_MergedBatch1, subset = nFeature_RNA > 100 & nFeature_RNA < 10000 & nCount_RNA < 90000 &
                                 percent.mt <= 20)
# visualize QC metrics as a violin plot after QC
VlnPlot(AT1LUAD_MergedBatch1, features = c("nFeature_RNA", "nCount_RNA","percent.mt"), group.by = c("Mouse_ID"), ncol = 3,
        log = TRUE, pt.size = 0) + NoLegend()


##2. Data normalization
# following data transformation step is required if you are using Seurat v5 -- convert Seurat object of v5 to v3
AT1LUAD_MergedBatch1[['RNA3']]<-as(object=AT1LUAD_MergedBatch1[['RNA']], Class='Assay') #create a v3 Assay
DefaultAssay(AT1LUAD_MergedBatch1)<-'RNA3'                                    # set v3 Assay as the default
AT1LUAD_MergedBatch1[['RNA']]<-NULL                                           # delete the v5 Assay
AT1LUAD_MergedBatch1<-RenameAssays(AT1LUAD_MergedBatch1, RNA3='RNA')                    # rename the v3 Assay as 'RNA'

# normalization
AT1LUAD_MergedBatch1@assays[["RNA"]]@data@x
AT1LUAD_MergedBatch1 <- NormalizeData(AT1LUAD_MergedBatch1, normalization.method = "LogNormalize", scale.factor = 10000)
AT1LUAD_MergedBatch1@assays[["RNA"]]@data@x

##3. Identification of highly variable features
AT1LUAD_MergedBatch1

AT1LUAD_MergedBatch1 <- FindVariableFeatures(AT1LUAD_MergedBatch1, selection.method = "vst", nfeatures = 2000) #vst = variance stablizing transformatin

# Top features and featurePlot
AT1LUAD_MergedBatch1@assays[["RNA"]]@var.features #first 1000 variable features
top10VariableFeatures <- head(VariableFeatures(AT1LUAD_MergedBatch1), 10)
plot1 <- VariableFeaturePlot(AT1LUAD_MergedBatch1)
plot2 <- LabelPoints(plot = plot1, points = top10VariableFeatures, repel = TRUE)
plot1 + plot2

##4. Data scaling
AT1LUAD_MergedBatch1 <- ScaleData(object = AT1LUAD_MergedBatch1)

AT1LUAD_MergedBatch1@assays[["RNA"]]@scale.data
View(AT1LUAD_MergedBatch1@commands)

# Perform PCA, linear dimensional reduction, on the scaled data (linear dimensional reduction)
AT1LUAD_MergedBatch1 <- RunPCA(AT1LUAD_MergedBatch1, features = VariableFeatures(object = AT1LUAD_MergedBatch1)) # generate 5 PCs, first 30 features in each PC


# Examine and visualize PCA results a few different ways: DimPlot(); VizDimReduction(); DimHeatmap()
DimPlot(AT1LUAD_MergedBatch1, reduction = "pca", dims = c(1,2))
DimPlot(AT1LUAD_MergedBatch1, reduction = "pca", dims = c(1,10))
DimPlot(AT1LUAD_MergedBatch1, reduction = "pca", dims = c(1,50))

# Determine the "dimensional" of the dataset
AT1LUAD_MergedBatch1 <- JackStraw(AT1LUAD_MergedBatch1, num.replicate = 100, dims = 20)
AT1LUAD_MergedBatch1 <- ScoreJackStraw(AT1LUAD_MergedBatch1, dims = 1:20)
JackStrawPlot(AT1LUAD_MergedBatch1, dims = 1:20)

# more direct way: ElbowPlot()
ElbowPlot(AT1LUAD_MergedBatch1) #default = 20 PC
ElbowPlot(AT1LUAD_MergedBatch1, ndims = 50, reduction = "pca") #more PCs, more precise 

DimHeatmap(AT1LUAD_MergedBatch1, dims = 1:15, cells = 500, balanced = TRUE)

# Cluster cells
AT1LUAD_MergedBatch1 <- FindNeighbors(AT1LUAD_MergedBatch1, dims = 1:16)
AT1LUAD_MergedBatch1 <- FindClusters(AT1LUAD_MergedBatch1, resolution = 0.5) #16 clusters


#Run non-linear dimensional reduction (UMAP/tSNE)
AT1LUAD_MergedBatch1 <- RunUMAP(AT1LUAD_MergedBatch1, dims = 1:16)
AT1LUAD_MergedBatch1$seurat_clusters <- as.factor(as.numeric(as.character(AT1LUAD_MergedBatch1$seurat_clusters)) +1)
DimPlot(AT1LUAD_MergedBatch1, reduction = "umap", label = FALSE, repel = TRUE)

saveRDS(AT1LUAD_MergedBatch1, file = "your/path/here/AT1LUAD_MergedBatch1.RDS")


################### for BATCH2 ################### 
#Standard pre-processing workflow:
#1. Quality control and selecting cells for further analysis
#2. Data normalization (NormalizeData)
#3. Identification of highly variable features (feature selection: FindVariableFeatures)
#4. Data scaling (ScaleData)
##1.1. view QC metrics
AT1LUAD_MergedBatch2 <- readRDS("your/path/here/AT1LUAD_MergedBatch2.RDS")

View(AT1LUAD_MergedBatch2@meta.data)
range(AT1LUAD_MergedBatch2$nFeature_RNA)
range(AT1LUAD_MergedBatch2$nCount_RNA)

# store mitochondrial percentage in object meta data
AT1LUAD_MergedBatch2 <- PercentageFeatureSet(AT1LUAD_MergedBatch2, pattern = "^mt-", col.name = "percent.mt")
range(AT1LUAD_MergedBatch2$percent.mt)

##1.2. selecting cells for further analysis
# visualize QC metrics as a violin plot
VlnPlot(AT1LUAD_MergedBatch2, features = c("nFeature_RNA", "nCount_RNA","percent.mt"), group.by = c("Disease_Status"), ncol = 3,
        log = TRUE, pt.size = 2) + NoLegend()
VlnPlot(AT1LUAD_MergedBatch2, features = c("nFeature_RNA", "nCount_RNA","percent.mt"), group.by = c("Genotype"), ncol = 3,
        log = TRUE, pt.size = 0) + NoLegend()
VlnPlot(AT1LUAD_MergedBatch2, features = c("nFeature_RNA", "nCount_RNA","percent.mt"), group.by = c("Mouse_ID"), ncol = 3,
        log = TRUE, pt.size = 0) + NoLegend()

AT1LUAD_MergedBatch2

# remove low-quality samples
AT1LUAD_MergedBatch2 <- subset(AT1LUAD_MergedBatch2, subset = nFeature_RNA > 100 & nFeature_RNA < 10000 & nCount_RNA < 90000 &
                                 percent.mt <= 20)
VlnPlot(AT1LUAD_MergedBatch2, features = c("nFeature_RNA", "nCount_RNA","percent.mt"), group.by = c("Mouse_ID"), ncol = 3,
        log = TRUE, pt.size = 0) + NoLegend()


##2. Data normalization
# following data transformation step is required if you are using Seurat v5 -- convert Seurat object of v5 to v3
AT1LUAD_MergedBatch2[['RNA3']]<-as(object=AT1LUAD_MergedBatch2[['RNA']], Class='Assay') #create a v3 Assay
DefaultAssay(AT1LUAD_MergedBatch2)<-'RNA3'                                    # set v3 Assay as the default
AT1LUAD_MergedBatch2[['RNA']]<-NULL                                           # delete the v5 Assay
AT1LUAD_MergedBatch2<-RenameAssays(AT1LUAD_MergedBatch2, RNA3='RNA')                    # rename the v3 Assay as 'RNA'
AT1LUAD_MergedBatch2@assays[["RNA"]]@data@x

AT1LUAD_MergedBatch2 <- NormalizeData(AT1LUAD_MergedBatch2, normalization.method = "LogNormalize", scale.factor = 10000)

AT1LUAD_MergedBatch2@assays[["RNA"]]@data@x

##3. Identification of highly variable features
AT1LUAD_MergedBatch2

AT1LUAD_MergedBatch2 <- FindVariableFeatures(AT1LUAD_MergedBatch2, selection.method = "vst", nfeatures = 2000) #vst = variance stablizing transformatin

#Top features and featurePlot
AT1LUAD_MergedBatch2@assays[["RNA"]]@var.features #first 1000 variable features
top10VariableFeatures <- head(VariableFeatures(AT1LUAD_MergedBatch2), 10)
plot1 <- VariableFeaturePlot(AT1LUAD_MergedBatch2)
plot2 <- LabelPoints(plot = plot1, points = top10VariableFeatures, repel = TRUE)
plot1 + plot2


##4. Data scaling
AT1LUAD_MergedBatch2 <- ScaleData(object = AT1LUAD_MergedBatch2)

AT1LUAD_MergedBatch2@assays[["RNA"]]$scale.data
View(AT1LUAD_MergedBatch2@commands)

### Perform PCA, linear dimensional reduction, on the scaled data (linear dimensional reduction)
AT1LUAD_MergedBatch2 <- RunPCA(AT1LUAD_MergedBatch2, features = VariableFeatures(object = AT1LUAD_MergedBatch2))
# generate 5 PCs, first 30 features in each PC

#Examine and visualize PCA results a few different ways: DimPlot(); VizDimReduction(); DimHeatmap()
DimPlot(AT1LUAD_MergedBatch2, reduction = "pca", dims = c(1,2))
DimPlot(AT1LUAD_MergedBatch2, reduction = "pca", dims = c(1,10))
DimPlot(AT1LUAD_MergedBatch2, reduction = "pca", dims = c(1,50))

#Determine the "dimensional" of the dataset
AT1LUAD_MergedBatch2 <- JackStraw(AT1LUAD_MergedBatch2, num.replicate = 100, dims = 20)
AT1LUAD_MergedBatch2 <- ScoreJackStraw(AT1LUAD_MergedBatch2, dims = 1:20)
JackStrawPlot(AT1LUAD_MergedBatch2, dims = 1:20)

#more direct way: ElbowPlot()
ElbowPlot(AT1LUAD_MergedBatch2) #default = 20 PC
ElbowPlot(AT1LUAD_MergedBatch2, ndims = 50, reduction = "pca") #more PCs, more precise 

DimHeatmap(AT1LUAD_MergedBatch2, dims = 1:20, cells = 500, balanced = TRUE)

#Cluster cells
AT1LUAD_MergedBatch2 <- FindNeighbors(AT1LUAD_MergedBatch2, dims = 1:20)
AT1LUAD_MergedBatch2 <- FindClusters(AT1LUAD_MergedBatch2, resolution = 0.2) #17 clusters

#Run non-linear dimensional reduction (UMAP/tSNE)
AT1LUAD_MergedBatch2 <- RunUMAP(AT1LUAD_MergedBatch2, dims = 1:20)
AT1LUAD_MergedBatch2$seurat_clusters <- as.factor(as.numeric(as.character(AT1LUAD_MergedBatch2$seurat_clusters)) +1)
DimPlot(AT1LUAD_MergedBatch2, reduction = "umap", label = FALSE, repel = TRUE)

saveRDS(AT1LUAD_MergedBatch2, file = "your/file/path/AT1LUAD_MergedBatch2.RDS")


################################## to integrate data from Batch1 and Batch2 (Harmony) ################################## 
AT1LUAD_BatchCombined <- merge(AT1LUAD_MergedBatch1, y = AT1LUAD_MergedBatch2, add.cell.ids = c("Batch1", "Batch2"), project = "AT1-LUAD")

#Standard pre-processing workflow:
#1. Quality control and selecting cells for further analysis
#2. Data normalization (NormalizeData)
#3. Identification of highly variable features (feature selection: FindVariableFeatures)
#4. Data scaling (ScaleData)
##1.1. view QC metrics
View(AT1LUAD_BatchCombined@meta.data)
range(AT1LUAD_BatchCombined$nFeature_RNA)
range(AT1LUAD_BatchCombined$nCount_RNA)


#store mitochondrial percentage in object meta data
AT1LUAD_BatchCombined <- PercentageFeatureSet(AT1LUAD_BatchCombined, pattern = "^mt-", col.name = "percent.mt")
range(AT1LUAD_BatchCombined$percent.mt)
# [1]  0.00000 20

##1.2. selecting cells for further analysis
# visualize QC metrics as a violin plot
# Supplemental figure 1a
VlnPlot(AT1LUAD_BatchCombined, features = c("nFeature_RNA", "nCount_RNA","percent.mt"), group.by = c("Mouse_ID"), ncol = 3,
        log = TRUE, pt.size = 0) + NoLegend()
# remove low-quality samples
AT1LUAD_BatchCombined <- subset(AT1LUAD_BatchCombined, subset = nFeature_RNA > 100 & nFeature_RNA < 10000 & nCount_RNA < 100000 &
                                  percent.mt <= 20)
VlnPlot(AT1LUAD_BatchCombined, features = c("nFeature_RNA", "nCount_RNA","percent.mt"), group.by = c("Mouse_ID"), ncol = 3,
        log = TRUE, pt.size = 0) + NoLegend()

##2. Data normalization
AT1LUAD_BatchCombined <- NormalizeData(AT1LUAD_BatchCombined, normalization.method = "LogNormalize", scale.factor = 10000)
##3. Identification of highly variable features
AT1LUAD_BatchCombined <- FindVariableFeatures(AT1LUAD_BatchCombined, selection.method = "vst", nfeatures = 2000) #vst = variance stablizing transformatin
#Top features and featurePlot
AT1LUAD_BatchCombined@assays[["RNA"]]@var.features #first 1000 variable features
top10VariableFeatures <- head(VariableFeatures(AT1LUAD_BatchCombined), 10)
##4. Data scaling
AT1LUAD_BatchCombined <- ScaleData(object = AT1LUAD_BatchCombined) #2000 identified variable features
###Perform PCA, linear dimensional reduction, on the scaled data (linear dimensional reduction)
AT1LUAD_BatchCombined <- RunPCA(AT1LUAD_BatchCombined, features = VariableFeatures(object = AT1LUAD_BatchCombined))
#Examine and visualize PCA results a few different ways: DimPlot(); VizDimReduction(); DimHeatmap()
DimPlot(AT1LUAD_BatchCombined, reduction = "harmony", dims = c(1,2))
DimPlot(AT1LUAD_BatchCombined, reduction = "harmony", dims = c(1,10))
DimPlot(AT1LUAD_BatchCombined, reduction = "harmony", dims = c(1,50))
AT1LUAD_BatchCombined <- JackStraw(AT1LUAD_BatchCombined, num.replicate = 100, dims = 20)
AT1LUAD_BatchCombined <- ScoreJackStraw(AT1LUAD_BatchCombined, dims = 1:20)
# Supplemental figure 1b
JackStrawPlot(AT1LUAD_BatchCombined, dims = 1:20)
#more direct way: ElbowPlot()
ElbowPlot(AT1LUAD_BatchCombined) #default = 20 PC
# Supplemental figure 1c
ElbowPlot(AT1LUAD_BatchCombined, ndims = 50, reduction = "pca") #more PCs, more precise 
# Supplemental figure 1d
DimHeatmap(AT1LUAD_BatchCombined, dims = 1:20, cells = 500, balanced = TRUE)
#Cluster cells
AT1LUAD_BatchCombined <- FindNeighbors(AT1LUAD_BatchCombined, dims = 1:16)
AT1LUAD_BatchCombined <- FindClusters(AT1LUAD_BatchCombined, resolution = 0.2) #20 clusters

### run Harmoney to perform data integration (for batch effect reduction)-----------------
AT1LUAD_BatchIntegrated <- AT1LUAD_BatchCombined %>%
  RunHarmony(group.by.vars = 'Batch', plot_covergence = FALSE)

AT1LUAD_BatchIntegrated@reductions

AT1LUAD_BatchIntegrated.embed <- Embeddings(AT1LUAD_BatchIntegrated, "harmony")
AT1LUAD_BatchIntegrated.embed[1:10,1:10]

# Do UMAP and clustering using ** Harmony embeddings instead of PCA **
AT1LUAD_BatchIntegrated <- AT1LUAD_BatchIntegrated %>%
  RunUMAP(reduction = 'harmony', dims = 1:20) %>%
  FindNeighbors(reduction = 'harmony', dims = 1:20) %>%
  FindClusters(resolution = 0.3) #16 clusters

set.seed(1234)
# Visualize
head(AT1LUAD_BatchIntegrated)

# Figure 1b-1e
AT1LUAD_BatchIntegrated$Cell_of_Origin <- factor(AT1LUAD_BatchIntegrated$Cell_of_Origin, levels = c("Neg_Controls", "Vehicle_Controls", "Spontaneous_AT2-derived LUAD", "AT2-derived LUAD", "AT1-derived LUAD"))
AT1LUAD_BatchIntegrated$seurat_clusters <- as.factor(as.numeric(as.character(AT1LUAD_BatchIntegrated$seurat_clusters)) +1)
DimPlot(AT1LUAD_BatchIntegrated, reduction = "umap", label = FALSE, group.by = "seurat_clusters", repel = TRUE)
DimPlot(AT1LUAD_BatchIntegrated, reduction = "umap", label = FALSE, group.by = "Disease_Status", cols = c("#c0ced6", "#DF1528", "#7262ac"))
DimPlot(AT1LUAD_BatchIntegrated, reduction = "umap", label = FALSE, group.by = "predicted.lineage_level1")
DimPlot(AT1LUAD_BatchIntegrated, reduction = "umap", label = FALSE, group.by = "predicted.lineage_level2", 
        cols=c("lightgrey", "lightgrey", "lightgrey", "lightgrey", "#40a4f2", "#000099", "#530de8", "lightgrey", "lightgrey", "lightgrey"))

# Supplemental figure 1e-1g
DimPlot(AT1LUAD_BatchIntegrated, reduction = "umap", label = FALSE, group.by = "Batch")
DimPlot(AT1LUAD_BatchIntegrated, reduction = "umap", label = FALSE, group.by = "Mouse_ID", repel = TRUE)
DimPlot(AT1LUAD_BatchIntegrated, reduction = "umap", label = FALSE, group.by = c("Cell_of_Origin"), cols = c("#0c0d0c", "#5f5f5f", "#7262ac", "#00a0e3",  "#e31e24"))

### Primer of feature plot and group by original identification (= "project" when loading the data)
# Supplementalfigure 1h
FeaturePlot(AT1LUAD_BatchIntegrated, features = c("Foxj1", "Dnah5"), blend = TRUE,
            cols = c("lightgrey", "blue", "red"))

#save
saveRDS(AT1LUAD_BatchIntegrated, file = "your/file/path/LUAD_BatchIntegrated.RDS")


########## Percent stacked barchart #############
T1 <- table(AT1LUAD_BatchIntegrated$predicted.lineage_level2, AT1LUAD_BatchIntegrated$Cell_of_Origin)
df <- data.frame(T1, row.names = NULL)
write.csv(df, file = "your/file/path/Percent stacked barchart.csv")

library(readxl)
Percent <- read_excel("your/file/path/Supplemental Table1_Overall cell compostion.xlsx")
Percent <- data.frame(Percent, row.names = NULL)
Percent$Cell_of_Origin <- factor(Percent$Cell_of_Origin, levels = c("Neg_Controls", "Vehicle_Controls", "Spontaneous_AT2-derived LUAD", "AT2-derived LUAD", "AT1-derived LUAD"))
Percent$Celltypes <- factor(Percent$Celltypes, levels = c("HSPCs", "Monocytes", "MDSCs", "Proliferating Myeloid Cells", "Immature Macrophages", "Macrophages", "DCs"))

# Figure 1f
ggplot(Percent, aes(fill=Lineages, y=Frequency, x=Cell_of_Origin)) + 
  geom_bar(position="fill", stat="identity") +
  ggtitle("Percentage of Cells") +
  theme(
    panel.background = element_rect(fill = "transparent", colour = NA), # Remove background
    plot.background = element_rect(fill = "transparent", colour = NA)   # Remove background from the plot area
  ) +
  RotatedAxis()

#################################################################################################################################################
##################################################### FIGURE 2,3,4 & Supplemental FIGURE2,4 #####################################################
#################################################################################################################################################
library(patchwork)
library(Seurat)
library(ggplot2)
library(dplyr)
library(biovizBase)
library(hdf5r)
library(tidyverse)
library(fgsea)

# load the previoysly saved RDS file
AT1LUAD_BatchIntegrated <- readRDS("your/file/path/LUAD_BatchIntegrated.RDS")

############# Extract Myeloid cells ############# 
DimPlot(AT1LUAD_BatchIntegrated, reduction = "umap", label = FALSE, group.by = "predicted.celltype_level3")
AM <- subset(x = AT1LUAD_BatchIntegrated, subset = (predicted.celltype_level3 == "AM"))
IM <- subset(x = AT1LUAD_BatchIntegrated, subset = (predicted.celltype_level3 == "IM"))
iMON <- subset(x = AT1LUAD_BatchIntegrated, subset = (predicted.celltype_level3 == "iMON"))
cDC1 <- subset(x = AT1LUAD_BatchIntegrated, subset = (predicted.celltype_level3 == "cDC1"))
cDC2 <- subset(x = AT1LUAD_BatchIntegrated, subset = (predicted.celltype_level3 == "cDC2"))
maDC <- subset(x = AT1LUAD_BatchIntegrated, subset = (predicted.celltype_level3 == "maDC"))

Myeloid <- merge(x = AM, y = list(IM, iMON, cDC1, cDC2, maDC), 
                           add.cell.ids = c("AM", "IM", "iMON", "cDC1", "cDC2", "maDC"))


### remove unspecific cells ###
# repeat the following filtration until the sample size remains a constant value (~366.9 MB)
Myeloid <- subset(x = Myeloid, subset = (predicted.lineage_level1 == c("Endothelial",
                                                                       "Epithelial",
                                                                       "Mesenchymal")), invert = TRUE)

Myeloid <- subset(x = Myeloid, subset = (predicted.lineage_level2 == c("Endothelial_Capillary",
                                                                        "Endothelial_Main",
                                                                        "Epithelial_Airway",
                                                                        "Epithelial_Alveolar",
                                                                        "Immune_InnateLymphoid",
                                                                        "Immune_Lymphoid",
                                                                        "Mesenchymal_Alveolar",
                                                                        "Mesenchymal_Airway/Vascular",
                                                                        "Mesenchymal_Pleura")), invert = TRUE)


############################ Standard Seurat procedures for Myeloid_Annotated ############################
Myeloid[["RNA"]] <- JoinLayers(Myeloid[["RNA"]])
Myeloid <- NormalizeData(Myeloid)
Myeloid <- FindVariableFeatures(Myeloid, selection.method = "vst", nfeatures = 2000)
Myeloid <- ScaleData(object = Myeloid)
Myeloid <- RunPCA(Myeloid, features = VariableFeatures(object = Myeloid))
ElbowPlot(Myeloid) 
ElbowPlot(Myeloid, ndims = 50, reduction = "pca") #more PCs, more precise 
DimHeatmap(Myeloid, dims = 1:25, cells = 500, balanced = TRUE)
Myeloid <- Myeloid %>%
  FindNeighbors(reduction = 'pca', dims = 1:21) %>%
  FindClusters(resolution = 0.8) %>% #17 clusters
  RunUMAP(reduction = 'pca', dims = 1:21) 

### run Harmony for Myeloid cells -----------------
Myeloid <- Myeloid %>%
  RunHarmony(group.by.vars = 'Batch', plot_covergence = FALSE)

Myeloid@reductions

Myeloid.embed <- Embeddings(Myeloid, "harmony")
Myeloid.embed[1:10,1:10]

# Do UMAP and clustering using ** Harmony embeddings instead of PCA **
Myeloid <- JackStraw(Myeloid, num.replicate = 100, dims = 20)  
Myeloid <- ScoreJackStraw(Myeloid, dims = 1:20)     
JackStrawPlot(Myeloid, dims = 1:20)
ElbowPlot(Myeloid) 
Myeloid <- Myeloid %>%
  FindNeighbors(reduction = 'harmony', dims = 1:21) %>%
  FindClusters(resolution = 1.4) %>%
  RunUMAP(reduction = 'harmony', dims = 1:21) #22 clusters

# save
saveRDS(Myeloid, file = "your/file/path/Myeloid.RDS")

### Supplemental figure 2a ### 
### Verify the resolution and clustering number via using Clustree
library(clustree)
# Extract clustering information and prepare data for clustree
clustering_data <- Myeloid@meta.data[, grepl("RNA_snn_res", names(Myeloid@meta.data))]
# Adding cell names as a new column
clustering_data$cell <- rownames(clustering_data)
# Removing rows with missing values in clustering data
clustering_data_clean <- na.omit(clustering_data)
# Now, use clustree on the cleaned data
clustree(clustering_data_clean, prefix = "RNA_snn_res.")


Myeloid$Cell_of_Origin <- factor(Myeloid$Cell_of_Origin, levels = c("Neg_Controls", "Vehicle_Controls", "Spontaneous_AT2-derived LUAD", "AT2-derived LUAD", "AT1-derived LUAD"))
Myeloid$seurat_clusters <- as.factor(as.numeric(as.character(Myeloid$seurat_clusters)) +1)
# Supplemental figure 2b #
DimPlot(Myeloid, reduction = "umap", label = FALSE, group.by = "seurat_clusters", repel = TRUE)
# Supplemental figure 2c #
DimPlot(Myeloid, reduction = "umap", label = FALSE, group.by = "predicted.celltype_level3")
# figure 3a #
DimPlot(object = Myeloid, 
        reduction = "umap", 
        group.by = "Cell_of_Origin",
        cols = c("#0c0d0c", "#5f5f5f", "#7262ac", "#00a0e3",  "#e31e24"),
        label = F,
        repel = TRUE)


### Try to use SingleR to refine cell type annotation (from other single-cell reference)
library(SingleR)
library(celldex)
library(SingleCellExperiment)
ref_use <- celldex::ImmGenData()

results1 <- SingleR(test = as.SingleCellExperiment(Myeloid), ref = ref_use, labels = ref_use$label.fine)
results2 <- SingleR(test = as.SingleCellExperiment(Myeloid), ref = ref_use, labels = ref_use$label.main)

Myeloid$singlr_labels <- results2$labels

# Supplemental figure 2d #
P1 <- DimPlot(Myeloid, group.by = "singlr_labels", reduction = "umap", label = TRUE, label.size = 3) +  
  theme(legend.key.size = unit(0.5, 'cm'),  legend.key.height = unit(0.5, 'cm'), 
        legend.key.width = unit(0.5, 'cm'), legend.title = element_text(size=10), 
        legend.text = element_text(size=10))
print(P1)


######################## Identify myeloid cell subtypes of Myeloid clusters ######################## 
### to obtain top 10 markers in each cluster ###
library(presto)
Myeloid_Annotated.Markers <- FindAllMarkers(Myeloid_Annotated, only.pos = TRUE, logfc.threshold = 0.25, verbose = FALSE)
write.csv(Myeloid_Annotated.Markers, 
          file="/Users/m.y./Documents/R/R_working directory/AT1LUAD_scRNA/Immune/Myeloid_Annotated.Markers_re.CSV")

Mye_top10 <- Myeloid_Annotated.Markers %>%
  group_by(cluster) %>%
  top_n(n=10, 
        wt = avg_log2FC)
write.csv(Mye_top10, 
          file="/Users/m.y./Documents/R/R_working directory/AT1LUAD_scRNA/Immune/Myeloid_Annotated.MarkersTOP10_re.CSV")

# rename the clusters
Myeloid_Annotated <- RenameIdents(object = Myeloid, 
                                  "0" = "MDSCs",
                                  "1" = "Macrophages",
                                  "2" = "Macrophages",
                                  "3" = "Macrophages",
                                  "4" = "TAMs",
                                  "5" = "Macrophages",
                                  "6" = "Macrophages",
                                  "7" = "Macrophages",
                                  "8" = "Macrophages",
                                  "9" = "Macrophages",
                                  "10" = "Proliferating Myeloid_Annotated Cells",
                                  "11" = "HSPCs",
                                  "12" = "Proliferating Myeloid_Annotated Cells",
                                  "13" = "Macrophages",
                                  "14" = "DCs",
                                  "15" = "Macrophages",
                                  "16" = "iMONs",
                                  "17" = "Macrophages",
                                  "18" = "Macrophages",
                                  "19" = "DCs",
                                  "20" = "HSPCs",
                                  "21" = "Granulocytes")

# create a new metadata column for myeloid cell subtype information
Myeloid_Annotated$Celltypes<-Idents(Myeloid_Annotated)
view (Myeloid_Annotated@meta.data)

# combine group ID & Myeloid cell subtypes
Myeloid_Annotated$NEW_celltype <- paste(Myeloid_Annotated$Cell_of_Origin, sep = "_", Idents(Myeloid_Annotated))
view (Myeloid_Annotated@meta.data)

# save
saveRDS(Myeloid_Annotated, file = "your/file/path/Myeloid_Annotated.RDS")

# Plot the UMAP
Myeloid_Annotated$Cell_of_Origin <- factor(Myeloid_Annotated$Cell_of_Origin, levels = c("Neg_Controls", "Vehicle_Controls", "Spontaneous_AT2-derived LUAD",  "AT2-derived LUAD", "AT1-derived LUAD"))
Myeloid_Annotated$Celltypes <- factor(Myeloid_Annotated$Celltypes, levels = c("HSPCs", "iMONs", "Proliferating Myeloid_Annotated Cells",  "MDSCs", "TAMs", "Macrophages", "DCs", "Granulocytes"))

# Figure 2a
DimPlot(object = Myeloid_Annotated, 
        reduction = "umap", 
        group.by = "Celltypes",
        cols = c("#e0271e", "#ff8ad7", "orange", "#ffff99", "#357337", "#2b9cd7", "#cfafd4", "#715ea9"),
        label = F,
        label.size = 4,
        repel = TRUE)


### Verify the manual cell annotation ###
#Features of Macrophages, MDSCs, ST-HSCs, Immature Macrophages, DCs, Proliferating mye, Monocytes, maDCs
Overall_Features <- c("Kit", "Ly6a", "Cd34",  # HSPCs
                      "Csf1r", "Itgam", "Cx3cr1", # iMONs
                      "Neil3", "Kif14", "Esco2", "Mki67",  #Prolif.myeloid
                      "Fcer1g", "Tyrobp", "Spi1", "Ly6c1",  # MDSCs
                      "Mmp8", "Mmp12", "Arg1","Chil3","Chil4", "Cd274", # TAMs
                      "Cd33", "Siglecf", "Mertk", "Itgax",  # Macrophages
                      "H2-Ab1", "H2-Aa", "H2-Eb1", "Cst3",  # DCs
                      "Csf3r", "Cd24a", "Trem1", "Ltf" # Granulocytes
)

# Figure 2b 
DotPlot(Myeloid_Annotated, features =  Overall_Features, 
        group.by = "Celltypes",
        scale = T, 
        col.min = 0,
        dot.scale = 7,
        cols = c("lightgrey", "blue")) + RotatedAxis()


########## Percent stacked barchart #############
T1 <- table(Myeloid_Annotated$Celltypes, Myeloid_Annotated$Cell_of_Origin)
df <- data.frame(T1, row.names = NULL)
write.csv(df, file = "your/file/path/Percent stacked barchart_Mye.csv")
T2 <- table(Myeloid_Annotated$Celltypes, Myeloid_Annotated$Mouse_ID)
df2 <- data.frame(T2, row.names = NULL)
write.csv(df2, file = "your/file/path/Percent stacked barchart_Mye_Mouse_ID.csv")

library(readxl)
Percent <- read_excel("your/file/path/Supplemental Table2_Myeloid cell composition.xlsx")
Mye_Percent <- data.frame(Percent, row.names = NULL)
Mye_Percent$Cell_of_Origin <- factor(Mye_Percent$Cell_of_Origin, levels = c("Neg_Controls", "Vehicle_Controls", "Spontaneous_AT2-derived LUAD", "AT2-derived LUAD", "AT1-derived LUAD"))
Mye_Percent$Celltypes <- factor(Mye_Percent$Celltypes, levels = c("HSPCs", "iMONs", "Proliferating Myeloid Cells", "MDSCs", "TAMs", "Macrophages", "DCs", "Granulocytes"))

colors <- c("#e0271e", "#ff8ad7", "orange", "#ffff99", "#357337", "#2b9cd7", "#cfafd4", "#715ea9")

# Figure 2c #
ggplot(Mye_Percent, aes(fill=Celltypes, y=Proportion, x=Cell_of_Origin)) + 
  geom_bar(position="fill", stat="identity") +
  scale_fill_manual(values = colors) +
  ggtitle("Percentage of Myeloid_Annotated Cells")+
  theme(
    panel.background = element_rect(fill = "transparent", colour = NA), # Remove background
    plot.background = element_rect(fill = "transparent", colour = NA)   # Remove background from the plot area
  ) +
  RotatedAxis()


### to generate the bar plots for showing cell proportions of iMONs, MDSC, TAMs, and Macrophages respectively ###
data_summary <- function(data, varname, groupnames){
  require(plyr)
  summary_func <- function(x, col){
    c(mean = mean(x[[col]], na.rm=TRUE),
      sd = sd(x[[col]], na.rm=TRUE))
  }
  data_sum<-ddply(data, groupnames, .fun=summary_func,
                  varname)
  data_sum <- rename(data_sum, c("mean" = varname))
  return(data_sum)
}

### iMONs ###
data_iMON <- read_excel("your/file/path/Supplemental Table3_Myeloid cell compostion per mouse ID.xlsx", sheet = 2)
data_iMON$Group_ID <- factor(data_iMON$Group_ID, levels = c("Neg_controls", "Vehicle_controls", "Sp_AT2-LUAD", "AT2-LUAD", "AT1-LUAD"))

data_iMON <- data_summary(data_iMON, varname="Proportion", 
                          groupnames=c("Group_ID"))
# Convert dose to a factor variable
data_iMON$Group_ID=as.factor(data_iMON$Group_ID)
head(data_iMON)

# Default bar plot
p <- ggplot(data_iMON, aes(x=Group_ID, y=Proportion, fill=Group_ID)) + 
  geom_bar(stat="identity", color="black", 
           position=position_dodge()) +
  geom_errorbar(aes(ymin=pmax(0, Proportion-sd), ymax=Proportion+sd), width=.2,
                position=position_dodge(.9)) 

# Print the initial plot
print(p)

# Finished bar plot with additional customizations
# Figure d #
p + labs(title="iMON", x="Group_ID", y="Proportion") +
  theme_classic() +
  scale_fill_manual(values=c("Neg_controls" = "#0c0d0c", 
                             "Vehicle_controls" = "#5f5f5f", 
                             "Sp_AT2-LUAD" = "#7262ac", 
                             "AT2-LUAD" = "#00a0e3", 
                             "AT1-LUAD" = "#e31e24"))

### MDSCs ###
data_MDSCs <- read_excel("your/file/path/Supplemental Table3_Myeloid cell compostion per mouse ID.xlsx", sheet = 3)
data_MDSCs$Group_ID <- factor(data_iMON$Group_ID, levels = c("Neg_controls", "Vehicle_controls", "Sp_AT2-LUAD", "AT2-LUAD", "AT1-LUAD"))

data_MDSCs <- data_summary(data_MDSCs, varname="Proportion", 
                          groupnames=c("Group_ID"))
# Convert dose to a factor variable
data_MDSCs$Group_ID=as.factor(data_MDSCs$Group_ID)
head(data_MDSCs)

# Default bar plot
p <- ggplot(data_MDSCs, aes(x=Group_ID, y=Proportion, fill=Group_ID)) + 
  geom_bar(stat="identity", color="black", 
           position=position_dodge()) +
  geom_errorbar(aes(ymin=pmax(0, Proportion-sd), ymax=Proportion+sd), width=.2,
                position=position_dodge(.9)) 

# Print the initial plot
print(p)

# Finished bar plot with additional customizations
# Figure d #
p + labs(title="MDSCs", x="Group_ID", y="Proportion") +
  theme_classic() +
  scale_fill_manual(values=c("Neg_controls" = "#0c0d0c", 
                             "Vehicle_controls" = "#5f5f5f", 
                             "Sp_AT2-LUAD" = "#7262ac", 
                             "AT2-LUAD" = "#00a0e3", 
                             "AT1-LUAD" = "#e31e24"))

### TAMs ###
data_TAMs <- read_excel("your/file/path/Supplemental Table3_Myeloid cell compostion per mouse ID.xlsx", sheet = 4)
data_TAMs$Group_ID <- factor(data_TAMs$Group_ID, levels = c("Neg_controls", "Vehicle_controls", "Sp_AT2-LUAD", "AT2-LUAD", "AT1-LUAD"))

data_TAMs <- data_summary(data_TAMs, varname="Proportion", 
                           groupnames=c("Group_ID"))
# Convert dose to a factor variable
data_TAMs$Group_ID=as.factor(data_TAMs$Group_ID)
head(data_TAMs)

# Default bar plot
p <- ggplot(data_TAMs, aes(x=Group_ID, y=Proportion, fill=Group_ID)) + 
  geom_bar(stat="identity", color="black", 
           position=position_dodge()) +
  geom_errorbar(aes(ymin=pmax(0, Proportion-sd), ymax=Proportion+sd), width=.2,
                position=position_dodge(.9)) 

# Print the initial plot
print(p)

# Finished bar plot with additional customizations
# Figure d #
p + labs(title="TAMs", x="Group_ID", y="Proportion") +
  theme_classic() +
  scale_fill_manual(values=c("Neg_controls" = "#0c0d0c", 
                             "Vehicle_controls" = "#5f5f5f", 
                             "Sp_AT2-LUAD" = "#7262ac", 
                             "AT2-LUAD" = "#00a0e3", 
                             "AT1-LUAD" = "#e31e24"))

### Macrophages ###
data_Macrophages <- read_excel("your/file/path/Supplemental Table3_Myeloid cell compostion per mouse ID.xlsx", sheet = 5)
data_Macrophages$Group_ID <- factor(data_Macrophages$Group_ID, levels = c("Neg_controls", "Vehicle_controls", "Sp_AT2-LUAD", "AT2-LUAD", "AT1-LUAD"))

data_Macrophages <- data_summary(data_Macrophages, varname="Proportion", 
                          groupnames=c("Group_ID"))
# Convert dose to a factor variable
data_Macrophages$Group_ID=as.factor(data_Macrophages$Group_ID)
head(data_Macrophages)

# Default bar plot
p <- ggplot(data_Macrophages, aes(x=Group_ID, y=Proportion, fill=Group_ID)) + 
  geom_bar(stat="identity", color="black", 
           position=position_dodge()) +
  geom_errorbar(aes(ymin=pmax(0, Proportion-sd), ymax=Proportion+sd), width=.2,
                position=position_dodge(.9)) 

# Print the initial plot
print(p)

# Finished bar plot with additional customizations
# Figure d #
p + labs(title="Macrophages", x="Group_ID", y="Proportion") +
  theme_classic() +
  scale_fill_manual(values=c("Neg_controls" = "#0c0d0c", 
                             "Vehicle_controls" = "#5f5f5f", 
                             "Sp_AT2-LUAD" = "#7262ac", 
                             "AT2-LUAD" = "#00a0e3", 
                             "AT1-LUAD" = "#e31e24"))


# Figure 3b #
Anti_tumoral_TAM <- Plot_Density_Joint_Only(seurat_object = Myeloid_Annotated, 
                                            features = c("Cd86",'Il1a', "Cxcl2", "Ccl4", 'Tnf'),
                                            custom_palette = BlueAndRed())


library(Nebulosa)
library(scCustomize)
library(ggplot2) # v3.4.4

# to test enrichment of immunoreactive macrophages #
Anti_tumoral_MP <- Plot_Density_Joint_Only(seurat_object = Myeloid_Annotated, 
                                            features = c("Cd86",'Il1a', "Cxcl2", "Ccl4", 'Tnf'),
                                            custom_palette = BlueAndRed())
# transfer the anti_tumoral_scoresscore to Seurat object
Anti_tumoral_MP_scores <- Anti_tumoral_MP$data$feature
names(Anti_tumoral_MP_scores) <- rownames(Anti_tumoral_MP$data)
Myeloid_Annotated <- AddMetaData(Myeloid_Annotated, metadata = Anti_tumoral_MP_scores, col.name = "Anti_tumoral_MP_scores")
# Verify the addition
head(Myeloid_Annotated@meta.data)

# figure 3b #
FeaturePlot_scCustom(Myeloid_Annotated, features = "Anti_tumoral_MP_scores", na_cutoff = 1.0e-08)


# to test enrichment of immunosuppressive macrophages #
Pro_tumoral_MP <- Plot_Density_Joint_Only(seurat_object = Myeloid_Annotated, 
                                           features = c("Msr1", "Mrc1", "Chil3", "Arg1",  "Cd274"),
                                           custom_palette = BlueAndRed())
# transfer the anti_tumoral_scoresscore to Seurat object
Pro_tumoral_MP_scores <- Pro_tumoral_MP$data$feature
names(Pro_tumoral_MP_scores) <- rownames(Pro_tumoral_MP$data)
Myeloid_Annotated <- AddMetaData(Myeloid_Annotated, metadata = Pro_tumoral_MP_scores, col.name = "Pro_tumoral_TAM_score")
# Verify the addition
head(Myeloid_Annotated@meta.data)

# figure 4b #
FeaturePlot_scCustom(Myeloid_Annotated, features = "Pro_tumoral_TAM_score", na_cutoff = 5.0e-08)


# to test enrichment of MDSCs #
p_density_MDSC <- Plot_Density_Joint_Only(seurat_object = Myeloid_Annotated, 
                                       features = c("S100a8", "S100a9", "Wfdc17", "Cd36", "Cd38", "Cd84"),
                                       custom_palette = BlueAndRed())
# transfer the anti_tumoral_scoresscore to Seurat object
MDSC_scores <- p_density_MDSC$data$feature
names(MDSC_scores) <- rownames(p_density_MDSC$data)
Myeloid_Annotated <- AddMetaData(Myeloid_Annotated, metadata = MDSC_scores, col.name = "MDSC_density_score")
# Verify the addition
head(Myeloid_Annotated@meta.data)

# figure 4b #
FeaturePlot_scCustom(Myeloid_Annotated, features = "MDSC_density_score", na_cutoff = 1.0e-09)


# to determine the overall macrophage reactivity score #
library(ggpubr)

# use comprehensive gene panel
M1TAM_ScoreFeatures <- c('Csf2', 'Isg15', 'Il1a', 'Il1b', 'Il6', 'Il8', 'Il22', 'Il23a', 'Cxcl3', 'Cxcl9', 'Cxcl10', 'Ccl4', 
                         "Cd80", "Cd86", "Il1b", "Il2", "Il18")
M1TAM_ScoreFeatures <- list(M1TAM_ScoreFeatures)

Myeloid_Annotated<- AddModuleScore(Myeloid_Annotated,
                                   features = M1TAM_ScoreFeatures,
                                   ctrl = 100,
                                   name = "M1_TAM_Score")
head(Myeloid_Annotated@meta.data)

# figure 3c #
ggviolin(Myeloid_Annotated@meta.data, 
         x="Cell_of_Origin", y="M1_TAM_Score1", 
         width = 0.8,color = "black",
         fill="Cell_of_Origin", 
         xlab = F,  
         add = 'mean_sd', 
         bxp.errorbar=T,        
         bxp.errorbar.width=0.05,           
         size=0.5, 
         legend = "right") +
  scale_fill_manual(values = c("Neg_Controls" = "#0c0d0c", 
                               "Vehicle_Controls" = "#5f5f5f", 
                               "Spontaneous_AT2-derived LUAD" = "#7262ac", 
                               "AT2-derived LUAD" = "#00a0e3", 
                               "AT1-derived LUAD" = "#e31e24")) + RotatedAxis()


# to determine the overall macrophage immunosuppression score #
# try some general & well-known immunosuppressive markers #
# Figure 4c #
FeaturePlot(Myeloid_Annotated, features = c("Slc7a2", "Slc7a11", "Mmp12", "Arg1", "Msr1", "Inhba", "H2-M2", "Chil4", "Mafb"), cols = c("lightgrey", "blue"))

# use comprehensive gene panel
M2TAM_ScoreFeatures <- c("Cd163", "Spp1", "Apoe", "Gpnmb", "Trem2", "Il10", "Il13", "Il33", 'Csf1', 
                         'Ccl2', 'Ccl5', 'Ccl17', 'Ccl22', 'Arg1', 'Fga', 'Fgb', 'Mmp2', 'Mmp12', 'Vcan', 'Csf1', 'Csf3', 
                         'Cd47', 'Enpp1', 'Cgas', 'Sting1')
M2TAM_ScoreFeatures <- list(M2TAM_ScoreFeatures)

Myeloid_Annotated<- AddModuleScore(Myeloid_Annotated, 
                                   features = M2TAM_ScoreFeatures,
                                   ctrl = 100,
                                   name = "M2_TAM_Score")

# figure 4d #
ggviolin(Myeloid_Annotated@meta.data, 
         x="Cell_of_Origin", y="M2_TAM_Score1", 
         width = 0.8,color = "black",
         fill="Cell_of_Origin",
         xlab = F,  
         add = 'mean_sd', 
         bxp.errorbar=T,        
         bxp.errorbar.width=0.05,           
         size=0.5, 
         legend = "right") +
  scale_fill_manual(values = c("Neg_Controls" = "#0c0d0c", 
                               "Vehicle_Controls" = "#5f5f5f", 
                               "Spontaneous_AT2-derived LUAD" = "#7262ac", 
                               "AT2-derived LUAD" = "#00a0e3", 
                               "AT1-derived LUAD" = "#e31e24")) + RotatedAxis()


### To Characterize the differentially expressed genes (DEGs) within macrophages between Gramd2+ KRASG12D LUAD and Sftpc+ KRASG12D LUAD via Volcano plot ###
# obtain DEGs
DEGs_MF_AT1vsAT2.markers <- FindMarkers(Myeloid_Annotated,
                                        ident.1 ="AT1-derived LUAD_Macrophages", ident.2 ="AT2-derived LUAD_Macrophages", 
                                        group.by = "NEW_celltype", min.pct = 0.01, verbose = FALSE, min.cells.feature = 0, min.cells.group =0)
DEGs_MF_AT1vsAT2.markers$gene <- rownames(DEGs_MF_AT1vsAT2.markers)
DEGs_MF_AT1vsAT2.markers$fdr_corrected_pvalue <- p.adjust(DEGs_MF_AT1vsAT2.markers$p_val, method = "BH")
write.csv(DEGs_MF_AT1vsAT2.markers, file="your/file/path/DEGs_MF_AT1vsAT2_0.01.markers_annotated_ALL.CSV")
print(DEGs_MF_AT1vsAT2.markers)
head(DEGs_MF_AT1vsAT2.markers)
FDRcorrected_DEGs_MF_AT1vsAT2.markers <- p.adjust(DEGs_MF_AT1vsAT2.markers, method = "BH")
write.csv(FDRcorrected_DEGs_MF_AT1vsAT2.markers, file = "your/file/path/Supplemental Table4_fdr_corrected MP DEGs_AT1v.s.AT2.CSV")


# Load necessary libraries
library(ggplot2)
library(dplyr)
library(ggrepel)

# Load the data
data <- read.csv("your/file/path/Supplemental Table4_fdr_corrected MP DEGs_AT1v.s.AT2.CSV")

# Add a column for -log10(p-value)
data <- data %>% mutate(`-log10(p_val)` = -log10(fdr_corrected_pvalue))

# Define significance thresholds
significance_threshold <- 0.05
log2fc_threshold <- 1

data <- data %>% mutate(Significant = ifelse(avg_log2FC > log2fc_threshold & fdr_corrected_pvalue < significance_threshold, "Upregulated",
                                             ifelse(avg_log2FC < -log2fc_threshold & fdr_corrected_pvalue < significance_threshold, "Downregulated", "Not Significant")))


# Create the volcano plot without gene labels
# Figure 3e #
ggplot(data, aes(x = avg_log2FC, y = `-log10(p_val)`)) +
  geom_point(aes(color = Significant)) +
  scale_color_manual(values = c("Upregulated" = "#e31e24", "Downregulated" = "#00a0e3", "Not Significant" = "grey")) +
  geom_vline(xintercept = c(-log2fc_threshold, log2fc_threshold), linetype = "dashed") +
  geom_hline(yintercept = -log10(significance_threshold), linetype = "dashed") +
  labs(title = "Volcano Plot", x = "Average Log2 Fold Change", y = "-log10(FDR Corrected p-value)", color = "Legend") +
  theme_minimal() +
  theme(
    panel.background = element_rect(fill = "transparent", color = NA), # bg of the panel
    plot.background = element_rect(fill = "transparent", color = NA),  # bg of the plot
    legend.background = element_rect(fill = "transparent", color = NA), # get rid of legend bg
    legend.box.background = element_rect(fill = "transparent", color = NA), # get rid of legend panel bg
    panel.grid.major = element_blank(), # remove major grid lines
    panel.grid.minor = element_blank(), # remove minor grid lines
    axis.line = element_line(color = "black"), # set axis lines to black
    axis.ticks = element_line(color = "black") # set axis ticks to black
  )

# Create the volcano plot with gene labels
if (!requireNamespace("ggrepel", quietly = TRUE)) {
  install.packages("ggrepel")
}

library(ggrepel)

ggplot(data, aes(x = avg_log2FC, y = `-log10(p_val)`)) +
  geom_point(aes(color = Significant)) +
  scale_color_manual(values = c("Upregulated" = "#e31e24", "Downregulated" = "#00a0e3", "Not Significant" = "grey")) +
  geom_vline(xintercept = c(-log2fc_threshold, log2fc_threshold), linetype = "dashed") +
  geom_hline(yintercept = -log10(significance_threshold), linetype = "dashed") +
  geom_text_repel(data = subset(data, Significant != "Not Significant"), aes(label = gene), size = 3, max.overlaps = 10) +
  labs(title = "Volcano Plot", x = "Average Log2 Fold Change", y = "-log10(FDR Corrected p-value)", color = "Legend") +
  theme_minimal() +
  theme(
    panel.background = element_rect(fill = "transparent", color = NA), # bg of the panel
    plot.background = element_rect(fill = "transparent", color = NA),  # bg of the plot
    legend.background = element_rect(fill = "transparent", color = NA), # get rid of legend bg
    legend.box.background = element_rect(fill = "transparent", color = NA), # get rid of legend panel bg
    panel.grid.major = element_blank(), # remove major grid lines
    panel.grid.minor = element_blank(), # remove minor grid lines
    axis.line = element_line(color = "black"), # set axis lines to black
    axis.ticks = element_line(color = "black") # set axis ticks to black
  )


# To interrogate the enrichment of biological function/pathways via GSEA based on PANTHER database # 
# Load necessary libraries
library(ggplot2)
library(readxl)
library(dplyr)

# Read the data from the CSV file
data <- read_excel('your/file/path/Supplemental Table5_PANTHER pathways for MP DEGs_AT1v.s.AT2.xlsx')

# Arrange data by Log2_Fold_Enrichment
data <- data %>%
  arrange(desc(Log2_Fold_Enrichment))

# Create the bubble plot
bubble_plot <- ggplot(data, aes(x = Log2_Fold_Enrichment, y = reorder(PANTHER_Pathways, Log2_Fold_Enrichment), size = gene_count, color = p_adj)) +
  geom_point(alpha = 1) +
  scale_color_gradient(low = "#7216b2", high = "#fcff00", name = "p_adj") +
  labs(title = "Bubble Plot of PANTHER Pathways",
       x = "Log2 Fold Enrichment",
       y = "PANTHER Pathways",
       size = "Gene count") +
  geom_vline(xintercept = c(0.5, -0.5), linetype = "dashed", color = "black") +
  theme_minimal(base_size = 15) +
  theme(
    plot.title = element_text(hjust = 0.5, face = "bold", size = 20),
    axis.title.x = element_text(face = "bold"),
    axis.title.y = element_text(face = "bold"),
    axis.text.y = element_text(size = 12),
    legend.position = "right",
    legend.title = element_text(face = "bold"),
    legend.text = element_text(size = 12),
    panel.background = element_rect(fill = "#E6E6E6", color = NA),
    plot.background = element_rect(fill = "transparent", color = NA),
    panel.grid.major = element_line(size = 0.5, linetype = 'solid', color = 'white'),
    axis.line = element_line(color = "black")
  ) +
  guides(size = guide_legend(override.aes = list(alpha = 1)))


# Display the plot
# Figure 3f #
print(bubble_plot)


########## generate supplemental figure 4 ########## 
########## Set up GSEA function ############
GSEA = function(gene_list, GO_file, pval) {
  set.seed(54321)
  library(dplyr)
  library(fgsea)
  
  if ( any( duplicated(names(gene_list)) )  ) {
    warning("Duplicates in gene names")
    gene_list = gene_list[!duplicated(names(gene_list))]
  }
  if  ( !all( order(gene_list, decreasing = TRUE) == 1:length(gene_list)) ){
    warning("Gene list not sorted")
    gene_list = sort(gene_list, decreasing = TRUE)
  }
  myGO = fgsea::gmtPathways(GO_file)
  
  fgRes <- fgsea::fgsea(pathways = myGO,
                        stats = gene_list,
                        minSize=15, ## minimum gene set size
                        maxSize=400 ## maximum gene set sizeDEGs_MF_AT1vsAT2_0.01.markers_annotated_ALL.CSV
  ) %>% 
    as.data.frame() %>% 
    dplyr::filter(padj < !!pval) %>% 
    arrange(desc(NES))
  message(paste("Number of signficant gene sets =", nrow(fgRes)))
  
  message("Collapsing Pathways -----")
  concise_pathways = collapsePathways(data.table::as.data.table(fgRes),
                                      pathways = myGO,
                                      stats = gene_list)
  fgRes = fgRes[fgRes$pathway %in% concise_pathways$mainPathways, ]
  message(paste("Number of gene sets after collapsing =", nrow(fgRes)))
  
  fgRes$Enrichment = ifelse(fgRes$NES > 0, "Up-regulated", "Down-regulated")
  filtRes = rbind(head(fgRes, n = 10),
                  tail(fgRes, n = 10 ))
  
  total_up = sum(fgRes$Enrichment == "Up-regulated")
  total_down = sum(fgRes$Enrichment == "Down-regulated")
  header = paste0("Top 10 (Total pathways: Up=", total_up,", Down=",    total_down, ")")
  
  colos = setNames(c("firebrick2", "dodgerblue2"),
                   c("Up-regulated", "Down-regulated"))
  
  g1= ggplot(filtRes, aes(reorder(pathway, NES), NES)) +
    geom_point( aes(fill = Enrichment, size = size), shape=21) +
    scale_fill_manual(values = colos ) +
    scale_size_continuous(range = c(2,10)) +
    geom_hline(yintercept = 0) +
    coord_flip() +
    labs(x="Pathway", y="Normalized Enrichment Score",
         title=header) 
  
  output = list("Results" = fgRes, "Plot" = g1)
  return(output)
}


############ Set up plotting function ###############
plot_geneset_clusters = function( gs_results, GO_file, min.sz = 4, main="GSEA clusters"){
  library(ggplot2)
  library(ggrepel)
  library(stringr)
  
  myGO = fgsea::gmtPathways(GO_file)
  df = matrix(nrow=nrow(gs_results), ncol = nrow(gs_results), data = 0)
  rownames(df) = colnames(df) = gs_results$pathway
  
  for ( i in 1:nrow(gs_results)) {
    genesI =  unlist(myGO[names(myGO) == gs_results$pathway[i] ])
    for (j in 1:nrow(gs_results)) {
      genesJ = unlist(myGO[names(myGO) == gs_results$pathway[j] ])
      ## Jaccards distance  1 - (intersection / union )
      overlap = sum(!is.na(match(genesI, genesJ )))
      jaccards = overlap / length(unique(c(genesI, genesJ) ))
      df[i,j] = 1-jaccards
    }
  }
  
  ## Cluster nodes using dynamic tree cut, for colors
  distMat = as.dist(df)
  dendro = hclust(distMat, method = "average" )
  clust = dynamicTreeCut::cutreeDynamicTree( dendro, minModuleSize = min.sz )
  ## Note: in dynamicTreeCut, cluster 0, is a garbage cluster for things that dont cluster, so we remove it
  
  gs_results$Cluster = clust
  gs_results = gs_results[gs_results$Cluster != 0, ]
  
  ## select gene sets to label for each clusters
  bests = gs_results %>%  
    group_by( Cluster ) %>% 
    top_n(wt = abs(size), n = 1) %>% 
    .$pathway
  ## determine cluster order for plotting
  clust_ords = gs_results %>% 
    group_by( Cluster ) %>% 
    reframe("Average" = NES ) %>% 
    arrange(desc(Average)) %>% 
    .$Cluster %>% 
    unique
  
  gs_results$Cluster = factor(gs_results$Cluster, levels = clust_ords)
  
  gs_results$Label = ""
  gs_results$Label[gs_results$pathway %in% bests ] = gs_results$pathway[gs_results$pathway %in% bests ]
  gs_results$Label = str_remove(gs_results$Label, "GO_")
  gs_results$Label = tolower(gs_results$Label)
  
  g1 = ggplot(gs_results, aes(x = Cluster, y = NES, label = Label )) +
    geom_jitter( aes(color = Cluster,  size = size), alpha = 0.8, height = 0, width = 0.2 ) +
    scale_size_continuous(range = c(0.5,5)) +
    geom_text_repel( force = 2, max.overlaps = Inf) +
    ggtitle(main) 
  
  return(g1)
}


########### Run GSEA #############
library(dplyr)
GO_file = "your/file/path/AT1LUAD_scRNA/m5.all.v2023.1.Mm.symbols.gmt"

S4table = read.csv("your/file/path/Macrophages/DEGs_MF_AT1vsAT2_0.01.markers_Annotated.CSV")


S4table <- S4table %>% filter(gene != "")
gene_list = S4table$avg_log2FC
names(gene_list) = S4table$gene
gene_list = sort(gene_list, decreasing = TRUE)
gene_list = gene_list[!duplicated(names(gene_list))]
head(gene_list)

res = GSEA(gene_list, GO_file, pval = 0.05)
dim(res$Results)
# supplemental figure 4 #
res$Plot


############################################################################################################################################################
####################################### inferCNV analysis - a prerequisite step for cell-cell communication analysis ####################################### 
############################################################################################################################################################

AT1LUAD_BatchIntegrated <- readRDS("your/file/path/LUAD_BatchIntegrated.RDS")
######################### Extract AT1 cells in AT1-derived LUAD group #########################
AT1_Cells <- subset(x = AT1LUAD_BatchIntegrated, subset = (predicted.celltype_level3 == "AT1"))

######################### Extract AT2 cells in AT1-derived LUAD group #########################
AT2_Cells <- subset(x = AT1LUAD_BatchIntegrated, subset = (predicted.celltype_level3 == "AT2"))

######################### Extract AT1 cells in AT1-derived LUAD group #########################
AT1AT2_Cells <- subset(x = AT1LUAD_BatchIntegrated, subset = (predicted.celltype_level3 == "AT1/AT2"))

################################# Do integration analysis for all AECs ################################# 
######################### Merge AT1 +  AT2 + AT1/AT2 to generate a merged Alveolar Epithelial Cell (AEC) object######################### 
AECs <- merge(AT1_Cells, y = c(AT2_Cells, AT1AT2_Cells), project = "AT1LUAD")
View(AECs)
range(AECs$nFeature_RNA)
range(AECs$nCount_RNA)
range(AECs$percent.mt)

##1.2. selecting cells for further analysis
# visualize QC metrics as a violin plot
VlnPlot(AECs, features = c("nFeature_RNA", "nCount_RNA","percent.mt"), group.by = c("Mouse_ID"), ncol = 3,
        log = TRUE, pt.size = 0) + NoLegend()

##2. Data normalization
AECs@assays[["RNA"]]@data@x

AECs <- NormalizeData(AECs, normalization.method = "LogNormalize", scale.factor = 10000)

AECs@assays[["RNA"]]@data@x

##3. Identification of highly variable features
AECs

AECs <- FindVariableFeatures(AECs, selection.method = "vst", nfeatures = 2000) #vst = variance stablizing transformatin

#Top features and featurePlot
AECs@assays[["RNA"]]@var.features #first 1000 variable features
top10VariableFeatures <- head(VariableFeatures(AECs), 10)
plot1 <- VariableFeaturePlot(AECs)
plot2 <- LabelPoints(plot = plot1, points = top10VariableFeatures, repel = TRUE)
plot1 + plot2


##4. Data scaling
AECs <- ScaleData(object = AECs) #2000 identified variable features
AECs@assays[["RNA"]]@scale.data
View(AECs@commands)


##5. Perform PCA, linear dimensional reduction, on the scaled data (linear dimensional reduction)
AECs <- RunPCA(AECs, features = VariableFeatures(object = AECs))

#Examine and visualize PCA results a few different ways: DimPlot(); VizDimReduction(); DimHeatmap()
DimPlot(AECs, reduction = "pca", dims = c(1,2))
DimPlot(AECs, reduction = "pca", dims = c(1,10))
DimPlot(AECs, reduction = "pca", dims = c(1,50))

#Determine the "dimensional" of the dataset
AECs <- JackStraw(AECs, num.replicate = 100, dims = 20)
AECs <- ScoreJackStraw(AECs, dims = 1:20)
JackStrawPlot(AECs, dims = 1:20)

#more direct way: ElbowPlot()
ElbowPlot(AECs) #default = 20 PC
ElbowPlot(AECs, ndims = 50, reduction = "pca") #more PCs, more precise 

DimHeatmap(AECs, dims = 1:15, cells = 500, balanced = TRUE)

### run Harmony -----------------
AECs <- AECs %>%
  RunHarmony(group.by.vars = 'Batch', plot_covergence = FALSE)

AECs@reductions

AECs.embed <- Embeddings(AECs, "harmony")
AECs.embed[1:10,1:10]

# remove the "Spontaneous_AT2-derived LUAD" group to solely retrieve malignant AECs from Sftpc+ KRASG12D LUAD and Gramd2+ KRASG12D LUAD
AECs <- subset(x = AECs, subset = (Cell_of_Origin == c("Spontaneous_AT2-derived LUAD")), invert = TRUE)
# to remove the unspecific cells
AECs <- subset(x = AECs, subset = (predicted.lineage_level1 == c("Immune",
                                                                 "Endothelial",
                                                                 "Mesenchymal")), invert = TRUE)
# Do UMAP and clustering using ** Harmony embeddings instead of PCA **
AECs <- AECs %>%
  RunUMAP(reduction = 'harmony', dims = 1:20) %>%
  FindNeighbors(reduction = 'harmony', dims = 1:20) %>%
  FindClusters(resolution = 0.3) #17 clusters

# to verify the identities of cells
AECs$seurat_clusters <- as.factor(as.numeric(as.character(AECs$seurat_clusters)) +1)
AECs$Cell_of_Origin <- factor(AECs$Cell_of_Origin, levels = c("Neg_Controls", "Vehicle_Controls", "AT2-derived LUAD", "AT1-derived LUAD"))

DimPlot(AECs, reduction = "umap", label = T, group.by = "Batch", repel = TRUE)
DimPlot(AECs, reduction = "umap", label = T, group.by = "seurat_clusters", repel = TRUE)
DimPlot(AECs, reduction = "umap", label = FALSE, group.by = "Disease_Status", cols = c("#535164", "#F2542C"))
DimPlot(AECs, reduction = "umap", label = FALSE, group.by = "Cell_of_Origin", cols = c("#0c0d0c", "#535164", "#F2542C", "#000099"))
DimPlot(AECs, reduction = "umap", label = FALSE, group.by = "predicted.lineage_level1")
DimPlot(AECs, reduction = "umap", label = FALSE, group.by = "predicted.lineage_level2")
DimPlot(AECs, reduction = "umap", label = FALSE, group.by = "predicted.celltype_level3")

p1 <- DimPlot(AECs, reduction = "umap", label = T, group.by = "seurat_clusters", repel = TRUE)
P1 <- DimPlot(AECs, reduction = "umap", label = FALSE, group.by = "predicted.celltype_level3")
p1 + P1

saveRDS(AECs, file = "your/file/path/AECs.RDS")


########################### to perform inferCNV ########################### 
Barcodes <- AECs@active.ident
Barcodes <- as.data.frame(Barcodes)
Barcodes$RowName <- rownames(Barcodes)
names(Barcodes)[1] <- "cluster"
names(Barcodes)[2] <- "Barcode"
AECs <- AddMetaData(AECs, Barcodes$Barcode, col.name = "Barcode")


# to retrieve exprMatrix from AECs Seurat object
# make raw count matrix
exprMatrix <- as.matrix(GetAssayData(AECs, slot='counts'))
write.table(exprMatrix, 'your/file/path/exprMatrix_AECs_Full.txt', col.names=NA, sep='\t')

# 准备 annotation file
AECs$LUADCluster <- paste(AECs$Disease_Status, sep = "_", AECs$seurat_clusters)


AEC_af <- FetchData(AECs, c("predicted.celltype_level2", "Disease_Status", "seurat_clusters", "Barcode")) %>%
  # filter(cell_id %in% cell_subset) %>%
  mutate(cell_type_infercnv = ifelse(Disease_Status != "Healthy", paste0("LUAD_", seurat_clusters), paste0("Normal_", seurat_clusters)),
         cell_id = str_replace_all(Barcode, ":", "_")) %>% 
  select(cell_id, cell_type_infercnv) %>% 
  as_tibble

baseline_cell_types <- sort(unique(AEC_af$cell_type_infercnv)[!str_detect(unique(AEC_af$cell_type_infercnv), "malignant")])

write.table(AEC_af, file = 'your/file/path/AEC_af_Full.txt', sep = "\t", col.names = F, row.names = F)


# prepare for the gene_order_file in python or terminal
#gene_position.txt can be downloaded from this link：wget https://data.broadinstitute.org/Trinity/CTAT/cnv/gencode_v21_gen_pos.complete.txt

# or DIY your own .txt file: https://github.com/broadinstitute/inferCNV/tree/master/scripts 
# download GTF file from: https://www.gencodegenes.org/mouse/  

# DIY processing:
#python gtf_to_position_file.py --attribute_name "gene_name"  gencode.vM34.basic.annotation.gtf gene_pos.txt

#filter out autosomes 1-19 from gene_position.txt:
#cat gene_pos.txt | grep -E 'chr[1-9]\>|chr1[0-9]\>' > filtered_gene_position.txt

#last check the identities of chromosomes in "filtered_gene_position.txt" file
#cat filtered_gene_position.txt | awk '{print $2}' |uniq 

# To find normal AECs from control samples as anchors/reference cells: determined Normal_7 & Normal_7 are normal healthy AECs from control samples
DimPlot(AECs, reduction = 'umap', group.by = "LUADCluster", label = TRUE)

# Create infercnv object for AECs
infercnv_obj_AECs = CreateInfercnvObject(delim = '\t',
                                         raw_counts_matrix = 'your/file/path/exprMatrix_AECs_Full.txt',
                                         annotations_file = 'your/file/path/AEC_af_Full.txt',
                                         gene_order_file = 'your/file/path/filtered_gene_position.txt',
                                         ref_group_names = c("Normal_7", "Normal_8"))


# run inferCNV
# perform infercnv operations to reveal cnv signal
dir.create("your/file/path/AECs")
infercnv_obj_AECs = infercnv::run(infercnv_obj_AECs,
                                  cutoff=0.1, 
                                  out_dir='your/file/path/AECs/', 
                                  cluster_by_groups=TRUE, 
                                  denoise=TRUE,
                                  HMM=TRUE,
                                  write_expr_matrix = TRUE)

# save inferCNV object for future usage
saveRDS(infercnv_obj_AECs, file="LUAD Malignancy_infercnv_obj_AECs_Full.rds", compress="xz")

# Data Visulization
plot_cnv(infercnv_obj_AECs,
         out_dir='inferCNV/AECs/',
         obs_title = "Malignant Cells",
         ref_title = "Normal Cells",
         cluster_by_groups = TRUE,
         plot_chr_scale = TRUE,
         color_safe_pal = TRUE,
         png_res=100,
         output_filename = "InferCNV_scaled_to_chr")


# VlnPlot to inspect CNV score
data_AECs=read.table("inferCNV/AECs/infercnv.observations.txt", header=T)
expr_AECs=data_AECs %>% as.matrix()
expr.scale =scale(t(expr_AECs))
tmp1=sweep(expr.scale, 2, apply(expr.scale, 2, min),'-')
tmp2=apply(expr.scale, 2, max) - apply(expr.scale,2,min)
expr_1=t(2*sweep(tmp1, 2, tmp2, "/")-1)
cnv_score_AECs=as.data.frame(colSums(expr_1 * expr_1))
colnames(cnv_score_AECs)="cnv_score"
cnv_score_AECs=rownames_to_column(cnv_score_AECs, var='cell')

# Ensure `cnv_score` is a data frame and not an atomic vector
cnv_score_AECs <- as.data.frame(cnv_score_AECs)
colnames(cnv_score_AECs)[1] <- "cell"
cnv_score_AECs$cell <- gsub("\\.1$", "-1", cnv_score_AECs$cell)

# Verify the changes
head(cnv_score_AECs$cell)
head(rownames(AECs@meta.data))

# Extract cell names from Seurat object
cells_AECs <- rownames(AECs@meta.data)
head(cells_AECs)

# Correctly filter to include only cells present in AECs
cnv_score_AECs <- cnv_score_AECs %>% filter(cell %in% cells_AECs)

# Reorder `cnv_score` to match the order of cells in `AECs`
cnv_score_AECs <- cnv_score_AECs[match(cells_AECs, cnv_score_AECs$cell), ]


AECs@meta.data$CNV_score <- cnv_score_AECs$cnv_score
FeaturePlot(AECs, features = "CNV_score", cols = c("lightgrey", "blue")) +
  labs(title = "UMAP Plot of CNV Score") +
  theme_minimal()


# to generate a new dataframe incoporating cell barcode indentities and cnv scores
test_AECs=merge(cnv_score_AECs, AEC_af, by = 'cell')
write.table(test_AECs, file = 'inferCNV/Barcode_Cluster_CNVscore.txt', sep = "\t", col.names = F, row.names = F)


# split the AECs of LUAD samples apart from AECs of Normal healthy samples
LUAD_data_AECs <- subset(test_AECs, grepl("LUAD", cell_type_infercnv))
Normal_data_AECs <- subset(test_AECs, grepl("Normal", cell_type_infercnv))

### to inspect the distribution of cnv score between LUAD samples and controls
# istribution of cnv score in LUAD samples
hist(LUAD_data_AECs$cnv_score, breaks = 50, col = "skyblue",
     main = "Histogram of Bimodal Distribution",
     xlab = "Value",
     ylab = "Frequency")
hist(log10(LUAD_data_AECs$cnv_score), breaks = 50, col = "lightgreen",
     main = "Histogram of Bimodal Distribution (Log Scale)",
     xlab = "Log10(Value)",
     ylab = "Frequency")

# istribution of cnv score in controls 
hist(Normal_data_AECs$cnv_score, breaks = 50, col = "skyblue",
     main = "Histogram of Bimodal Distribution",
     xlab = "Value",
     ylab = "Frequency")
hist(log10(Normal_data_AECs$cnv_score), breaks = 50, col = "lightgreen",
     main = "Histogram of Bimodal Distribution (Log Scale)",
     xlab = "Log10(Value)",
     ylab = "Frequency")

# determine cutoff of cnv score based on 95 percentile of cnv score in controls 
cutoff_AECs <- quantile(Normal_data_AECs$cnv_score, 0.95)
print(cutoff_AECs)
cnv_threshold_AECs <- cutoff_AECs

# Create the FeaturePlot
P <- FeaturePlot(AECs, reduction = "umap", features = "CNV_score") + 
  scale_color_gradientn(colors = c("grey", "#2017a5", "#fb6180", "#fb9c4a", "#eef54c"))
print(P)
# Extract the data used in the FeaturePlot
plot_data <- P$data

# Identify cells above the threshold
highlight_data <- plot_data[plot_data$CNV_score > cnv_threshold_AECs, ]

# Add the highlighting layer
P + geom_point(data = highlight_data, aes(x = umap_1, y = umap_2), color = "firebrick", size = 0.5) +
  ggtitle("UMAP Plot of CNV Score with Highlighted Clusters") +
  theme_minimal()
# By far, we determined LUAD_2, 3, 4, 5, 6 manifest significant high CNV score and CNV pattern

######################################################################################################################################################################## 
############################################################# Construct Cell-cell interaction (CCI) objects############################################################# 
######################################################################################################################################################################## 

############################################################# Construct AECs objects for each LUAD models ############################################################# 
AECs <- readRDS("your/file/path/AECs.RDS")

Myeloid_Annotated <- readRDS("/Users/m.y./Documents/R/R_working directory/AT1LUAD_scRNA/Myeloid_Annotated.RDS")

AECs$CoO_Cluster <- paste(AECs$Cell_of_Origin, sep = "_", AECs$seurat_clusters)
DimPlot(AECs, reduction = "umap", label = T, group.by = "CoO_Cluster")

MalignantAECs_in_AT1LUAD <- subset(x = AECs, subset = (CoO_Cluster == c("AT1-derived LUAD_2",
                                                                        "AT1-derived LUAD_3",
                                                                        "AT1-derived LUAD_4",
                                                                        "AT1-derived LUAD_5",
                                                                        "AT1-derived LUAD_6")))
MalignantAECs_in_AT2LUAD <- subset(x = AECs, subset = (CoO_Cluster == c("AT2-derived LUAD_2",
                                                                        "AT2-derived LUAD_3",
                                                                        "AT2-derived LUAD_4",
                                                                        "AT2-derived LUAD_5",
                                                                        "AT2-derived LUAD_6")))
# inspect quality of cells
range(MalignantAECs_in_AT1LUAD$nFeature_RNA)
range(MalignantAECs_in_AT1LUAD$nCount_RNA)
range(MalignantAECs_in_AT2LUAD$nFeature_RNA)
range(MalignantAECs_in_AT2LUAD$nCount_RNA)
range(MalignantAECs_in_AT1LUAD$percent.mt)
range(MalignantAECs_in_AT2LUAD$percent.mt)

##1.2. selecting cells for further analysis
# visualize QC metrics as a violin plot
VlnPlot(MalignantAECs_in_AT1LUAD, features = c("nFeature_RNA", "nCount_RNA","percent.mt"), group.by = c("Mouse_ID"), ncol = 3,
        log = TRUE, pt.size = 0) + NoLegend()

##2. Data normalization
MalignantAECs_in_AT1LUAD <- NormalizeData(MalignantAECs_in_AT1LUAD, normalization.method = "LogNormalize", scale.factor = 10000)
MalignantAECs_in_AT2LUAD <- NormalizeData(MalignantAECs_in_AT2LUAD, normalization.method = "LogNormalize", scale.factor = 10000)

##3. Identification of highly variable features
MalignantAECs_in_AT1LUAD <- FindVariableFeatures(MalignantAECs_in_AT1LUAD, selection.method = "vst", nfeatures = 2000) #vst = variance stablizing transformatin
MalignantAECs_in_AT2LUAD <- FindVariableFeatures(MalignantAECs_in_AT2LUAD, selection.method = "vst", nfeatures = 2000) #vst = variance stablizing transformatin

##4. Data scaling
MalignantAECs_in_AT1LUAD <- ScaleData(object = MalignantAECs_in_AT1LUAD) 
MalignantAECs_in_AT2LUAD <- ScaleData(object = MalignantAECs_in_AT2LUAD)

MalignantAECs_in_AT1LUAD@assays[["RNA"]]@scale.data
View(MalignantAECs_in_AT1LUAD@commands)


###Perform PCA, linear dimensional reduction, on the scaled data (linear dimensional reduction)
MalignantAECs_in_AT1LUAD <- RunPCA(MalignantAECs_in_AT1LUAD, features = VariableFeatures(object = MalignantAECs_in_AT1LUAD))
MalignantAECs_in_AT2LUAD <- RunPCA(MalignantAECs_in_AT2LUAD, features = VariableFeatures(object = MalignantAECs_in_AT2LUAD))
# generate 5 PCs, first 30 features in each PC

#more direct way: ElbowPlot()
ElbowPlot(MalignantAECs_in_AT1LUAD) #default = 20 PC
ElbowPlot(MalignantAECs_in_AT1LUAD, ndims = 50, reduction = "pca") #dims = 15
ElbowPlot(MalignantAECs_in_AT2LUAD) #default = 20 PC
ElbowPlot(MalignantAECs_in_AT2LUAD, ndims = 50, reduction = "pca") #dims = 15


### run Harmoney -----------------
MalignantAECs_in_AT1LUAD <- MalignantAECs_in_AT1LUAD %>%
  RunHarmony(group.by.vars = 'Batch', plot_covergence = FALSE)

MalignantAECs_in_AT1LUAD@reductions

MalignantAECs_in_AT1LUAD.embed <- Embeddings(MalignantAECs_in_AT1LUAD, "harmony")
MalignantAECs_in_AT1LUAD.embed[1:10,1:10]

MalignantAECs_in_AT2LUAD <- MalignantAECs_in_AT2LUAD %>%
  RunHarmony(group.by.vars = 'Batch', plot_covergence = FALSE)

MalignantAECs_in_AT2LUAD@reductions

MalignantAECs_in_AT2LUAD.embed <- Embeddings(MalignantAECs_in_AT2LUAD, "harmony")
MalignantAECs_in_AT2LUAD.embed[1:10,1:10]
# Do UMAP and clustering using ** Harmony embeddings instead of PCA **
MalignantAECs_in_AT1LUAD <- MalignantAECs_in_AT1LUAD %>%
  RunUMAP(reduction = 'harmony', dims = 1:15) %>%
  FindNeighbors(reduction = 'harmony', dims = 1:15) %>%
  FindClusters(resolution = 0.5) #6 clusters

MalignantAECs_in_AT2LUAD <- MalignantAECs_in_AT2LUAD %>%
  RunUMAP(reduction = 'harmony', dims = 1:15) %>%
  FindNeighbors(reduction = 'harmony', dims = 1:15) %>%
  FindClusters(resolution = 0.5) #9 clusters


MalignantAECs_in_AT1LUAD$seurat_clusters <- as.factor(as.numeric(as.character(MalignantAECs_in_AT1LUAD$seurat_clusters)) +1)
MalignantAECs_in_AT2LUAD$seurat_clusters <- as.factor(as.numeric(as.character(MalignantAECs_in_AT2LUAD$seurat_clusters)) +1)

DimPlot(MalignantAECs_in_AT1LUAD, reduction = "umap", label = T, group.by = "Batch", repel = TRUE)
DimPlot(MalignantAECs_in_AT1LUAD, reduction = "umap", label = T, group.by = "seurat_clusters", repel = TRUE)
DimPlot(MalignantAECs_in_AT1LUAD, reduction = "umap", label = FALSE, group.by = "Mouse_ID")

DimPlot(MalignantAECs_in_AT2LUAD, reduction = "umap", label = T, group.by = "Batch", repel = TRUE)
DimPlot(MalignantAECs_in_AT2LUAD, reduction = "umap", label = T, group.by = "seurat_clusters", repel = TRUE)
DimPlot(MalignantAECs_in_AT2LUAD, reduction = "umap", label = FALSE, group.by = "Mouse_ID")

# If you want to replace 'active.ident' with 'cell_types' completely
MalignantAECs_in_AT1LUAD@meta.data$AECcelltypes <- MalignantAECs_in_AT1LUAD$predicted.celltype_level3
DimPlot(MalignantAECs_in_AT1LUAD, group.by = 'AECcelltypes', reduction = "umap", label = T, repel = TRUE)
MalignantAECs_in_AT2LUAD@meta.data$AECcelltypes <- MalignantAECs_in_AT2LUAD$predicted.celltype_level3
DimPlot(MalignantAECs_in_AT2LUAD, group.by = 'AECcelltypes', reduction = "umap", label = T, repel = TRUE)
# double check if the "active.ident" has been replaced successfully
DimPlot(MalignantAECs_in_AT1LUAD, reduction = 'umap')
DimPlot(MalignantAECs_in_AT2LUAD, reduction = 'umap')

# save for future usage
saveRDS(MalignantAECs_in_AT1LUAD, file = "/Users/m.y./Documents/R/R_working directory/AT1LUAD_scRNA/MalignantAECs_in_AT1LUAD.RDS")
saveRDS(MalignantAECs_in_AT2LUAD, file = "/Users/m.y./Documents/R/R_working directory/AT1LUAD_scRNA/MalignantAECs_in_AT2LUAD.RDS")


############################################################# Construct Myeloid objects for each LUAD models ############################################################# 
Myeloid_in_AT1LUAD <- subset(x = Myeloid_Annotated, subset = (Cell_of_Origin == c("AT1-derived LUAD")))
Myeloid_in_AT2LUAD <- subset(x = Myeloid_Annotated, subset = (Cell_of_Origin == c("AT2-derived LUAD")))

range(Myeloid_in_AT1LUAD$nFeature_RNA)
range(Myeloid_in_AT1LUAD$nCount_RNA)
range(Myeloid_in_AT2LUAD$nFeature_RNA)
range(Myeloid_in_AT2LUAD$nCount_RNA)
range(Myeloid_in_AT1LUAD$percent.mt)
range(Myeloid_in_AT2LUAD$percent.mt)

##1.2. selecting cells for further analysis
# visualize QC metrics as a violin plot
VlnPlot(Myeloid_in_AT1LUAD, features = c("nFeature_RNA", "nCount_RNA","percent.mt"), group.by = c("Mouse_ID"), ncol = 3,
        log = TRUE, pt.size = 0) + NoLegend()

##2. Data normalization
Myeloid_in_AT1LUAD <- NormalizeData(Myeloid_in_AT1LUAD, normalization.method = "LogNormalize", scale.factor = 10000)
Myeloid_in_AT2LUAD <- NormalizeData(Myeloid_in_AT2LUAD, normalization.method = "LogNormalize", scale.factor = 10000)

##3. Identification of highly variable features
Myeloid_in_AT1LUAD <- FindVariableFeatures(Myeloid_in_AT1LUAD, selection.method = "vst", nfeatures = 2000) #vst = variance stablizing transformatin
Myeloid_in_AT2LUAD <- FindVariableFeatures(Myeloid_in_AT2LUAD, selection.method = "vst", nfeatures = 2000) #vst = variance stablizing transformatin

##4. Data scaling
Myeloid_in_AT1LUAD <- ScaleData(object = Myeloid_in_AT1LUAD) #2000 identified variable features
Myeloid_in_AT2LUAD <- ScaleData(object = Myeloid_in_AT2LUAD)

Myeloid_in_AT1LUAD@assays[["RNA"]]@scale.data
View(Myeloid_in_AT1LUAD@commands)


###Perform PCA, linear dimensional reduction, on the scaled data (linear dimensional reduction)
Myeloid_in_AT1LUAD <- RunPCA(Myeloid_in_AT1LUAD, features = VariableFeatures(object = Myeloid_in_AT1LUAD))
Myeloid_in_AT2LUAD <- RunPCA(Myeloid_in_AT2LUAD, features = VariableFeatures(object = Myeloid_in_AT2LUAD))
# generate 5 PCs, first 30 features in each PC

#more direct way: ElbowPlot()
ElbowPlot(Myeloid_in_AT1LUAD) #default = 20 PC
ElbowPlot(Myeloid_in_AT1LUAD, ndims = 50, reduction = "pca") #dims = 18
ElbowPlot(Myeloid_in_AT2LUAD) #default = 20 PC
ElbowPlot(Myeloid_in_AT2LUAD, ndims = 50, reduction = "pca") #dims = 14


### run Harmoney -----------------
Myeloid_in_AT1LUAD <- Myeloid_in_AT1LUAD %>%
  RunHarmony(group.by.vars = 'Batch', plot_covergence = FALSE)

Myeloid_in_AT1LUAD@reductions

Myeloid_in_AT1LUAD.embed <- Embeddings(Myeloid_in_AT1LUAD, "harmony")
Myeloid_in_AT1LUAD.embed[1:10,1:10]

Myeloid_in_AT2LUAD <- Myeloid_in_AT2LUAD %>%
  RunHarmony(group.by.vars = 'Batch', plot_covergence = FALSE)

Myeloid_in_AT2LUAD@reductions

Myeloid_in_AT2LUAD.embed <- Embeddings(Myeloid_in_AT2LUAD, "harmony")
Myeloid_in_AT2LUAD.embed[1:10,1:10]
# Do UMAP and clustering using ** Harmony embeddings instead of PCA **
Myeloid_in_AT1LUAD <- Myeloid_in_AT1LUAD %>%
  RunUMAP(reduction = 'harmony', dims = 1:18) %>%
  FindNeighbors(reduction = 'harmony', dims = 1:18) %>%
  FindClusters(resolution = 0.5) #9 clusters

Myeloid_in_AT2LUAD <- Myeloid_in_AT2LUAD %>%
  RunUMAP(reduction = 'harmony', dims = 1:14) %>%
  FindNeighbors(reduction = 'harmony', dims = 1:14) %>%
  FindClusters(resolution = 0.5) #9 clusters


Myeloid_in_AT1LUAD$seurat_clusters <- as.factor(as.numeric(as.character(Myeloid_in_AT1LUAD$seurat_clusters)) +1)
Myeloid_in_AT2LUAD$seurat_clusters <- as.factor(as.numeric(as.character(Myeloid_in_AT2LUAD$seurat_clusters)) +1)


DimPlot(Myeloid_in_AT1LUAD, reduction = "umap", label = T, group.by = "Batch", repel = TRUE)
DimPlot(Myeloid_in_AT1LUAD, reduction = "umap", label = T, group.by = "seurat_clusters", repel = TRUE)
DimPlot(Myeloid_in_AT1LUAD, reduction = "umap", label = FALSE, group.by = "Mouse_ID")
DimPlot(Myeloid_in_AT1LUAD, reduction = "umap", label = T, group.by = "Celltypes", repel = TRUE)

DimPlot(Myeloid_in_AT2LUAD, reduction = "umap", label = T, group.by = "Batch", repel = TRUE)
DimPlot(Myeloid_in_AT2LUAD, reduction = "umap", label = T, group.by = "seurat_clusters", repel = TRUE)
DimPlot(Myeloid_in_AT2LUAD, reduction = "umap", label = FALSE, group.by = "Mouse_ID")
DimPlot(Myeloid_in_AT2LUAD, reduction = "umap", label = T, group.by = "Celltypes", repel = TRUE)

saveRDS(Myeloid_in_AT1LUAD, file = "/Users/m.y./Documents/R/R_working directory/AT1LUAD_scRNA/Myeloid_in_AT1LUAD.RDS")
saveRDS(Myeloid_in_AT2LUAD, file = "/Users/m.y./Documents/R/R_working directory/AT1LUAD_scRNA/Myeloid_in_AT2LUAD.RDS")


########################### construct CCI objects by merging Myeloid cells and malignant AECs in each LUAD model respectively ###########################
CCIObj_in_AT1LUAD <- merge(MalignantAECs_in_AT1LUAD, y = c(Myeloid_in_AT1LUAD), project = "AT1LUAD")
CCIObj_in_AT2LUAD <- merge(MalignantAECs_in_AT2LUAD, y = c(Myeloid_in_AT2LUAD), project = "AT2LUAD")


range(CCIObj_in_AT1LUAD$nFeature_RNA)
range(CCIObj_in_AT1LUAD$nCount_RNA)
range(CCIObj_in_AT2LUAD$nFeature_RNA)
range(CCIObj_in_AT2LUAD$nCount_RNA)
range(CCIObj_in_AT1LUAD$percent.mt)
range(CCIObj_in_AT2LUAD$percent.mt)

##1.2. selecting cells for further analysis
# visualize QC metrics as a violin plot
VlnPlot(CCIObj_in_AT1LUAD, features = c("nFeature_RNA", "nCount_RNA","percent.mt"), group.by = c("Mouse_ID"), ncol = 3,
        log = TRUE, pt.size = 0) + NoLegend()

##2. Data normalization
CCIObj_in_AT1LUAD <- NormalizeData(CCIObj_in_AT1LUAD, normalization.method = "LogNormalize", scale.factor = 10000)
CCIObj_in_AT2LUAD <- NormalizeData(CCIObj_in_AT2LUAD, normalization.method = "LogNormalize", scale.factor = 10000)

##3. Identification of highly variable features
CCIObj_in_AT1LUAD <- FindVariableFeatures(CCIObj_in_AT1LUAD, selection.method = "vst", nfeatures = 2000) #vst = variance stablizing transformatin
CCIObj_in_AT2LUAD <- FindVariableFeatures(CCIObj_in_AT2LUAD, selection.method = "vst", nfeatures = 2000) #vst = variance stablizing transformatin

##4. Data scaling
CCIObj_in_AT1LUAD <- ScaleData(object = CCIObj_in_AT1LUAD) #2000 identified variable features
CCIObj_in_AT2LUAD <- ScaleData(object = CCIObj_in_AT2LUAD)

CCIObj_in_AT1LUAD@assays[["RNA"]]@scale.data
View(CCIObj_in_AT1LUAD@commands)


###Perform PCA, linear dimensional reduction, on the scaled data (linear dimensional reduction)
CCIObj_in_AT1LUAD <- RunPCA(CCIObj_in_AT1LUAD, features = VariableFeatures(object = CCIObj_in_AT1LUAD))
CCIObj_in_AT2LUAD <- RunPCA(CCIObj_in_AT2LUAD, features = VariableFeatures(object = CCIObj_in_AT2LUAD))
# generate 5 PCs, first 30 features in each PC

#more direct way: ElbowPlot()
ElbowPlot(CCIObj_in_AT1LUAD) #default = 20 PC
ElbowPlot(CCIObj_in_AT1LUAD, ndims = 50, reduction = "pca") #dims = 20
ElbowPlot(CCIObj_in_AT2LUAD) #default = 20 PC
ElbowPlot(CCIObj_in_AT2LUAD, ndims = 50, reduction = "pca") #dims = 20


### run Harmoney -----------------
CCIObj_in_AT1LUAD <- CCIObj_in_AT1LUAD %>%
  RunHarmony(group.by.vars = 'Batch', plot_covergence = FALSE)

CCIObj_in_AT1LUAD@reductions

CCIObj_in_AT1LUAD.embed <- Embeddings(CCIObj_in_AT1LUAD, "harmony")
CCIObj_in_AT1LUAD.embed[1:10,1:10]

CCIObj_in_AT2LUAD <- CCIObj_in_AT2LUAD %>%
  RunHarmony(group.by.vars = 'Batch', plot_covergence = FALSE)

CCIObj_in_AT2LUAD@reductions

CCIObj_in_AT2LUAD.embed <- Embeddings(CCIObj_in_AT2LUAD, "harmony")
CCIObj_in_AT2LUAD.embed[1:10,1:10]
# Do UMAP and clustering using ** Harmony embeddings instead of PCA **
CCIObj_in_AT1LUAD <- CCIObj_in_AT1LUAD %>%
  RunUMAP(reduction = 'harmony', dims = 1:20) %>%
  FindNeighbors(reduction = 'harmony', dims = 1:20) %>%
  FindClusters(resolution = 0.7) #12 clusters

CCIObj_in_AT2LUAD <- CCIObj_in_AT2LUAD %>%
  RunUMAP(reduction = 'harmony', dims = 1:20) %>%
  FindNeighbors(reduction = 'harmony', dims = 1:20) %>%
  FindClusters(resolution = 0.5) #10 clusters


CCIObj_in_AT1LUAD$seurat_clusters <- as.factor(as.numeric(as.character(CCIObj_in_AT1LUAD$seurat_clusters)) +1)
CCIObj_in_AT2LUAD$seurat_clusters <- as.factor(as.numeric(as.character(CCIObj_in_AT2LUAD$seurat_clusters)) +1)


DimPlot(CCIObj_in_AT1LUAD, reduction = "umap", label = T, repel = TRUE)
DimPlot(CCIObj_in_AT1LUAD, reduction = "umap", label = T, group.by = "Batch", repel = TRUE)
DimPlot(CCIObj_in_AT1LUAD, reduction = "umap", label = T, group.by = "seurat_clusters", repel = TRUE)
DimPlot(CCIObj_in_AT1LUAD, reduction = "umap", label = FALSE, group.by = "Mouse_ID")
DimPlot(CCIObj_in_AT1LUAD, reduction = "umap", label = T, group.by = "predicted.celltype_level3", repel = TRUE)
DimPlot(CCIObj_in_AT1LUAD, reduction = "umap", label = T, group.by = "Celltypes", repel = TRUE)
FeaturePlot()

DimPlot(CCIObj_in_AT2LUAD, reduction = "umap", label = T, repel = TRUE)
DimPlot(CCIObj_in_AT2LUAD, reduction = "umap", label = T, group.by = "Batch", repel = TRUE)
DimPlot(CCIObj_in_AT2LUAD, reduction = "umap", label = T, group.by = "seurat_clusters", repel = TRUE)
DimPlot(CCIObj_in_AT2LUAD, reduction = "umap", label = FALSE, group.by = "Mouse_ID")
DimPlot(CCIObj_in_AT2LUAD, reduction = "umap", label = T, group.by = "predicted.celltype_level3", repel = TRUE)
DimPlot(CCIObj_in_AT2LUAD, reduction = "umap", label = T, group.by = "Celltypes", repel = TRUE)

FeaturePlot(CCIObj_in_AT1LUAD, features = c("EGFP"), cols = c("lightgrey", "blue"))
FeaturePlot(CCIObj_in_AT2LUAD, features = c("EGFP"), cols = c("lightgrey", "blue"))


### combine celltypes of AECs and myeloid cells into one metadata column and then replace active.ident with it
### do CCIObj_in_AT1LUAD firstly
# to get barcode information first
Barcodes <- CCIObj_in_AT1LUAD@active.ident
Barcodes <- as.data.frame(Barcodes)
Barcodes$RowName <- rownames(Barcodes)
names(Barcodes)[1] <- "cluster"
names(Barcodes)[2] <- "Barcode"
CCIObj_in_AT1LUAD <- AddMetaData(CCIObj_in_AT1LUAD, Barcodes$Barcode, col.name = "Barcode")

# to generate a dataframe incorporting celltypes of AECs and myeloid cells
af <- FetchData(CCIObj_in_AT1LUAD, c("Celltypes", "AECcelltypes", "Barcode")) %>%
  # filter(cell_id %in% cell_subset) %>%
  mutate(cell_id = str_replace_all(Barcode, ":", "_")) %>% 
  select(cell_id, Celltypes, AECcelltypes) %>% 
  as_tibble
write.csv(af, file = "/Users/m.y./Documents/R/R_working directory/AT1LUAD_scRNA/CCI/LIANA/CCIObj_in_AT1LUAD_Annotation.csv")
af <- af %>%
  mutate(CombinedTypes = paste0(ifelse(is.na(Celltypes), "", Celltypes), ifelse(is.na(AECcelltypes), "", AECcelltypes)))
head(af)

# Create a dataframe with the CombinedTypes column and appropriate row names
new_metadata <- af[, c("cell_id", "CombinedTypes"), drop = FALSE]
head(new_metadata)

# Add the new metadata column to the Seurat object
CCIObj_in_AT1LUAD <- AddMetaData(CCIObj_in_AT1LUAD, metadata = new_metadata, col.name = "CombinedTypes")
DimPlot(CCIObj_in_AT1LUAD, reduction = "umap", label = T, group.by = "CombinedTypes", repel = TRUE)
Idents(CCIObj_in_AT1LUAD) <- CCIObj_in_AT1LUAD$CombinedTypes
DimPlot(CCIObj_in_AT1LUAD, reduction = "umap", label = T, repel = TRUE)


### do CCIObj_in_AT2LUAD secondly
# to get barcode information first
Barcodes <- CCIObj_in_AT2LUAD@active.ident
Barcodes <- as.data.frame(Barcodes)
Barcodes$RowName <- rownames(Barcodes)
names(Barcodes)[1] <- "cluster"
names(Barcodes)[2] <- "Barcode"
CCIObj_in_AT2LUAD <- AddMetaData(CCIObj_in_AT2LUAD, Barcodes$Barcode, col.name = "Barcode")

# to generate a dataframe incorporting celltypes of AECs and myeloid cells
af2 <- FetchData(CCIObj_in_AT2LUAD, c("Celltypes", "AECcelltypes", "Barcode")) %>%
  # filter(cell_id %in% cell_subset) %>%
  mutate(cell_id = str_replace_all(Barcode, ":", "_")) %>% 
  select(cell_id, Celltypes, AECcelltypes) %>% 
  as_tibble

write.csv(af, file = "/Users/m.y./Documents/R/R_working directory/AT1LUAD_scRNA/CCI/LIANA/CCIObj_in_AT2LUAD_Annotation.csv")
af2 <- af2 %>%
  mutate(CombinedTypes = paste0(ifelse(is.na(Celltypes), "", Celltypes), ifelse(is.na(AECcelltypes), "", AECcelltypes)))
head(af2)

# Create a dataframe with the CombinedTypes column and appropriate row names
new_metadata2 <- af2[, c("cell_id", "CombinedTypes"), drop = FALSE]
head(new_metadata2)

# Add the new metadata column to the Seurat object
CCIObj_in_AT2LUAD <- AddMetaData(CCIObj_in_AT2LUAD, metadata = new_metadata2, col.name = "CombinedTypes")
DimPlot(CCIObj_in_AT2LUAD, reduction = "umap", label = T, group.by = "CombinedTypes", repel = TRUE)
Idents(CCIObj_in_AT2LUAD) <- CCIObj_in_AT2LUAD$CombinedTypes
DimPlot(CCIObj_in_AT2LUAD, reduction = "umap", label = T, repel = TRUE)

# save the updated objects; from here, you are ready to perform Cell-cell communication (CCC) via using LIANA package
saveRDS(CCIObj_in_AT1LUAD, file = "/Users/m.y./Documents/R/R_working directory/AT1LUAD_scRNA/CCIObj_in_AT1LUAD.RDS")
saveRDS(CCIObj_in_AT2LUAD, file = "/Users/m.y./Documents/R/R_working directory/AT1LUAD_scRNA/CCIObj_in_AT2LUAD.RDS")
```
