## Supplemental File 3 for "Cell of origin alters myeloid-mediated immunosuppression in lung adenocarcinoma": Supplemental file_4 Trajectory analysis.ipynb

```
# %% [markdown]
# R code to convert seurat to h5ad

# %% [raw]
library(Seurat)
library(SeuratObject)
library(SeuratDisk)
library(ggplot2)
library(scCustomize)
library(dplyr)
library(pheatmap)

packageVersion("Seurat") ## might need version 4 (but 5 is no longer on beta soit probably works)
# [1] ‘5.0.3’
packageVersion("SeuratObject")
# [1] ‘5.0.1’
packageVersion('SeuratDisk')
# [1] ‘0.0.0.9021’

### load seurat object #######################################################
seurat.obj<-readRDS(file="data/Myeloid_Annotated.RDS")

### explore ##################################################################
seurat.obj
# An object of class Seurat 
# 17602 features across 19092 samples within 1 assay 
# Active assay: RNA (17602 features, 2000 variable features)
# 3 layers present: counts, data, scale.data
# 3 dimensional reductions calculated: pca, harmony, umap

seurat.obj@version
# [1] ‘5.0.1’

seurat.obj@assays$RNA
# $RNA
# Assay data with 17602 features for 19092 cells
# Top 10 variable features:
#   Inhba, Cxcl2, Apoe, Mmp12, Ttn, Rtkn2, Igkc, Slc7a11, Tg, Cxcl10 


### fix column str ##########################################################
#DimPlot(seurat.obj,group.by = 'Celltypes')
seurat.obj<-RenameIdents(seurat.obj,"Monocytes" = "Macrophages")
seurat.obj$Celltypes <- seurat.obj@active.ident
table(seurat.obj$Celltypes)

DimPlot(seurat.obj,group.by = 'seurat_clusters')
seurat.obj$MY_anno<-seurat.obj$seurat_clusters
# DimPlot(seurat.obj,group.by = 'MY_anno')
Idents(seurat.obj)<-seurat.obj$MY_anno
# DimPlot(seurat.obj)
seurat.obj<-RenameIdents(seurat.obj,"1"="MDSCs")
seurat.obj<-RenameIdents(seurat.obj,"2"="Macrophage")
seurat.obj<-RenameIdents(seurat.obj,"3"="Macrophage")
seurat.obj<-RenameIdents(seurat.obj,"4"="Macrophage")
seurat.obj<-RenameIdents(seurat.obj,"5"="TAMs")
seurat.obj<-RenameIdents(seurat.obj,"6"="Macrophage")
seurat.obj<-RenameIdents(seurat.obj,"7"="Macrophage")
seurat.obj<-RenameIdents(seurat.obj,"8"="Macrophage")
seurat.obj<-RenameIdents(seurat.obj,"9"="Macrophage")
seurat.obj<-RenameIdents(seurat.obj,"10"="Macrophage")
seurat.obj<-RenameIdents(seurat.obj,"11"="Proliferating Myeloid Cells")
seurat.obj<-RenameIdents(seurat.obj,"12"="HSPCs")
seurat.obj<-RenameIdents(seurat.obj,"13"="Proliferating Myeloid Cells")
seurat.obj<-RenameIdents(seurat.obj,"14"="Macrophage")
seurat.obj<-RenameIdents(seurat.obj,"15"="DCs")
seurat.obj<-RenameIdents(seurat.obj,"16"="Macrophage")
seurat.obj<-RenameIdents(seurat.obj,"17"="iMONs")
seurat.obj<-RenameIdents(seurat.obj,"18"="Macrophage")
seurat.obj<-RenameIdents(seurat.obj,"19"="Macrophage")
seurat.obj<-RenameIdents(seurat.obj,"20"="DCs")
seurat.obj<-RenameIdents(seurat.obj,"21"="HSPCs")
seurat.obj<-RenameIdents(seurat.obj,"22"="Granulocytes")
seurat.obj$MY_anno <- seurat.obj@active.ident
DimPlot(seurat.obj, group.by=c('MY_anno','Mouse_ID'))

seurat.obj$macro_anno<-seurat.obj$seurat_clusters
celltype_cluster<-as.data.frame.matrix(table(seurat.obj$Cell_of_Origin,seurat.obj$macro_anno))
pheatmap(celltype_cluster,scale='column')

# create the fixed_CoO
seurat.obj$fixed_CoO<-seurat.obj$Cell_of_Origin
seurat.obj$fixed_CoO<-as.character(seurat.obj$fixed_CoO)
table(seurat.obj$fixed_CoO)
seurat.obj$fixed_CoO[seurat.obj$fixed_CoO=='Neg_Controls']<-'Non-LUAD'
seurat.obj$fixed_CoO[seurat.obj$fixed_CoO=='Vehicle_Controls']<-'Non-LUAD'
seurat.obj$fixed_CoO[seurat.obj$fixed_CoO=='Spontaneous_AT2-derived LUAD']<-NA
table(seurat.obj$fixed_CoO, useNA = 'always')
table(seurat.obj$fixed_CoO, seurat.obj$Cell_of_Origin, useNA='always')

DimPlot(seurat.obj, group.by=c('seurat_clusters'),split.by='fixed_CoO',cols=polychrome_pal,pt.size=1,order=T,ncol=2, label = T)

## annotate macro_anno
celltype_cluster<-as.data.frame.matrix(table(seurat.obj$macro_anno,seurat.obj$fixed_CoO))
pheatmap(celltype_cluster,scale='column')

chisq <- chisq.test(celltype_cluster)
chisq

chisq$observed
round(chisq$expected,2)
round(chisq$residuals, 3)

pheatmap(chisq$residuals, color=c('#1a1a1a','#bababa','#fddbc7','#b2182b'),breaks=c(-20,-10,0,10,20))

Idents(seurat.obj)<-seurat.obj$macro_anno
seurat.obj<-RenameIdents(seurat.obj,"1"="MDSCs")
seurat.obj<-RenameIdents(seurat.obj,"2"="Macrophage-AT2")
seurat.obj<-RenameIdents(seurat.obj,"3"="Macrophage")
seurat.obj<-RenameIdents(seurat.obj,"4"="Macrophage")
seurat.obj<-RenameIdents(seurat.obj,"5"="TAMs")
seurat.obj<-RenameIdents(seurat.obj,"6"="Macrophage-AT1")
seurat.obj<-RenameIdents(seurat.obj,"7"="Macrophage")
seurat.obj<-RenameIdents(seurat.obj,"8"="Macrophage")
seurat.obj<-RenameIdents(seurat.obj,"9"="Macrophage-Neg")
seurat.obj<-RenameIdents(seurat.obj,"10"="Macrophage")
seurat.obj<-RenameIdents(seurat.obj,"11"="Proliferating Myeloid Cells")
seurat.obj<-RenameIdents(seurat.obj,"12"="HSPCs")
seurat.obj<-RenameIdents(seurat.obj,"13"="Proliferating Myeloid Cells")
seurat.obj<-RenameIdents(seurat.obj,"14"="Macrophage")
seurat.obj<-RenameIdents(seurat.obj,"15"="DCs")
seurat.obj<-RenameIdents(seurat.obj,"16"="Macrophage")
seurat.obj<-RenameIdents(seurat.obj,"17"="iMONs")
seurat.obj<-RenameIdents(seurat.obj,"18"="Macrophage")
seurat.obj<-RenameIdents(seurat.obj,"19"="Macrophage")
seurat.obj<-RenameIdents(seurat.obj,"20"="DCs")
seurat.obj<-RenameIdents(seurat.obj,"21"="HSPCs")
seurat.obj<-RenameIdents(seurat.obj,"22"="Granulocytes")
seurat.obj$macro_anno <- seurat.obj@active.ident
DimPlot(seurat.obj, group.by=c('macro_anno'),split.by='Cell_of_Origin',cols=polychrome_pal,pt.size=1,order=T,ncol=2, label = T)
DimPlot(seurat.obj, group.by=c('macro_anno'),split.by='fixed_CoO',cols=polychrome_pal,pt.size=1,order=T,ncol=2, label = T)

str(seurat.obj@meta.data)
seurat.obj$Cell_of_Origin<-as.character(seurat.obj$Cell_of_Origin)
seurat.obj$fixed_CoO<-as.character(seurat.obj$fixed_CoO)
seurat.obj$Celltypes<-as.character(seurat.obj$Celltypes)
seurat.obj$seurat_clusters<-as.character(seurat.obj$seurat_clusters)
seurat.obj$MY_anno<-as.character(seurat.obj$MY_anno)
seurat.obj$macro_anno<-as.character(seurat.obj$macro_anno)
str(seurat.obj@meta.data)

DimPlot(seurat.obj,group.by=c('Mouse_ID','seurat_clusters','MY_anno','Cell_of_Origin','macro_anno'),cols=polychrome_pal,pt.size=0.5,order=T,ncol=2)

#### explore data set #######################################################
DimPlot(seurat.obj,group.by=c('Celltypes','predicted.celltype_level3'))

table_compare<-as.data.frame.matrix(table(seurat.obj$predicted.celltype_level3, seurat.obj$Celltypes))
pheatmap(table_compare,scale='column')

MY_markers<-c('Mmp12','Arg1','Chil3','Chil4', # TAMs
              'H2-Ab1','H2-Aa','H2-Eb1','Cst3') # DC

myeloid_markers<-c('Chil3','Atp6v0d2','Plet1','Ctsd', # Alv Macrophage
                   'G0s2','Asprv1','S100a9','Cstdc4', # Neut
                   'Plac8','Ifitm3','Ifi27l2a','Pou2f2','S100a4', # Mono
                   'Pf4','C1qa','C1qb','C1qc','Apoe', # Int Macrophage
                   'Camp','Ngp','Ltf','Lcn2','Stfa3', # Neur
                   'Il6','Mcpt8','Ccl4','Ifitm1','Ccl3', # Mast Ba2
                   'H2-Ab1','H2-Aa','H2-Eb1','Cd74','Cst3', # DC1
                   'Ccl5','Ccr7','Fscn1','Traf1','Epsti1') # DC2

DoHeatmap(seurat.obj, features=MY_markers, group.by='Celltypes')
DotPlot(seurat.obj, features='Cd33', group.by='MY_anno') + coord_flip()
DotPlot(seurat.obj, features='Cd33', group.by='seurat_clusters') + coord_flip()

DoHeatmap(seurat.obj, features=myeloid_markers, group.by='predicted.celltype_level3')
DoHeatmap(seurat.obj, features=myeloid_markers, group.by='seurat_clusters')


myeloid_markers <- FindAllMarkers(seurat.obj, only.pos = TRUE)

myeloid_markers %>%
  group_by(cluster) %>%
  dplyr::filter(avg_log2FC > 1)

myeloid_markers %>%
  group_by(cluster) %>%
  dplyr::filter(avg_log2FC > 1) %>%
  slice_head(n = 10) %>%
  ungroup() -> top10

DoHeatmap(seurat.obj, features = top10$gene) + NoLegend()


########### subset by cell of origin
DimPlot(seurat.obj, reduction='umap',group.by=c('seurat_clusters','MY_anno'),split.by='Cell_of_Origin',cols=polychrome_pal,pt.size=1, order=T,ncol=2, label=T)
DimPlot(seurat.obj, reduction='umap',group.by=c('Mouse_ID'),split.by='Cell_of_Origin',cols=polychrome_pal,pt.size=1, order=T,ncol=2)
DimPlot(seurat.obj, reduction='umap',split.by=c('Mouse_ID'),group.by='Cell_of_Origin',cols=polychrome_pal,pt.size=1, order=T,ncol=2)
Idents(seurat.obj)<-'Cell_of_Origin'
str(seurat.obj@meta.data)
myeloid_AT2<-subset(seurat.obj, idents="AT2-derived LUAD")
myeloid_AT1<-subset(seurat.obj,  idents="AT1-derived LUAD")
myeloid_neg<-subset(seurat.obj,  idents=c("Vehicle_Controls",'Neg_Controls'))

#### reset seurat object #####################################################
myeloid_AT2<-CreateSeuratObject(counts=myeloid_AT2@assays$RNA$counts, meta.data=myeloid_AT2@meta.data)  # change the input
myeloid_AT2[["RNA"]] <- split(myeloid_AT2[["RNA"]], f=myeloid_AT2$Mouse_ID)
myeloid_AT2

myeloid_AT1<-CreateSeuratObject(counts=myeloid_AT1@assays$RNA$counts, meta.data=myeloid_AT1@meta.data)  # change the input
myeloid_AT1[["RNA"]] <- split(myeloid_AT1[["RNA"]], f=myeloid_AT1$Mouse_ID)
myeloid_AT1

myeloid_neg<-CreateSeuratObject(counts=myeloid_neg@assays$RNA$counts, meta.data=myeloid_neg@meta.data)  # change the input
myeloid_neg[["RNA"]] <- split(myeloid_neg[["RNA"]], f=myeloid_neg$Mouse_ID)
myeloid_neg

# pre-processing
myeloid_AT2 <- NormalizeData(myeloid_AT2)
myeloid_AT2 <- FindVariableFeatures(myeloid_AT2)
myeloid_AT2 <- ScaleData(myeloid_AT2)
myeloid_AT2 <- RunPCA(myeloid_AT2)
ElbowPlot(myeloid_AT1, ndims=50) # run code below

myeloid_AT1 <- NormalizeData(myeloid_AT1)
myeloid_AT1 <- FindVariableFeatures(myeloid_AT1)
myeloid_AT1 <- ScaleData(myeloid_AT1)
myeloid_AT1 <- RunPCA(myeloid_AT1)
ElbowPlot(myeloid_AT1, ndims=50) # run code below

myeloid_neg <- NormalizeData(myeloid_neg)
myeloid_neg <- FindVariableFeatures(myeloid_neg)
myeloid_neg <- ScaleData(myeloid_neg)
myeloid_neg <- RunPCA(myeloid_neg)
ElbowPlot(myeloid_neg, ndims=50) # run code below

##########################################################################################
#### Elbow plot: quantitative approach
# https://hbctraining.github.io/scRNA-seq_online/lessons/elbow_plot_metric.html
### first metric
# Determine percent of variation associated with each PC
pct <- myeloid_AT2@reductions$pca@stdev / sum(myeloid_AT2@reductions$pca@stdev) * 100
# Calculate cumulative percents for each PC
cumu <- cumsum(pct)
# Determine which PC exhibits cumulative percent greater than 90% and % variation associated with the PC as less than 5
co1 <- which(cumu > 90 & pct < 5)[1]
co1
### second metric
# Determine the difference between variation of PC and subsequent PC
co2 <- sort(which((pct[1:length(pct) - 1] - pct[2:length(pct)]) > 0.1), decreasing = T)[1] + 1
# last point where change of % of variation is more than 0.1%.
co2
# Minimum of the two calculation
pcs <- min(co1, co2)
pcs
# Create a dataframe with values
plot_df <- data.frame(pct = pct, 
                      cumu = cumu, 
                      rank = 1:length(pct))
# Elbow plot to visualize 
ggplot(plot_df, aes(cumu, pct, label = rank, color = rank > pcs)) + 
  geom_text() + 
  geom_vline(xintercept = 90, color = "grey") + 
  geom_hline(yintercept = min(pct[pct > 5]), color = "grey") +
  theme_bw()
#############################################################################################

# integration
myeloid_AT2<- IntegrateLayers(myeloid_AT2, method = HarmonyIntegration, orig.reduction = "pca",new.reduction = "harmony")
myeloid_AT2[["RNA"]] <- JoinLayers(myeloid_AT2[["RNA"]])
myeloid_AT2

myeloid_AT2<- IntegrateLayers(myeloid_AT2, method = HarmonyIntegration, orig.reduction = "pca",new.reduction = "harmony")
myeloid_AT2[["RNA"]] <- JoinLayers(myeloid_AT2[["RNA"]])
myeloid_AT2

myeloid_AT2<- IntegrateLayers(myeloid_AT2, method = HarmonyIntegration, orig.reduction = "pca",new.reduction = "harmony")
myeloid_AT2[["RNA"]] <- JoinLayers(myeloid_AT2[["RNA"]])
myeloid_AT2

# clustering and umap
myeloid_AT2 <- FindNeighbors(myeloid_AT2, reduction = "harmony", dims = 1:20)
myeloid_AT2 <- FindClusters(myeloid_AT2, resolution=0.8, cluster.name = "harmony_clusters") 
myeloid_AT2 <- RunUMAP(myeloid_AT2 , reduction = "harmony", dims = 1:20, reduction.name = "umap")
DimPlot(myeloid_AT2, reduction='umap',group.by=c('Mouse_ID','seurat_clusters','MY_anno','fixed_CoO'),cols=polychrome_pal,pt.size=1, order=T,ncol=2)

myeloid_AT1 <- FindNeighbors(myeloid_AT1, reduction = "harmony", dims = 1:20)
myeloid_AT1 <- FindClusters(myeloid_AT1, resolution=0.8, cluster.name = "harmony_clusters") 
myeloid_AT1 <- RunUMAP(myeloid_AT1 , reduction = "harmony", dims = 1:20, reduction.name = "umap")
DimPlot(myeloid_AT1, reduction='umap',group.by=c('Mouse_ID','seurat_clusters','MY_anno','fixed_CoO'),cols=polychrome_pal,pt.size=1, order=T,ncol=2)

myeloid_neg <- FindNeighbors(myeloid_neg, reduction = "harmony", dims = 1:20)
myeloid_neg <- FindClusters(myeloid_neg, resolution=0.8, cluster.name = "harmony_clusters") 
myeloid_neg <- RunUMAP(myeloid_neg , reduction = "harmony", dims = 1:20, reduction.name = "umap")
DimPlot(myeloid_neg, reduction='umap',group.by=c('Mouse_ID','seurat_clusters','MY_anno','fixed_CoO'),cols=polychrome_pal,pt.size=1, order=T,ncol=2)


# convert to RNA3 format
myeloid_AT2[["RNA3"]] <- as(myeloid_AT2[["RNA"]], Class = "Assay")
DefaultAssay(myeloid_AT2) <- "RNA3"
myeloid_AT2[["RNA"]] <- NULL
myeloid_AT2 <- RenameAssays(myeloid_AT2, RNA3 = 'RNA')

myeloid_AT1[["RNA3"]] <- as(myeloid_AT1[["RNA"]], Class = "Assay")
DefaultAssay(myeloid_AT1) <- "RNA3"
myeloid_AT1[["RNA"]] <- NULL
myeloid_AT1 <- RenameAssays(myeloid_AT1, RNA3 = 'RNA')

myeloid_neg[["RNA3"]] <- as(myeloid_neg[["RNA"]], Class = "Assay")
DefaultAssay(myeloid_neg) <- "RNA3"
myeloid_neg[["RNA"]] <- NULL
myeloid_neg <- RenameAssays(myeloid_neg, RNA3 = 'RNA')


# save to h5ad
SaveH5Seurat(myeloid_AT2, filename = "results/myeloid_AT2.h5Seurat") # 11 pcs
Convert("results/myeloid_AT2.h5Seurat", dest = "h5ad")

SaveH5Seurat(myeloid_AT1, filename = "results/myeloid_AT1.h5Seurat") # 12 pcs
Convert("results/myeloid_AT1.h5Seurat", dest = "h5ad")

SaveH5Seurat(myeloid_neg, filename = "results/myeloid_neg.h5Seurat") # 8 pcs
Convert("results/myeloid_neg.h5Seurat", dest = "h5ad")

# %% [markdown]
# Installation of CellRank and dependencies

# %% [raw]
## per the cellrank installation instruction, use mamba is recommended for installation
## https://cellrank.readthedocs.io/en/stable/installation.html

## check to see if you have mamba
mamba -V

#mamba 1.5.8
#conda 24.3.0

## otherwise, do a clean install of it using miniforge
## https://github.com/conda-forge/miniforge

# on unix-like platforms
curl -L -O "https://github.com/conda-forge/miniforge/releases/latest/download/Miniforge3-$(uname)-$(uname -m).sh"
bash Miniforge3-$(uname)-$(uname -m).sh

## set up the Jupyter kernel for trajectory analysis
mamba create --name trajectory python=3.9.12
mamba activate trajectory

# install the conda-forge packages
mamba install -c conda-forge cellrank ipykernel

# install the pip packages
pip install moscot 'git+https://github.com/dpeerlab/Palantir' scvi-tools==1.1.1

# install scvi (it appears to be broken at the moment, the lines below should be a fix)
# pip install scvi-tools==1.1.1

# instal fa2 (it had issues but the lines below should fix it)
git clone https://github.com/bhargavchippada/forceatlas2
cd forceatlas2
pip install . --user
cd ..
rm -rf forceatlas2

## once the above packages are installed, run the line below to set up the Jupyter kernel
python -m ipykernel install --user --name trajectory --display-name "Trajectory"

# %% [code]
## packages to load
# misc package
import os
import numpy as np
import scvelo as scv
import cellrank as cr
import scanpy as sc
import scanpy.external as sce
import seaborn as sns
import pandas as pd
import magic

# CellRank packages
from cellrank.kernels import RealTimeKernel
from moscot.problems.time import TemporalProblem
from cellrank.estimators import GPCCA

# Plotting
import matplotlib
import matplotlib.pyplot as plt

# Inline plotting
%matplotlib inline

# warnings
import warnings
from numba.core.errors import NumbaDeprecationWarning

warnings.filterwarnings(action="ignore", category=NumbaDeprecationWarning)
warnings.filterwarnings(
    action="ignore", module="scanpy", message="No data for colormapping")

# settings
sc.settings.set_figure_params(frameon=False, dpi=100)
cr.settings.verbosity = 2

# %% [markdown]
# AT1 subset

# %% [code]
adata=sc.read("data/myeloid_AT1.h5ad")
adata

# %% [code]
adata.uns['macro_anno_colors']=['#cfafd4','#715ea9','#e0271e','#ffff99','#2b9cd7','#00ffef','#000080','#737ca1','#FFA500','#357337','#ff8ad7']

# %% [code]
sc.pl.embedding(adata, basis="umap",color=['MY_anno','macro_anno'], wspace=0.5)

# %% [code]
sc.pp.neighbors(adata, use_rep='X_harmony', n_neighbors=30, n_pcs=12) # 12 pcs = change of % variation is more than 0.1%

# %% [code]
# FA embedding of neighbors
sc.tl.draw_graph(adata)
sc.pl.draw_graph(adata, color=['MY_anno','macro_anno','Mouse_ID'])

# %% [code]
# FA embedding of neighbors with denoising via diffmap
sc.tl.diffmap(adata)
sc.pp.neighbors(adata,use_rep="X_diffmap",n_neighbors=30)
sc.tl.draw_graph(adata)
sc.pl.draw_graph(adata, color=['MY_anno','macro_anno'])

# %% [code]
sc.tl.leiden(adata, resolution=1)
sc.tl.paga(adata, groups="leiden")

# %% [code]
sc.pl.paga(adata, color=['leiden','MY_anno','macro_anno'], threshold=0.5)

# %% [code]
sc.pl.draw_graph(adata, color=["leiden",'MY_anno','macro_anno'], wspace=0.5)

# %% [code]
sc.pl.draw_graph(adata, color=["leiden"], legend_loc="on data", wspace=0.5)

# %% [code]
sc.tl.diffmap(macs)

# %% [code]
scv.pl.scatter(macs,basis="diffmap",color="leiden",components=["1,2"])

# %% [code]
# prolif myeloid start
macs.obsm['X_diffmap'][:, 1].argmin()

# %% [code]
root_ixs = 1517

# %% [code]
scv.pl.scatter(macs,basis="diffmap",color=["leiden",root_ixs],components=["1,2"],legend_loc="right",wspace=0.6)

# %% [code]
macs.uns["iroot"] = root_ixs

# %% [code]
sc.tl.dpt(macs)

# %% [code]
sc.pl.embedding(
    macs,
    basis="draw_graph_fa",
    color=["dpt_pseudotime"],
    color_map="gnuplot2",
)

# %% [code]
sc.pl.violin(
    macs,
    keys=["dpt_pseudotime"],
    groupby="leiden",
    rotation=90)

# %% [code]
pk = cr.kernels.PseudotimeKernel(macs, time_key="dpt_pseudotime")
pk.compute_transition_matrix()

# %% [code]
pk.plot_projection(basis="draw_graph_fa", color='macro_anno',legend_loc="right margin")

# %% [code]
pk.plot_random_walks(
    max_iter=1000,
    start_ixs={"leiden": '6'},
    basis="draw_graph_fa",
    seed=0,
    dpi=100,
    size=10)

# %% [code]
g = GPCCA(pk)
g

# %% [code]
g.compute_schur()
g.plot_spectrum(real_only=True)

# %% [code]
g.compute_macrostates(n_states=10, cluster_key="leiden")
g.plot_macrostates(which='all',basis='draw_graph_fa',s=100)

# %% [code]
## set terminal states, use the random walk plot as a guild
g.set_terminal_states(states=["5","7","11"])
g.plot_macrostates(which="terminal", legend_loc="right", basis='draw_graph_fa',s=100)

# %% [code]
g.set_initial_states(states=["6"])
g.plot_macrostates(which="initial",legend_loc="right", basis='draw_graph_fa',s=100)

# %% [code]
g.compute_fate_probabilities()
g.plot_fate_probabilities(legend_loc="right", basis='draw_graph_fa')

# %% [code]
g.plot_fate_probabilities(legend_loc="right", basis='draw_graph_fa',same_plot=False)

# %% [code]
g.plot_fate_probabilities(legend_loc="right", basis='diffmap',components=["4,2"],same_plot=False)

# %% [code]
# 11 is macrophage
mac11 = g.compute_lineage_drivers(lineages=["11"], cluster_key="leiden")
mac11.head(10)

# %% [code]
# 7 AT1-derived specific macrophage
at1_mac7 = g.compute_lineage_drivers(lineages=["7"], cluster_key="leiden")
at1_mac7.head(10)

# %% [code]
# 5 another AT1-derived specific macrophage
at1_mac5 = g.compute_lineage_drivers(lineages=["5"], cluster_key="leiden")
at1_mac5.head(10)

# %% [code]
model = cr.models.GAM(macs, link='identity')

# %% [code]
data_magic = magic.MAGIC().fit_transform(macs)

# %% [code]
macs.layers["magic"]=data_magic.X

# %% [code]
sc.pl.embedding(
    macs,
    basis="draw_graph_fa",
    color=["macro_anno"])

# %% [code]
cr.pl.heatmap(
    macs,
    model=model,
    lineages="11",
    cluster_key="macro_anno",
    show_fate_probabilities=False,
    data_key='magic',
    genes=mac11.head(10).index,
    time_key="dpt_pseudotime",
    figsize=(8,3),
    weight_threshold=(1e-3, 1e-3),
    show_all_genes=True,
    scale=True)

# %% [code]
cr.pl.heatmap(
    macs,
    model=model,
    lineages="7",
    cluster_key="macro_anno",
    show_fate_probabilities=False,
    data_key='magic',
    genes=at1_mac7.head(10).index,
    time_key="dpt_pseudotime",
    figsize=(8,3),
    weight_threshold=(1e-3, 1e-3),
    show_all_genes=True,
    scale=True)

# %% [code]
cr.pl.heatmap(
    macs,
    model=model,
    lineages="5",
    cluster_key="macro_anno",
    show_fate_probabilities=False,
    data_key='magic',
    genes=at1_mac5.head(10).index,
    time_key="dpt_pseudotime",
    figsize=(8,3),
    weight_threshold=(1e-3, 1e-3),
    show_all_genes=True,
    scale=True)

# %% [markdown]
# AT2 subset

# %% [code]
adata=sc.read("data/myeloid_AT2.h5ad")
adata

# %% [code]
# fixed colors
adata.uns['macro_anno_colors']=['#cfafd4','#715ea9','#e0271e','#ffff99','#2b9cd7','#00ffef','#000080','#737ca1','#FFA500','#357337','#ff8ad7']

# %% [code]
sc.pl.embedding(adata, basis="umap",color=['MY_anno','macro_anno'], wspace=0.6)

# %% [code]
sc.pp.neighbors(adata, use_rep='X_harmony', n_neighbors=30, n_pcs=11) # 11 pcs = change of % variation is more than 0.1%

# %% [code]
# FA embedding of neighbors
sc.tl.draw_graph(adata)
sc.pl.draw_graph(adata, color=['MY_anno','macro_anno','Mouse_ID'])

# %% [code]
# FA embedding of neighbors with denoising via diffmap
sc.tl.diffmap(adata)
sc.pp.neighbors(adata,use_rep="X_diffmap",n_neighbors=30)
sc.tl.draw_graph(adata)
sc.pl.draw_graph(adata, color=['MY_anno','macro_anno'])

# %% [code]
sc.tl.leiden(adata, resolution=1.5)
sc.tl.paga(adata, groups="leiden")

# %% [code]
sc.pl.paga(adata, color=['leiden','MY_anno','macro_anno'], threshold=0.5)

# %% [code]
sc.pl.paga(adata, color=['leiden','MY_anno','macro_anno'], threshold=0.3)

# %% [code]
sc.pl.draw_graph(adata, color=["leiden",'MY_anno','macro_anno'], wspace=0.5)

# %% [code]
sc.pl.draw_graph(adata, color=["leiden"], legend_loc="on data", wspace=0.5)

# %% [code]
sc.tl.diffmap(macs)

# %% [code]
with plt.rc_context({"figure.figsize": (8,8)}):
    scv.pl.scatter(macs,basis="diffmap",color="leiden",components=["1,2"])

# %% [code]
# macrophage start
macs.obsm['X_diffmap'][:, 2].argmax()

# %% [code]
root_ixs = 552

# %% [code]
scv.pl.scatter(macs,basis="diffmap",color=["leiden",root_ixs],components=["1,2"],legend_loc="right",wspace=0.6)

# %% [code]
macs.uns["iroot"] = root_ixs

# %% [code]
sc.tl.dpt(macs)

# %% [code]
sc.pl.embedding(
    macs,
    basis="draw_graph_fa",
    color=["dpt_pseudotime"],
    color_map="gnuplot2",
)

# %% [code]
sc.pl.violin(
    macs,
    keys=["dpt_pseudotime"],
    groupby="leiden",
    rotation=90)

# %% [code]
pk = cr.kernels.PseudotimeKernel(macs, time_key="dpt_pseudotime")
pk.compute_transition_matrix()

# %% [code]
pk.plot_projection(basis="draw_graph_fa", color='macro_anno',legend_loc="right margin")

# %% [code]
pk.plot_random_walks(
    max_iter=500,
    start_ixs={"leiden": '32'},
    basis="draw_graph_fa",
    seed=0,
    dpi=100,
    size=10)

# %% [code]
g = GPCCA(pk)
g

# %% [code]
g.compute_schur()
g.plot_spectrum(real_only=True)

# %% [code]
g.compute_macrostates(n_states=12, cluster_key="leiden")
g.plot_macrostates(which='all',basis='draw_graph_fa',s=100)

# %% [code]
## set terminal states, use the random walk plot as a guild
g.set_terminal_states(states=["9","27","0"])
g.plot_macrostates(which="terminal", legend_loc="right", basis='draw_graph_fa',s=100)

# %% [code]
g.set_initial_states(states=["32"])
g.plot_macrostates(which="initial",legend_loc="right", basis='draw_graph_fa',s=100)

# %% [code]
g.compute_fate_probabilities()
g.plot_fate_probabilities(legend_loc="right", basis='draw_graph_fa')

# %% [code]
g.plot_fate_probabilities(legend_loc="right", basis='draw_graph_fa',same_plot=False)

# %% [code]
# 9 is TAMs/AT2 macrophage
TAMs9_df = g.compute_lineage_drivers(lineages=["9"], cluster_key="leiden")
TAMs9_df.head(10)

# %% [code]
# 27 is MDSCs
MDSCs27_df = g.compute_lineage_drivers(lineages=["27"], cluster_key="leiden")
MDSCs27_df.head(10)

# %% [code]
# 0 is another set of TAMs/AT2 macrophage
TAMs0_df = g.compute_lineage_drivers(lineages=["0"], cluster_key="leiden")
TAMs0_df.head(10)

# %% [code]
model = cr.models.GAM(macs, link='identity')
model

# %% [code]
data_magic = magic.MAGIC().fit_transform(macs)

# %% [code]
macs.layers["magic"]=data_magic.X

# %% [code]
sc.pl.embedding(
    macs,
    basis="draw_graph_fa",
    color=["macro_anno"],
)

# %% [code]
cr.pl.heatmap(
    macs,
    model=model,
    lineages="9",
    cluster_key="macro_anno",
    show_fate_probabilities=False,
    data_key='magic',
    genes=TAMs9_df.head(10).index,
    time_key="dpt_pseudotime",
    figsize=(8,3),
    weight_threshold=(1e-3, 1e-3),
    show_all_genes=True,
    scale=True)

# %% [code]
cr.pl.heatmap(
    macs,
    model=model,
    lineages="27",
    cluster_key="macro_anno",
    show_fate_probabilities=False,
    data_key='magic',
    genes=MDSCs27_df.head(10).index,
    time_key="dpt_pseudotime",
    figsize=(8,3),
    weight_threshold=(1e-3, 1e-3),
    show_all_genes=True,
    scale=True)

# %% [code]
cr.pl.heatmap(
    macs,
    model=model,
    lineages="0",
    cluster_key="macro_anno",
    show_fate_probabilities=False,
    data_key='magic',
    genes=TAMs0_df.head(10).index,
    time_key="dpt_pseudotime",
    figsize=(8,3),
    weight_threshold=(1e-3, 1e-3),
    show_all_genes=True,
    scale=True)

# %% [markdown]
# Negative and vehicle control subset

# %% [code]
adata=sc.read("data/myeloid_neg.h5ad")
adata

# %% [code]
adata.uns['macro_anno_colors']=['#cfafd4','#715ea9','#e0271e','#2b9cd7','#82caff','#737ca1','#FFA500','#357337','#ff8ad7']

# %% [code]
sc.pl.embedding(adata, basis="umap",color=['MY_anno','macro_anno'], wspace=0.5)

# %% [code]
sc.pp.neighbors(adata, use_rep='X_harmony', n_neighbors=30, n_pcs=8)  # 8 pcs = change of % variation is more than 0.1%

# %% [code]
# FA embedding of neighbors
sc.tl.draw_graph(adata)
sc.pl.draw_graph(adata, color=['MY_anno','macro_anno','Mouse_ID'])

# %% [code]
# FA embedding of neighbors with denoising via diffmap
sc.tl.diffmap(adata)
sc.pp.neighbors(adata,use_rep="X_diffmap",n_neighbors=30)
sc.tl.draw_graph(adata)
sc.pl.draw_graph(adata, color=['MY_anno','macro_anno'])

# %% [code]
sc.pl.draw_graph(adata, color=['MY_anno','macro_anno'], wspace=0.6)

# %% [code]
sc.tl.leiden(adata, resolution=1)
sc.tl.paga(adata, groups="leiden")

# %% [code]
sc.pl.paga(adata, color=['leiden','MY_anno','macro_anno'], threshold=0.4)

# %% [code]
sc.pl.draw_graph(adata, color=["leiden",'MY_anno','macro_anno'], wspace=0.5)

# %% [code]
sc.pl.draw_graph(adata, color=["leiden"], legend_loc="on data", wspace=0.5)

# %% [code]
macs=adata[adata.obs['leiden'].isin(['2','4','14','12','3','1','18','13','0',
                                    '5','11','8','17'])].copy()

# %% [code]
sc.pl.draw_graph(macs, color=["leiden"], legend_loc="on data", wspace=0.5)

# %% [code]
sc.pl.draw_graph(macs, color=["macro_anno"], wspace=0.5)

# %% [code]
sc.tl.diffmap(macs)

# %% [code]
scv.pl.scatter(macs,basis="diffmap",color="leiden",components=["1,2"])

# %% [code]
# macrophage start
macs.obsm['X_diffmap'][:, 1].argmin()

# %% [code]
root_ixs = 1464

# %% [code]
scv.pl.scatter(macs,basis="diffmap",color=["leiden",root_ixs],components=["1,2"],legend_loc="right",wspace=0.6)

# %% [code]
macs.uns["iroot"] = root_ixs

# %% [code]
sc.tl.dpt(macs)

# %% [code]
sc.pl.embedding(
    macs,
    basis="draw_graph_fa",
    color=["dpt_pseudotime"],
    color_map="gnuplot2",
)

# %% [code]
sc.pl.violin(
    macs,
    keys=["dpt_pseudotime"],
    groupby="leiden",
    rotation=90)

# %% [code]
pk = cr.kernels.PseudotimeKernel(macs, time_key="dpt_pseudotime")
pk.compute_transition_matrix()

# %% [code]
pk.plot_projection(basis="draw_graph_fa", color='macro_anno',legend_loc="right margin")

# %% [code]
pk.plot_random_walks(
    max_iter=1000,
    start_ixs={"leiden": '4'},
    basis="draw_graph_fa",
    seed=1)

# %% [code]
g = GPCCA(pk)
g

# %% [code]
g.compute_schur()
g.plot_spectrum(real_only=True)

# %% [code]
g.compute_macrostates(n_states=4, cluster_key="leiden")
g.plot_macrostates(which='all',basis='draw_graph_fa',s=100)

# %% [code]
## set terminal states, use the random walk plot as a guild
g.set_terminal_states(states=['17','14'])
g.plot_macrostates(which="terminal", legend_loc="right", basis='draw_graph_fa',s=100)

# %% [code]
g.set_initial_states(states=["4"])
g.plot_macrostates(which="initial",legend_loc="right", basis='draw_graph_fa',s=100)

# %% [code]
g.compute_fate_probabilities()
g.plot_fate_probabilities(legend_loc="right", basis='draw_graph_fa')

# %% [code]
g.plot_fate_probabilities(legend_loc="right", basis='draw_graph_fa',same_plot=False)

# %% [code]
# 17 is macrophages
macs17 = g.compute_lineage_drivers(lineages=['17'], cluster_key="leiden")
macs17.head(10)

# %% [code]
model = cr.models.GAM(macs, link='identity')

# %% [code]
data_magic = magic.MAGIC().fit_transform(macs)

# %% [code]
macs.layers["magic"]=data_magic.X

# %% [code]
# fixes
macs.uns['macro_anno_colors']=['#cfafd4','#e0271e','#2b9cd7','#82caff','#737ca1','#FFA500','#357337','#ff8ad7']

# %% [code]
sc.pl.embedding(
    macs,
    basis="draw_graph_fa",
    color=["macro_anno"],
)

# %% [code]
cr.pl.heatmap(
    macs,
    model=model,
    lineages="17",
    cluster_key="macro_anno",
    show_fate_probabilities=False,
    data_key='magic',
    genes=macs17.head(10).index,
    time_key="dpt_pseudotime",
    figsize=(8,3),
    weight_threshold=(1e-3, 1e-3),
    show_all_genes=True,
    scale=True)
```
