## Supplemental File 6 for "Cell of origin alters myeloid-mediated immunosuppression in lung adenocarcinoma": Supplemental File 7_CCC_analysis via LIANA.Rmd

````
---
title: "LIANA Analysis"
author:
  - name: Daniel Dimitrov
    affiliation:
        - Saezlab, Heidelberg University
   
date: "`r Sys.Date()`"
output: 
  BiocStyle::html_document:
      self_contained: true
      toc: true
      toc_float: true
      toc_depth: 3
      code_folding: show
vignette: >
  %\VignetteIndexEntry{LIANA Tutorial}
  %\VignetteEngine{knitr::rmarkdown}
  %\VignetteEncoding{UTF-8}
---

```{r chunk_setup, include = FALSE}
knitr::opts_chunk$set(
    collapse = TRUE,
    comment = "#>"
)
```

# `liana`: Intro

The continuous developments of single-cell RNA-Seq (scRNA-Seq) have sparked an
immense interest in understanding intercellular crosstalk. Multiple tools and
resources that aid the investigation of cell-cell communication (CCC) were 
published recently. However, these methods and resources are usually in a fixed
combination of a tool and its corresponding resource, but in principle any
resource could be combined with any method.

To this end, we built `liana` - a framework to decouple the tools from their
corresponding resources.

![](ligrec_pipe.png)

### install LIANA package 
```{r}
if (!requireNamespace("BiocManager", quietly = TRUE))
    install.packages("BiocManager")

if (!requireNamespace("remotes", quietly = TRUE))
    install.packages("remotes")

remotes::install_github('saezlab/liana')
```


## Load required packages
```{r load_pkgs, message=FALSE, warning=FALSE}
library(scater)
library(tidyverse)
library(magrittr)
library(liana)
library(circlize)
```

## CCC Resources
`liana` provides CCC resources obtained and formatted via [`OmnipathR`](https://github.com/saezlab/OmnipathR) which are then converted to the appropriate format to each method.
```{r liana_resource, warning = FALSE}
# Resource currently included in OmniPathR (and hence `liana`) include:
show_resources()

# A list of resources can be obtained using the `select_resource()` function:
# See `?select_resource()` documentation for further information.
# select_resource(c('OmniPath')) %>% dplyr::glimpse() 
```
  
  
## CCC Methods
Each of the resources can then be run with any of the following methods:
```{r liana_method, warning = FALSE}
# Resource currently included in OmniPathR (and hence `liana`) include:
show_methods()
```
Note that the different algorithms (or scoring measures) used in `sca`, `natmi`,
`connectome`, `cellphonedb`, `cytotalk`'s crosstalk scores, and `logfc` were re-implemented in LIANA.
Yet, the original method pipelines can be called via the `call_*` functions.

## `liana` wrapper function

To run `liana`, we will use a down-sampled toy *HUMAN* PBMCs scRNA-Seq data set, obtained from [SeuratData](https://github.com/satijalab/seurat-data).


`liana` takes `Seurat` and `SingleCellExperiment` objects as input, containing processed counts and clustered cells.
```{r load_data}
liana_path <- system.file(package = "liana")
testdata <- readRDS("your/file/path/CCIObj_in_AT1LUAD.RDS")

testdata %>% dplyr::glimpse()
```


```{r load_data}
liana_path <- system.file(package = "liana")
testdata2 <- readRDS("your/file/path/CCIObj_in_AT2LUAD.RDS")

testdata2 %>% dplyr::glimpse()
```

`liana_wrap` calls a number of methods and and each method is run with the provided resource(s).

We will now call all methods that are currently available in liana.

Here we use only the `Consensus` (Default) CCC resource, but any of the 
aforementioned ones (available via `show_resources()`) can be added to the `resource` parameter
```{r liana_run, message = FALSE, print = FALSE, results='hide'}
# Run liana
liana_test <- liana_wrap(testdata,
                         resource = c("MouseConsensus"),
                         idents_col = NULL,
                         min_cells = 5,
                         return_all = FALSE,
                         supp_columns = NULL,
                         verbose = TRUE,
                         assay = NULL,
                         .simplify = TRUE,
                         cell.adj = NULL,
                         base = NULL)


# Liana returns a list of results, each element of which corresponds to a method
liana_test %>% dplyr::glimpse()
```


```{r liana_run, message = FALSE, print = FALSE, results='hide'}
# Run liana
liana_test2 <- liana_wrap(testdata2,
                         resource = c("MouseConsensus"),
                         idents_col = NULL,
                         min_cells = 5,
                         return_all = FALSE,
                         supp_columns = NULL,
                         verbose = TRUE,
                         assay = NULL,
                         .simplify = TRUE,
                         cell.adj = NULL,
                         base = NULL)


# Liana returns a list of results, each element of which corresponds to a method
liana_test2 %>% dplyr::glimpse()
```

LIANA currently provides a mixture of re-implemented methods and pipelines which externally call specific LR methods.
By default, LIANA will call *only the internal scoring function*, i.e. those that are re-implemented in LIANA.

One can use LIANA to also run the original methods.
For more about the original methods see [LIANA++](https://saezlab.github.io/liana/articles/liana_devel.html).


## Aggregate and Obiain Consensus Ranks
`liana` also provides consensus ranks for the results obtained using different
methods. By default, `liana` will provide mean, median, and aggregate* consensus
ranks

# run through the code below line by line!!!
```{r liana_ranks, warning=FALSE}
# We can aggregate these results into a tibble with consensus ranks
liana_test <- liana_test %>%
  liana_aggregate()

# Load necessary library
library(data.table)

# View the first few rows of the dataframe
head(liana_test)

# Perform FDR correction using the Benjamini-Hochberg method
liana_test$fdr_corrected_pvalue <- p.adjust(liana_test$cellphonedb.pvalue, method = "BH")

# View the updated dataframe
head(liana_test)

# Save the updated dataframe to a new CSV file (optional)
output_file_path1 <- "fdrcorrected_Significant_CCC_Malignant_AT1andMYE.csv"  # Adjust the path as needed
fwrite(liana_test, file = output_file_path1)

# Confirmation message
cat("FDR correction applied and saved to", output_file_path1, "\n")

corrected_liana_test1 <- read.csv("fdrcorrected_Significant_CCC_Malignant_AT1andMYE.csv")

dplyr::glimpse(corrected_liana_test1)
```

# run through the code below line by line!!!
```{r liana_ranks, warning=FALSE}
# We can aggregate these results into a tibble with consensus ranks
liana_test2 <- liana_test2 %>%
  liana_aggregate()

# Load necessary library
library(data.table)

# View the first few rows of the dataframe
head(liana_test2)

# Perform FDR correction using the Benjamini-Hochberg method
liana_test2$fdr_corrected_pvalue <- p.adjust(liana_test2$cellphonedb.pvalue, method = "BH")

# View the updated dataframe
head(liana_test2)

# Save the updated dataframe to a new CSV file (optional)
output_file_path2 <- "fdrcorrected_Significant_CCC_Malignant_AT2andMYE.csv"  # Adjust the path as needed
fwrite(liana_test2, file = output_file_path2)

# Confirmation message
cat("FDR correction applied and saved to", output_file_path2, "\n")


corrected_liana_test2 <- read.csv("fdrcorrected_Significant_CCC_Malignant_AT2andMYE.csv")

dplyr::glimpse(corrected_liana_test2)
```

Voila! That's it. A very brief intro to LIANA and how to obtain the scoring functions†
for each method implemented in it, as well as an aggregate_rank* which serves as a consensus across methods.

(†) Note that here we focus on the scores recommended to be used to prioritize interaction in a 
single sample system. Most of these, with the exception of SingleCellSignalR's LRscore, take
the specificity of the cluster pair into account.

(*) The aggregate consensus rank (`aggregate_rank`) is obtained using a re-implementation of the [`RRA`](https://www.ncbi.nlm.nih.gov/pmc/articles/PMC3278763/) method from the `RobustRankAggreg` package.

`RRA` scores can be interpreted as p-values and interactions which are
ranked consistently higher than random are assigned low scores/p-values.

## Simple DotPlot

We will now plot the results. By default, we use the `LRscore` from SingleCellSignalR
to represent the magnitude of expression of the ligand and receptor, and NATMI's `specificity weights`
to show how specific a given interaction is to the `source`(L) and `target`(R) cell types.

Note that the top 20 interactions (`ntop`), are defined as unique ligand-receptors 
ordered consequentially, regardless of the cell types. Done subsequent to filtering to the `source_groups` and `target_groups` of interest. In this case, we plot interactions in which B cells are the `source` (express the ~ligands) and the other 3 cell types are the `target` cell types (those which express the ~receptors).

By default, `liana_aggregate` would order LIANA's 
output according to `rank_aggregate` in ascending order. 

Note that using `liana_dotplot` /w `ntop` assumes that liana's results are 
ranked accordingly! Refer to `rank_method`, if you wish to easily rank
interactions for a single method (see example below).


# Assuming 'liana_res' is your data object
FIGURE 5b & 5c
# To visualize the CCC between cancer cells and myeloid cell population within Gramd2+ KRASG12D LUAD
# run through the code below line by line!!!
```{r}
# dotplot of CCC pattern from AT1 cells to myeloid cells in Gramd2+ KRASG12D LUAD
dotplot <- liana_dotplot(corrected_liana_test1,
                source_groups = c("AT1"),
                target_groups = c("iMONs", "TAMs", "Macrophages","DCs"),
                ntop = 20) 
dotplot$data$target <- factor(dotplot$data$target, levels=c("iMONs", "TAMs", "Macrophages","DCs"))
dotplot + scale_x_discrete(drop = FALSE)


# dotplot of CCC pattern from myeloid cells to AT1 cells in Gramd2+ KRASG12D LUAD
dotplot <- liana_dotplot(corrected_liana_test1,
                source_groups = c("iMONs", "TAMs", "Macrophages","DCs"),
                target_groups = c("AT1"),
                ntop = 20) 
dotplot$data$source <- factor(dotplot$data$source, levels=c("iMONs", "TAMs", "Macrophages","DCs"))
dotplot + scale_x_discrete(drop = FALSE)
```

FIGURE 5d & 5e
# To visualize the CCC between cancer cells and myeloid cell population within Sftpc+ KRASG12D LUAD
# run through the code below line by line!!!
```{r liana_dotplot, warning=FALSE, fig.width=11, fig.height=8}
# dotplot of CCC pattern from AT2 cells to myeloid cells in Sftpc+ KRASG12D LUAD
corrected_liana_test2 %>%
  liana_dotplot(source_groups = c("AT2"),
                target_groups = c("iMONs", "MDSCs", "TAMs", "Macrophages","DCs"),
                ntop = 20)

dotplot <- liana_dotplot(corrected_liana_test2,
                source_groups = c("AT2"),
                target_groups = c("iMONs", "MDSCs", "TAMs", "Macrophages","DCs"),
                ntop = 20) 
dotplot$data$target <- factor(dotplot$data$target, levels=c("iMONs", "MDSCs", "TAMs", "Macrophages","DCs"))
dotplot + scale_x_discrete(drop = FALSE)
```
# run through the code below line by line!!!
```{r liana_dotplot, warning=FALSE, fig.width=11, fig.height=8}
# dotplot of CCC pattern from myeloid cells to AT2 cells in Sftpc+ KRASG12D LUAD
corrected_liana_test2 %>%
  liana_dotplot(source_groups = c("iMONs", "MDSCs", "TAMs", "Macrophages","DCs"),
                target_groups = c("AT2"),
                ntop = 20)

dotplot <- liana_dotplot(corrected_liana_test2,
                source_groups = c("iMONs", "MDSCs", "TAMs", "Macrophages","DCs"),
                target_groups = c("AT2"),
                ntop = 20) 
dotplot$data$source <- factor(dotplot$data$source, levels=c("iMONs", "MDSCs", "TAMs", "Macrophages","DCs"))
dotplot + scale_x_discrete(drop = FALSE)
```
Note that missing dots are interactions which are not expressed in at least 10% of the cells (by default)
in both cell clusters.

In this case, we consider the specificity of interactions as defined by 
[NATMI's](https://www.nature.com/articles/s41467-020-18873-z) edge specificity weights. 
NATMI's specificity edges range from 0 to 1, where `1` means both the ligand
and receptor are uniquely expressed in a given pair of cells. 
Expression magnitude, represented by [SingleCellExperiment's LRScore](https://academic.oup.com/nar/article/48/10/e55/5810485), is on the other hand, 
meant to represent a non-negative regularized score, comparable between datasets.


## Frequency Heatmap
We will now plot the frequencies of interactions for each pair of *potentially* communicating cell types.

This heatmap was inspired by CellChat's and CellPhoneDB's heatmap designs.

First, we filter interactions by `aggregate_rank`, which can itself
 be treated as a p-value of for the robustly, highly ranked interactions.
Nevertheless, one can also filter according to [CPDB's p-value](https://www.nature.com/articles/s41596-020-0292-x) (<=0.05) or
[SingleCellExperiments LRScore](https://academic.oup.com/nar/article/48/10/e55/5810485), etc.

```{r heatfreq, warning=FALSE, fig.width=7, fig.height=5}
liana_trunc <- corrected_liana_test1 %>%
   # only keep interactions concordant between methods
  filter(aggregate_rank <= 0.01) # note that these pvals are already corrected

heat_freq(liana_trunc)
```

```{r heatfreq, warning=FALSE, fig.width=7, fig.height=5}
liana_trunc2 <- corrected_liana_test2 %>%
   # only keep interactions concordant between methods
  filter(aggregate_rank <= 0.01) # note that these pvals are already corrected

heat_freq(liana_trunc2)
```
Here, we see that `NK`-`CD8 T` share a relatively large number of the inferred
interactions, with many of those being send from `NK` - note large sum of 
interactions (gray barplot) in which NK is the `Sender`.

**NB!** Here, an assumption is implied that the number of interactions inferred between
cell types is informative of the communication events occurring in the system.
This is a rather strong assumption that one should consider prior to making any 
conclusions. [Our suggestion](https://www.biorxiv.org/content/10.1101/2021.05.21.445160v1.abstract) is that any conclusions should be complimented with further information, such as biological prior knowledge, spatial information, etc.

FIGURE 5a
Generate the correlation matrix plot by incorporating the two heatmaps generated above
# run through the code below line by line!!!
```{r}
# Load necessary libraries
library(ggplot2)
library(dplyr)
library(tidyr)


# Verify liana_trunc is correctly structured
str(liana_trunc)
liana_trunc <- as_tibble(liana_trunc)
str(liana_trunc)
# Create frequency tables using the correct column names
freq_table1 <- liana_trunc %>%
  count(source, target) %>%
  spread(key = target, value = n, fill = 0)

# Ensure the column names are correct and inspect a simple count operation
test_count <- liana_trunc %>%
    group_by(source, target) %>%
    tally()

# Print the result to check
print(test_count)

# Simplified spread operation to check the structure
freq_table1 <- test_count %>%
  spread(key = target, value = n, fill = 0)

# Print the resulting frequency table
print(freq_table1)

# Print the first few rows of liana_trunc
print(head(liana_trunc))

# Check the data types of the columns
print(sapply(liana_trunc, class))

# Attempt a simpler operation to ensure dplyr functions work
test_group <- liana_trunc %>%
  group_by(source, target) %>%
  summarize(count = n())

# Print the test_group result
print(test_group)


# Assume liana_trunc2 is the second dataset, ensure it is a tibble
liana_trunc2 <- as_tibble(liana_trunc2)


# Check the structure of liana_trunc2
str(liana_trunc2)

# Print the first few rows to inspect the data
print(head(liana_trunc2))
# Perform a simple count operation to check if dplyr functions work
test_count2 <- liana_trunc2 %>%
  group_by(source, target) %>%
  tally()

# Print the result to check
print(test_count2)

# Create frequency tables using the correct column names
freq_table2 <- test_count2 %>%
  spread(key = target, value = n, fill = 0)

# Print the resulting frequency table
print(freq_table2)

# Ensure both frequency tables have the same structure
all_senders <- union(freq_table1$source, freq_table2$source)
all_receivers <- union(names(freq_table1)[-1], names(freq_table2)[-1])


# Merge the data frames by source and target
merged_counts <- merge(test_count, test_count2, by = c("source", "target"), suffixes = c("_1", "_2"))

# Calculate the difference between the n columns
merged_counts <- merged_counts %>%
  mutate(diff = n_1 - n_2)

# Specify the order of levels
order_levels <- c("AT1", "AT1/AT2", "AT2", "HSPCs", "iMONs", "Proliferating Myeloid Cells", "MDSCs", "TAMs", "Macrophages", "DCs", "Granulocytes")

# Verify the levels in source and target
valid_levels_source <- intersect(order_levels, unique(merged_counts$source))
valid_levels_target <- intersect(order_levels, unique(merged_counts$target))

# Print the valid levels to verify
print(valid_levels_source)
print(valid_levels_target)

# Convert source and target to factors with the valid levels
merged_counts$source <- factor(merged_counts$source, levels = valid_levels_source)
merged_counts$target <- factor(merged_counts$target, levels = valid_levels_target)

# Spread the data for heatmap plotting with the specified order
diff_matrix_df <- merged_counts %>%
  select(source, target, diff) %>%
  spread(key = target, value = diff, fill = 0)

# Convert to matrix and set row names
diff_matrix <- as.matrix(diff_matrix_df[, -1])
rownames(diff_matrix) <- diff_matrix_df$source

# Ensure columns and rows are ordered correctly using valid levels
diff_matrix <- diff_matrix[valid_levels_source, valid_levels_target]

# Replace any remaining NA, NaN, or Inf values with 0
diff_matrix[is.na(diff_matrix)] <- 0
diff_matrix[is.nan(diff_matrix)] <- 0
diff_matrix[is.infinite(diff_matrix)] <- 0

# Inspect the final matrix to ensure order is correct and no invalid values
print(diff_matrix)

# Plot the heatmap with pheatmap, ensuring the specified order
pheatmap(diff_matrix, cluster_rows = FALSE, cluster_cols = FALSE,
         color = colorRampPalette(c("navy", "white", "firebrick3"))(100),
         xlab = "Receiver (Cell types)", ylab = "Sender (Cell types)",
         main = "Difference in Frequency of Interactions Between Cell Types")


pheatmap(diff_matrix,
         cluster_rows = FALSE,
         cluster_cols = FALSE,
         color = colorRampPalette(c("navy", "white", "firebrick3"))(100),
         border_color = NA,
         fontsize_row = 10,
         fontsize_col = 10,
         cellwidth = 20,
         cellheight = 20,
         main = "Difference in Frequency of Interactions Between Cell Types",
         display_numbers = F,
         number_color = "black",
         annotation_names_row = TRUE,
         annotation_names_col = TRUE,
         legend = TRUE,
         legend_labels = c("Low", "Medium", "High"),
         angle_col = "45")


```
````
