## Supplemental File 8 for "Cell of origin alters myeloid-mediated immunosuppression in lung adenocarcinoma": Supplemental file 9_TIME Visium Analysis.R

```
#### LOAD PACKAGES #########################
library(Seurat)
library(ggplot2)
library(patchwork)
library(dplyr)
library(hdf5r)
library(limma)
library(scales)
library(harmony)
library(readr)
library(estimate)
library(utils)
library(janitor)
library(ggpubr)
library(tidyr)

#### LOAD SFTPC-CreERT2 Control  ################
SftpcCre_data_dir <- '/Users/noahshulkin/Library/CloudStorage/OneDrive-CityofHopeNationalMedicalCenter/Marconett Lab/Visium Files/RUN2/Spc-Cre_Lung'
Sftpc_Cre<-Load10X_Spatial(SftpcCre_data_dir, filename = "filtered_feature_bc_matrix.h5", assay = "Spatial", slice = "slice1", filter.matrix = TRUE, to.upper = FALSE, image = NULL)
Sftpc_Cre$orig.ident <- "Sftpc_Cre"
Sftpc_Cre$Run <- "Run 2"
Sftpc_Cre <-AddMetaData(Sftpc_Cre, metadata = "Sftpc", col.name = "cell.of.origin")
Sftpc_Cre <-AddMetaData(Sftpc_Cre, metadata = "normal", col.name = "cancer")
Sftpc_Cre <- SCTransform(Sftpc_Cre, assay = "Spatial", verbose = FALSE)
Sftpc_Cre <- RunPCA(Sftpc_Cre, assay = "SCT", verbose = FALSE)
Sftpc_Cre <- FindNeighbors(Sftpc_Cre, reduction = "pca", dims = 1:30)
Sftpc_Cre <- FindClusters(Sftpc_Cre, verbose = FALSE)
Sftpc_Cre <- RunUMAP(Sftpc_Cre, reduction = "pca", dims = 1:30)
Sftpc_Cre <- RunTSNE(Sftpc_Cre, reduction = "pca", dims = 1:30)
#### LOAD GRAMD2-CreERT2 Control  ################
Gramd2_Cre_data_dir <- '/Users/noahshulkin/Library/CloudStorage/OneDrive-CityofHopeNationalMedicalCenter/Marconett Lab/Visium Files/RUN2/Gramd2-Cre_Lung'
Gramd2_Cre<-Load10X_Spatial(Gramd2_Cre_data_dir, filename = "filtered_feature_bc_matrix.h5", assay = "Spatial", slice = "slice1", filter.matrix = TRUE, to.upper = FALSE, image = NULL)
Gramd2_Cre$orig.ident <- "Gramd2_Cre"
Gramd2_Cre$Run <- "Run 2"
Gramd2_Cre <-AddMetaData(Gramd2_Cre, metadata = "Gramd2", col.name = "cell.of.origin")
Gramd2_Cre <-AddMetaData(Gramd2_Cre, metadata = "normal", col.name = "cancer")
Gramd2_Cre <- SCTransform(Gramd2_Cre, assay = "Spatial", verbose = FALSE)
Gramd2_Cre <- RunPCA(Gramd2_Cre, assay = "SCT", verbose = FALSE)
Gramd2_Cre <- FindNeighbors(Gramd2_Cre, reduction = "pca", dims = 1:30)
Gramd2_Cre <- FindClusters(Gramd2_Cre, verbose = FALSE)
Gramd2_Cre <- RunUMAP(Gramd2_Cre, reduction = "pca", dims = 1:30)
Gramd2_Cre <- RunTSNE(Gramd2_Cre, reduction = "pca", dims = 1:30)
#### LOAD SFTPC:KRAS-G12D Lung 1 ################
Sftpc1_data_dir <- '/Users/noahshulkin/Library/CloudStorage/OneDrive-CityofHopeNationalMedicalCenter/Marconett Lab/Visium Files/RUN1/Sftpc-KrasG12D_lung1-A'
Sftpc_lung1<-Load10X_Spatial(Sftpc1_data_dir, filename = "filtered_feature_bc_matrix.h5", assay = "Spatial", slice = "slice1", filter.matrix = TRUE, to.upper = FALSE, image = NULL)
Sftpc_lung1$orig.ident <- "Sftpc_1"
Sftpc_lung1$Run <- "Run 1"
Sftpc_lung1 <-AddMetaData(Sftpc_lung1, metadata = "Sftpc", col.name = "cell.of.origin")
Sftpc_lung1 <-AddMetaData(Sftpc_lung1, metadata = "LUAD", col.name = "cancer")
Sftpc_lung1 <- SCTransform(Sftpc_lung1, assay = "Spatial", verbose = FALSE)
Sftpc_lung1 <- RunPCA(Sftpc_lung1, assay = "SCT", verbose = FALSE)
Sftpc_lung1 <- FindNeighbors(Sftpc_lung1, reduction = "pca", dims = 1:30)
Sftpc_lung1 <- FindClusters(Sftpc_lung1, verbose = FALSE)
Sftpc_lung1 <- RunUMAP(Sftpc_lung1, reduction = "pca", dims = 1:30)
Sftpc_lung1 <- RunTSNE(Sftpc_lung1, reduction = "pca", dims = 1:30)
#### LOAD SFTPC:KRAS-G12D Lung 2 ################
Sftpc2_data_dir <- '/Users/noahshulkin/Library/CloudStorage/OneDrive-CityofHopeNationalMedicalCenter/Marconett Lab/Visium Files/RUN1/Sftpc-KrasG12D_lung1-D'
Sftpc_lung2<-Load10X_Spatial(Sftpc2_data_dir, filename = "filtered_feature_bc_matrix.h5", assay = "Spatial", slice = "slice1", filter.matrix = TRUE, to.upper = FALSE, image = NULL)
Sftpc_lung2$orig.ident <- "Sftpc_2"
Sftpc_lung2$Run <- "Run 1"
Sftpc_lung2 <-AddMetaData(Sftpc_lung2, metadata = "Sftpc", col.name = "cell.of.origin")
Sftpc_lung2 <-AddMetaData(Sftpc_lung2, metadata = "LUAD", col.name = "cancer")
Sftpc_lung2 <- SCTransform(Sftpc_lung2, assay = "Spatial", verbose = FALSE)
Sftpc_lung2 <- RunPCA(Sftpc_lung2, assay = "SCT", verbose = FALSE)
Sftpc_lung2 <- FindNeighbors(Sftpc_lung2, reduction = "pca", dims = 1:30)
Sftpc_lung2 <- FindClusters(Sftpc_lung2, verbose = FALSE)
Sftpc_lung2 <- RunUMAP(Sftpc_lung2, reduction = "pca", dims = 1:30)
Sftpc_lung2 <- RunTSNE(Sftpc_lung2, reduction = "pca", dims = 1:30)
#### LOAD GRAMD2:KRAS-G12D Lung 1 ###############
Gramd2_1_data_dir <- '/Users/noahshulkin/Library/CloudStorage/OneDrive-CityofHopeNationalMedicalCenter/Marconett Lab/Visium Files/RUN1/Gramd2-KrasG12D_lung1-B'
Gramd2_lung1<-Load10X_Spatial(Gramd2_1_data_dir, filename = "filtered_feature_bc_matrix.h5", assay = "Spatial", slice = "slice1", filter.matrix = TRUE, to.upper = FALSE, image = NULL)
Gramd2_lung1$orig.ident <- "Gramd2_1"
Gramd2_lung1$Run <- "Run 1"
Gramd2_lung1 <-AddMetaData(Gramd2_lung1, metadata = "Gramd2", col.name = "cell.of.origin")
Gramd2_lung1 <-AddMetaData(Gramd2_lung1, metadata = "LUAD", col.name = "cancer")
Gramd2_lung1 <- SCTransform(Gramd2_lung1, assay = "Spatial", verbose = FALSE)
Gramd2_lung1 <- RunPCA(Gramd2_lung1, assay = "SCT", verbose = FALSE)
Gramd2_lung1 <- FindNeighbors(Gramd2_lung1, reduction = "pca", dims = 1:30)
Gramd2_lung1 <- FindClusters(Gramd2_lung1, verbose = FALSE)
Gramd2_lung1 <- RunUMAP(Gramd2_lung1, reduction = "pca", dims = 1:30)
Gramd2_lung1 <- RunTSNE(Gramd2_lung1, reduction = "pca", dims = 1:30)
#### LOAD GRAMD2:KRAS-G12D Lung 2 ###############
Gramd2_2_data_dir <- '/Users/noahshulkin/Library/CloudStorage/OneDrive-CityofHopeNationalMedicalCenter/Marconett Lab/Visium Files/RUN1/Gramd2-KrasG12D_lung2-C'
Gramd2_lung2<-Load10X_Spatial(Gramd2_2_data_dir, filename = "filtered_feature_bc_matrix.h5", assay = "Spatial", slice = "slice1", filter.matrix = TRUE, to.upper = FALSE, image = NULL)
Gramd2_lung2$orig.ident <- "Gramd2_2"
Gramd2_lung2$Run <- "Run 1"
Gramd2_lung2 <-AddMetaData(Gramd2_lung2, metadata = "Gramd2", col.name = "cell.of.origin")
Gramd2_lung2 <-AddMetaData(Gramd2_lung2, metadata = "LUAD", col.name = "cancer")
Gramd2_lung2 <- SCTransform(Gramd2_lung2, assay = "Spatial", verbose = FALSE)
Gramd2_lung2 <- RunPCA(Gramd2_lung2, assay = "SCT", verbose = FALSE)
Gramd2_lung2 <- FindNeighbors(Gramd2_lung2, reduction = "pca", dims = 1:30)
Gramd2_lung2 <- FindClusters(Gramd2_lung2, verbose = FALSE)
Gramd2_lung2 <- RunUMAP(Gramd2_lung2, reduction = "pca", dims = 1:30)
Gramd2_lung2 <- RunTSNE(Gramd2_lung2, reduction = "pca", dims = 1:30)
#### LOAD GRAMD2:KRAS-G12D Lung 3 ################
Gramd2_3_data_dir <- '/Users/noahshulkin/Library/CloudStorage/OneDrive-CityofHopeNationalMedicalCenter/Marconett Lab/Visium Files/RUN2/GK024_Lung2'
Gramd2_lung3<-Load10X_Spatial(Gramd2_3_data_dir, filename = "filtered_feature_bc_matrix.h5", assay = "Spatial", slice = "slice1", filter.matrix = TRUE, to.upper = FALSE, image = NULL)
Gramd2_lung3$orig.ident <- "Gramd2_3"
Gramd2_lung3$Run <- "Run 2"
Gramd2_lung3 <-AddMetaData(Gramd2_lung3, metadata = "Gramd2", col.name = "cell.of.origin")
Gramd2_lung3 <-AddMetaData(Gramd2_lung3, metadata = "LUAD", col.name = "cancer")
Gramd2_lung3 <- SCTransform(Gramd2_lung3, assay = "Spatial", verbose = FALSE)
Gramd2_lung3 <- RunPCA(Gramd2_lung3, assay = "SCT", verbose = FALSE)
Gramd2_lung3 <- FindNeighbors(Gramd2_lung3, reduction = "pca", dims = 1:30)
Gramd2_lung3 <- FindClusters(Gramd2_lung3, verbose = FALSE)
Gramd2_lung3 <- RunUMAP(Gramd2_lung3, reduction = "pca", dims = 1:30)
Gramd2_lung3 <- RunTSNE(Gramd2_lung3, reduction = "pca", dims = 1:30)
#### LOAD SFTPC:KRAS-G12D Lung 3 ################
Sftpc_3_data_dir <- '/Users/noahshulkin/Library/CloudStorage/OneDrive-CityofHopeNationalMedicalCenter/Marconett Lab/Visium Files/RUN2/SK019_Lung2'
Sftpc_lung3<-Load10X_Spatial(Sftpc_3_data_dir, filename = "filtered_feature_bc_matrix.h5", assay = "Spatial", slice = "slice1", filter.matrix = TRUE, to.upper = FALSE, image = NULL)
Sftpc_lung3$orig.ident <- "Sftpc_3"
Sftpc_lung3$Run <- "Run 2"
Sftpc_lung3 <-AddMetaData(Sftpc_lung3, metadata = "Sftpc", col.name = "cell.of.origin")
Sftpc_lung3 <-AddMetaData(Sftpc_lung3, metadata = "LUAD", col.name = "cancer")
Sftpc_lung3 <- SCTransform(Sftpc_lung3, assay = "Spatial", verbose = FALSE)
Sftpc_lung3 <- RunPCA(Sftpc_lung3, assay = "SCT", verbose = FALSE)
Sftpc_lung3 <- FindNeighbors(Sftpc_lung3, reduction = "pca", dims = 1:30)
Sftpc_lung3 <- FindClusters(Sftpc_lung3, verbose = FALSE)
Sftpc_lung3 <- RunUMAP(Sftpc_lung3, reduction = "pca", dims = 1:30)
Sftpc_lung3 <- RunTSNE(Sftpc_lung3, reduction = "pca", dims = 1:30)

#### SAVE ALL INDIVIDUAL SETS POST-CLUSTERING ####
save(Sftpc_Cre, Gramd2_Cre, Sftpc_lung1, Sftpc_lung2, Gramd2_lung1, Gramd2_lung2, Gramd2_lung3, Sftpc_lung3, file = "/Users/noahshulkin/Library/CloudStorage/OneDrive-CityofHopeNationalMedicalCenter/Marconett Lab/Visium Files/postcluster.RData")
load("/Users/noahshulkin/Library/CloudStorage/OneDrive-CityofHopeNationalMedicalCenter/Marconett Lab/Visium Files/postcluster.RData")

#### MERGE ALL SPATIAL SETS ################
LUAD.merge <- merge(Sftpc_Cre, y = list(Sftpc_lung1, Sftpc_lung2, Sftpc_lung3, Gramd2_Cre, Gramd2_lung1, Gramd2_lung2, Gramd2_lung3), add.cell.ids = c('Sftpc_Cre', 'Sftpc_Kras_1', 'Sftpc_Kras_2', 'Sftpc_Kras_3', 'Gramd2_Cre', 'Gramd2_Kras_1', 'Gramd2_Kras_2', 'Gramd2_Kras_3'))
DefaultAssay(LUAD.merge) <- "SCT"
VariableFeatures(LUAD.merge) <- c(VariableFeatures(Sftpc_lung1), VariableFeatures(Sftpc_lung2), VariableFeatures(Sftpc_lung3), VariableFeatures(Sftpc_Cre), VariableFeatures(Gramd2_lung1), VariableFeatures(Gramd2_lung2), VariableFeatures(Gramd2_lung3), VariableFeatures(Gramd2_Cre))
LUAD.merge <- RunPCA(LUAD.merge, verbose = FALSE)
LUAD.merge <- FindNeighbors(LUAD.merge, dims = 1:30)
LUAD.merge <- FindClusters(LUAD.merge, res = 1, verbose = FALSE) 
LUAD.merge <- RunUMAP(LUAD.merge, dims = 1:30)

#### EXAMINE INITIAL UMAPS ####
DimPlot(LUAD.merge, reduction = "umap", label = T)
DimPlot(LUAD.merge, reduction = "umap", group.by = "Run")

#### RUN HARMONY INTEGRATION TO REDUCE BATCH EFFECTS ####
LUAD.merge <- LUAD.merge %>%
  RunHarmony(group.by.vars = 'Run', plot_covergence = FALSE)

LUAD.merge@reductions

LUAD.merge.embed <- Embeddings(LUAD.merge, "harmony")
LUAD.merge.embed[1:10,1:10]
#### CLUSTERING AFTER HARMONY INTEGRATION ####
LUAD.merge <- FindNeighbors(LUAD.merge, reduction = "harmony", dims = 1:30)
LUAD.merge <- FindClusters(LUAD.merge, reduction = "harmony", verbose = FALSE)
LUAD.merge <- RunUMAP(LUAD.merge, reduction = "harmony", dims = 1:30)
DimPlot(LUAD.merge, reduction = "umap", group.by = "Run")
DimPlot(LUAD.merge, reduction = "umap")

#### RENAMING CLUSTERS TO ICS ####
LUAD.merge$seurat_clusters <- as.factor(as.numeric(as.character(LUAD.merge$seurat_clusters)) + 1)
LUAD.merge$seurat_clusters <- paste("IC", LUAD.merge$seurat_clusters, sep = "")
Idents(LUAD.merge) <- "seurat_clusters"
levels(Idents(LUAD.merge)) <- c("IC1", "IC2", "IC3", "IC4", "IC5", "IC6", "IC7", "IC8", "IC9", "IC10", "IC11", "IC12", "IC13", "IC14", "IC15", "IC16", "IC17")
ReorderIdent(LUAD.merge, "seurat_clusters")
LUAD.merge$ICs <- Idents(LUAD.merge)
DimPlot(LUAD.merge, reduction = "umap", label = T)
head(Idents(LUAD.merge))

#### SAVING MERGED OBJECT FOLLOWING BATCH INTEGRATION ####
saveRDS(LUAD.merge, "/Users/noahshulkin/Library/CloudStorage/OneDrive-CityofHopeNationalMedicalCenter/Marconett Lab/Visium Files/postharmony.RDS")
LUAD.merge <- readRDS("/Users/noahshulkin/Library/CloudStorage/OneDrive-CityofHopeNationalMedicalCenter/Marconett Lab/Visium Files/postharmony.RDS")

#### EXAMINE QUALITY PER SPOT ##############
VlnPlot(LUAD.merge, features = "nCount_Spatial", pt.size = 0.1) + NoLegend()
VlnPlot(LUAD.merge, features = "nCount_Spatial", group.by = c('orig.ident'), pt.size = 0.1) + NoLegend()

SpatialFeaturePlot(LUAD.merge, features = "nCount_Spatial") + theme(legend.position = "right")
SpatialDimPlot(LUAD.merge, ncol = 4) + theme(legend.position = "right")
DimPlot(LUAD.merge, reduction = "umap", label = F, group.by = "cell.of.origin")
DimPlot(LUAD.merge, reduction = "umap", label = F, group.by = "cancer", cols = c("purple", "orange"))
DimPlot(LUAD.merge, reduction = "umap", label = F, group.by = "orig.ident")

#### GENE ENRICHMENT PER CLUSTER ####
LUAD.merge<-PrepSCTFindMarkers(LUAD.merge)
#Found 8 SCT models. Recorrecting SCT counts using minimum median counts: 2189
LUAD.markers <- FindAllMarkers(LUAD.merge, only.pos = TRUE, min.pct = 0.25, logfc.threshold = 0.25)
LUAD.markers <- read_csv("/Users/noahshulkin/Library/CloudStorage/OneDrive-CityofHopeNationalMedicalCenter/Marconett Lab/Visium Files/LUADmarkers.csv")


### Estimate for TME components ###########
## loaded package from download off R forge : estimate_1.0.13.tar.gz
library(utils)
rforge <- "http://r-forge.r-project.org"
install.packages("estimate", repos=rforge, dependencies=TRUE)
library(estimate)
library(help="estimate")

#### MODIFYING ESTIMATE TO WORK WITH MOUSE GENES ####
#GET ALL MOUSE AND HUMAN ORTHOLOGS, DOWNLOADED FROM MGI
setwd("/Users/noahshulkin/Documents/ESTIMATE")
gene.key <- read.delim("/Users/noahshulkin/Documents/ESTIMATE/genekey.txt")
#subset out mouse and human genes
gene.key.mouse <- which(gene.key$Common.Organism.Name == "mouse, laboratory")
gene.key.mouse <- gene.key[gene.key.mouse,] #gene.key.mouse will contain mouse genes
gene.key.human <- which(gene.key$Common.Organism.Name == "human")
gene.key.human <- gene.key[gene.key.human,] #gene.key.human will contain human genes

#MODIFYING COMMON GENES LIST IN ESTIMATE WITH MOUSE ORTHOLOGS
original.common.genes <- common_genes #replicate common genes data from estimate package
estimate.common.genes.human <- common_genes$GeneSymbol #extract human gene symbols
pos <- which(gene.key.human$Symbol %in% estimate.common.genes.human) #positions of genes to keep

common.gene.key <- gene.key.human[pos,1] #get DB.class.key corresponding to human genes

pos2 <- which(gene.key.mouse$DB.Class.Key %in% common.gene.key)
estimate.common.genes.mouse <- gene.key.mouse[pos2, 3] #get mouse symbols corresponding to DB.class.key list

dups <- which(duplicated(estimate.common.genes.mouse) == TRUE) #identify positions of duplicated genes
estimate.common.genes.mouse <- estimate.common.genes.mouse[-dups] #remove dups from mouse key

common_genes <- common_genes[1:10296,] #manipulate common_genes file to accomodate shorter gene list
common_genes$GeneSymbol <- estimate.common.genes.mouse #replace human gene symbols with mouse
common_genes$EntrezID <- "N/A"
common_genes$Synonyms <- "N/A"
common_genes$GeneName <- "N/A"
common_genes$Chromosome <- "N/A" #these columns just contain metadata and aren't required

#MODIFYING GENE SIGNATURES IN ESTIMATE WITH MOUSE ORTHOLOGS
original.genesets <- SI_geneset #replicate original gene sets
stromal.signature.human <- SI_geneset[1,2:142] #extract human stromal score
immune.signature.human <- SI_geneset[2,2:142] #extract human immune score

pos <- which(gene.key.human$Symbol %in% stromal.signature.human) #positions of genes to keep
stromal.gene.key <- gene.key.human[pos,1] #get corresponding DB.class.key
pos2 <- which(gene.key.mouse$DB.Class.Key %in% stromal.gene.key)
stromal.signature.mouse <- gene.key.mouse[pos2, 3] #get mouse symbols corresponding to determined keys

pos <- which(gene.key.human$Symbol %in% immune.signature.human) #positions of genes to keep
immune.gene.key <- gene.key.human[pos,1] #get corresponding DB.class.key
pos2 <- which(gene.key.mouse$DB.Class.Key %in% immune.gene.key)
immune.signature.mouse <- gene.key.mouse[pos2, 3] #get mouse symbols corresponding to determined keys

l <- list(1:143, 1:187) #put both signatures in a list
l[[1]] <- stromal.signature.mouse
l[[2]] <- immune.signature.mouse
names(l) <- c("StromalSignature", "ImmuneSignature")
df<- data.frame(lapply(l, `length<-`, max(lengths(l))))
SI_geneset <- t(df)

#### ESTIMATE on SFTPC-CreERT2     #########
setwd("/Users/noahshulkin/Library/CloudStorage/OneDrive-CityofHopeNationalMedicalCenter/Marconett Lab/Visium Files/RUN2/Spc-Cre_Lung")
ESTIMATE_Sftpc_Cre <- Read10X_h5(filename = "filtered_feature_bc_matrix.h5", use.names = TRUE, unique.features = TRUE)
class(ESTIMATE_Sftpc_Cre)
dim(ESTIMATE_Sftpc_Cre)
### write the matrix to excel csv file to format in excel:
write.csv(ESTIMATE_Sftpc_Cre, file = "ESTIMATE_Sftpc_Cre_matrixtoformat1.csv")
### need to manually change format (per Rania) by doing the following:
# re-save the .csv as a .txt (tab delimited text), then close file and manually change to .gct extension
# Add additional column 2 that's blank for "DESCRIPTION" and fill the cells in the column with "NA"
# Add row at the top, and in cell A1 write "#1.2"
# Add another, 2nd row with cell A2 the number of genes (rows) and cell B2 the number of spots (columns) in the SeuratObject
# use Data -> Table Tools -> remove duplicates ###
getwd()
filterCommonGenes(input.f = "ESTIMATE_Sftpc_Cre_matrixtoformat1.gct", output.f = "filtered.ESTIMATE.Sftpc_Cre-1.gct", id = "GeneSymbol")

estimateScore("filtered.ESTIMATE.Sftpc_Cre-1.gct", "ESTIMATE_Sftpc_Cre_estimate_score1.gct", platform="affymetrix")
## Open output file and format
# Remove top 2 lines so X... headers are at the top
# Remove "DESCRIPTION" column
# Remove erroneous First data column that's out of order and 1-base off
# save
estimate.Sftpc.Cre <- read.delim("/Users/noahshulkin/Documents/RUN2/Spc-Cre_Lung/ESTIMATE_Sftpc_Cre_estimate_score1.gct", row.names=1)
head(estimate.Sftpc.Cre[,1:5])
dim(estimate.Sftpc.Cre)

colnames(estimate.Sftpc.Cre) <- c(Cells(Sftpc_Cre))
dim(estimate.Sftpc.Cre)
head(estimate.Sftpc.Cre[,1:5])

#write.csv(estimate.Sftpc1, file="estimate.Sftpc1.csv")

estimate.assay.Sftpc.Cre <- CreateAssayObject(counts = estimate.Sftpc.Cre)
head(estimate.assay.Sftpc.Cre[1:4,1:4])
Sftpc_Cre[["ESTIMATE"]] <- estimate.assay.Sftpc.Cre
dim(Sftpc_Cre)
dim(estimate.assay.Sftpc.Cre)
DefaultAssay(Sftpc_Cre) <- "ESTIMATE"

save(Sftpc_Cre, file = "/Users/noahshulkin/Library/CloudStorage/OneDrive-CityofHopeNationalMedicalCenter/Marconett Lab/Visium Files/ESTIMATE.RData")

#### Plotting ESTIMATE results for Sftpc-Cre Control ####
Sftpc_Cre1 <- SpatialPlot(object = Sftpc_Cre,
                           features = "ImmuneScore",
                           slot = "counts",
                           crop = FALSE,
                           pt.size.factor = 1.2,
                           stroke = .1,
                           image.alpha = 0) +
  ggtitle(NULL) +
  theme(legend.direction = "vertical", legend.title = element_text(face = "bold"))
Sftpc_Cre1
Sftpc_Cre2 <- SpatialPlot(object = Sftpc_Cre,
                           features = "TumorPurity",
                           slot = "counts",
                           crop = FALSE,
                           pt.size.factor = 1.2,
                           stroke = .1,
                           image.alpha = 0) +
  ggtitle(NULL) +
  theme(legend.direction = "vertical", legend.title = element_text(face = "bold"))
Sftpc_Cre2
Sftpc_Cre3 <- SpatialPlot(object = Sftpc_Cre,
                           features = "StromalScore",
                           slot = "counts",
                           crop = FALSE,
                           pt.size.factor = 1.2,
                           stroke = .1,
                           image.alpha = 0) +
  ggtitle(NULL) +
  theme(legend.direction = "vertical", legend.title = element_text(face = "bold"))
Sftpc_Cre3
Sftpc_Cre1 + Sftpc_Cre2 + Sftpc_Cre3


#### ESTIMATE on Gramd2-CreERT2 ####
setwd("/Users/noahshulkin/Library/CloudStorage/OneDrive-CityofHopeNationalMedicalCenter/Marconett Lab/Visium Files/RUN2/Gramd2-Cre_Lung")
ESTIMATE_Gramd2_Cre <- Read10X_h5(filename = "filtered_feature_bc_matrix.h5", use.names = TRUE, unique.features = TRUE)
class(ESTIMATE_Gramd2_Cre)
dim(ESTIMATE_Gramd2_Cre)
### write the matrix to excel csv file to format in excel:
write.csv(ESTIMATE_Gramd2_Cre, file = "ESTIMATE_Gramd2_Cre_matrixtoformat1.csv")
### need to manually change format (per Rania) by doing the following:
# re-save the .csv as a .txt (tab delimited text), then close file and manually change to .gct extension
# Add additional column 2 that's blank for "DESCRIPTION" and fill the cells in the column with "NA"
# Add row at the top, and in cell A1 write "#1.2"
# Add another, 2nd row with cell A2 the number of genes (rows) and cell B2 the number of spots (columns) in the SeuratObject
# use Data -> Table Tools -> remove duplicates ###
getwd()
filterCommonGenes(input.f = "ESTIMATE_Gramd2_Cre_matrixtoformat1.gct", output.f = "filtered.ESTIMATE.Gramd2_Cre-1.gct", id = "GeneSymbol")

estimateScore("filtered.ESTIMATE.Gramd2_Cre-1.gct", "ESTIMATE_Gramd2_Cre_estimate_score1.gct", platform="affymetrix")
## Open output file and format
# Remove top 2 lines so X... headers are at the top
# Remove "DESCRIPTION" column
# Remove erroneous First data column that's out of order and 1-base off
# save
estimate.Gramd2.Cre <- read.delim("/Users/noahshulkin/Library/CloudStorage/OneDrive-CityofHopeNationalMedicalCenter/Marconett Lab/Visium Files/RUN2/Gramd2-Cre_Lung/ESTIMATE_Gramd2_Cre_estimate_score1.gct", row.names=1)
head(estimate.Gramd2.Cre[,1:5])
dim(estimate.Gramd2.Cre)

colnames(estimate.Gramd2.Cre) <- c(Cells(Gramd2_Cre))
dim(estimate.Gramd2.Cre)
head(estimate.Gramd2.Cre[,1:5])

#write.csv(estimate.Gramd2.Cre, file="estimate.Gramd2.Cre.csv")

estimate.assay.Gramd2.Cre <- CreateAssayObject(counts = estimate.Gramd2.Cre)
head(estimate.assay.Gramd2.Cre[1:4,1:4])
Gramd2_Cre[["ESTIMATE"]] <- estimate.assay.Gramd2.Cre
dim(Gramd2_Cre)
dim(estimate.assay.Gramd2.Cre)
DefaultAssay(Gramd2_Cre) <- "ESTIMATE"
#### Plotting ESTIMATE results for Gramd2-Cre Control ####
Gramd2_Cre1 <- SpatialPlot(object = Gramd2_Cre,
                           features = "ImmuneScore",
                           slot = "counts",
                           crop = FALSE,
                           pt.size.factor = 1.2,
                           stroke = .1,
                           image.alpha = 0) +
  ggtitle(NULL) +
  theme(legend.direction = "vertical", legend.title = element_text(face = "bold"))
Gramd2_Cre1
Gramd2_Cre2 <- SpatialPlot(object = Gramd2_Cre,
                           features = "TumorPurity",
                           slot = "counts",
                           crop = FALSE,
                           pt.size.factor = 1.2,
                           stroke = .1,
                           image.alpha = 0) +
  ggtitle(NULL) +
  theme(legend.direction = "vertical", legend.title = element_text(face = "bold"))
Gramd2_Cre2
Gramd2_Cre3 <- SpatialPlot(object = Gramd2_Cre,
                           features = "StromalScore",
                           slot = "counts",
                           crop = FALSE,
                           pt.size.factor = 1.2,
                           stroke = .1,
                           image.alpha = 0) +
  ggtitle(NULL) +
  theme(legend.direction = "vertical", legend.title = element_text(face = "bold"))
Gramd2_Cre3
Gramd2_Cre1 + Gramd2_Cre2 + Gramd2_Cre3


#### ESTIMATE on SFTPC:KRAS_LUNG_1 #########
setwd("/Users/noahshulkin/Library/CloudStorage/OneDrive-CityofHopeNationalMedicalCenter/Marconett Lab/Visium Files/RUN1/Sftpc-KrasG12D_lung1-A")
ESTIMATE_Sftpc_lung1 <- Read10X_h5(filename = "filtered_feature_bc_matrix.h5", use.names = TRUE, unique.features = TRUE)
class(ESTIMATE_Sftpc_lung1)
dim(ESTIMATE_Sftpc_lung1)
### write the matrix to excel csv file to format in excel:
write.csv(ESTIMATE_Sftpc_lung1, file = "ESTIMATE_Sftpc_Lung1_matrixtoformat1.csv")
### need to manually change format (per Rania) by doing the following:
# switch .csv to .gct extension
# Add additional column 2 that's blank for "DESCRIPTION" 
# Add row at the top, and in cell A1 write "#1.2"
# Add another, 2nd row with cell A2 the number of genes (rows) and cell B2 the number of spots (columns) in the SeuratObject
# use Data -> Table Tools to remove duplicates ###
getwd()
filterCommonGenes(input.f = "ESTIMATE_Sftpc_Lung1_matrixtoformat1.gct", output.f = "filtered.ESTIMATE.Sftpc_lung1-1.gct", id = "GeneSymbol")

estimateScore("filtered.ESTIMATE.Sftpc_lung1-1.gct", "ESTIMATE_Sftpc_lung1_estimate_score1.gct", platform="affymetrix")
## Open output file and format
# Remove top 2 lines so X... headers are at the top
# Remove "DESCRIPTION" column
# Remove erroneous First data column that's out of order and 1-base off
# save
estimate.Sftpc1 <- read.delim("/Users/noahshulkin/Library/CloudStorage/OneDrive-CityofHopeNationalMedicalCenter/Marconett Lab/Visium Files/RUN1/Sftpc-KrasG12D_lung1-A/ESTIMATE_Sftpc_lung1_estimate_score1.gct", row.names=1)
head(estimate.Sftpc1[,1:5])
dim(estimate.Sftpc1)

colnames(estimate.Sftpc1) <- c(Cells(Sftpc_lung1))
dim(estimate.Sftpc1)
head(estimate.Sftpc1[,1:5])

#write.csv(estimate.Sftpc1, file="estimate.Sftpc1.csv")

estimate.assay.Sftpc_Lung1 <- CreateAssayObject(counts = estimate.Sftpc1)
head(estimate.assay.Sftpc_Lung1[1:4,1:4])
Sftpc_lung1[["ESTIMATE"]] <- estimate.assay.Sftpc_Lung1
dim(Sftpc_lung1)
dim(estimate.assay.Sftpc_Lung1)
DefaultAssay(Sftpc_lung1) <- "ESTIMATE"

#### Plotting ESTIMATE results for Sftpc:Kras Lung 1 ####
A1 <- SpatialPlot(object = Sftpc_lung1,
                  features = "ImmuneScore",
                  slot = "counts",
                  crop = FALSE,
                  pt.size.factor = 1.2,
                  stroke = .1,
                  image.alpha = 0) +
  ggtitle(NULL) +
  theme(legend.direction = "vertical", legend.title = element_text(face = "bold"))
A1
A2 <- SpatialPlot(object = Sftpc_lung1,
                  features = "TumorPurity",
                  slot = "counts",
                  crop = FALSE,
                  pt.size.factor = 1.2,
                  stroke = .1,
                  image.alpha = 0) +
  ggtitle(NULL) +
  theme(legend.direction = "vertical", legend.title = element_text(face = "bold"))
A2
A3 <- SpatialPlot(object = Sftpc_lung1,
                  features = "StromalScore",
                  slot = "counts",
                  crop = FALSE,
                  pt.size.factor = 1.2,
                  stroke = .1,
                  image.alpha = 0) +
  ggtitle(NULL) +
  theme(legend.direction = "vertical", legend.title = element_text(face = "bold"))
A1 + A2 + A3
A3


#### ESTIMATE on SFTPC:KRAS_LUNG_2 #########
setwd("/Users/noahshulkin/Library/CloudStorage/OneDrive-CityofHopeNationalMedicalCenter/Marconett Lab/Visium Files/RUN1/Sftpc-KrasG12D_lung1-D")
ESTIMATE_Sftpc_lung2 <- Read10X_h5(filename = "filtered_feature_bc_matrix.h5", use.names = TRUE, unique.features = TRUE)
class(ESTIMATE_Sftpc_lung2)
dim(ESTIMATE_Sftpc_lung2)
#19465 (rows/genes/features) #1588 (columns/barcodes/spots)

### write the matrix to excel csv file to format in excel:
write.csv(ESTIMATE_Sftpc_lung2, file = "ESTIMATE_Sftpc_Lung2_matrixtoformat1.csv")
### need to manually change format (per Rania) by doing the following:
# switch .csv to .gct extension
# Add additional column 2 that's blank for "DESCRIPTION" 
# Add row at the top, and in cell A1 write "#1.2"
# Add another, 2nd row with cell A2 the number of genes (rows) and cell B2 the number of spots (columns) in the SeuratObject
# use Data -> Table Tools to remove duplicates ###
getwd()
filterCommonGenes(input.f = "ESTIMATE_Sftpc_Lung2_matrixtoformat1.gct", output.f = "filtered.ESTIMATE.Sftpc_lung2-1.gct", id = "GeneSymbol")

estimateScore("filtered.ESTIMATE.Sftpc_lung2-1.gct", "ESTIMATE_Sftpc_lung2_estimate_score1.gct", platform="affymetrix")
## Open output file and format
# Remove top 2 lines so X... headers are at the top
# Remove "DESCRIPTION" column
# Remove erroneous First data column that's out of order and 1-base off
# save

estimate.Sftpc2 <- read.delim("/Users/noahshulkin/Library/CloudStorage/OneDrive-CityofHopeNationalMedicalCenter/Marconett Lab/Visium Files/RUN1/Sftpc-KrasG12D_lung1-D/ESTIMATE_Sftpc_lung2_estimate_score1.gct", row.names=1)
head(estimate.Sftpc2[,1:5])
dim(estimate.Sftpc2)

colnames(estimate.Sftpc2) <- c(Cells(Sftpc_lung2))
dim(estimate.Sftpc2)
head(estimate.Sftpc2[,1:5])

#write.csv(estimate.Sftpc2, file="estimate.Sftpc2.csv")

estimate.assay.Sftpc_Lung2 <- CreateAssayObject(counts = estimate.Sftpc2)
head(estimate.assay.Sftpc_Lung2[1:4,1:4])
Sftpc_lung2[["ESTIMATE"]] <- estimate.assay.Sftpc_Lung2
dim(Sftpc_lung2)
dim(estimate.assay.Sftpc_Lung2)
DefaultAssay(Sftpc_lung2) <- "ESTIMATE"
#### Plotting ESTIMATE results for Sftpc:Kras Lung 2 ####
B1 <- SpatialPlot(object = Sftpc_lung2,
                  features = "ImmuneScore",
                  slot = "counts",
                  crop = FALSE,
                  pt.size.factor = 1.2,
                  stroke = .1,
                  image.alpha = 0) +
  ggtitle(NULL) +
  theme(legend.direction = "vertical", legend.title = element_text(face = "bold"))
B1
B2 <- SpatialPlot(object = Sftpc_lung2,
                  features = "TumorPurity",
                  slot = "counts",
                  crop = FALSE,
                  pt.size.factor = 1.2,
                  stroke = .1,
                  image.alpha = 0) +
  ggtitle(NULL) +
  theme(legend.direction = "vertical", legend.title = element_text(face = "bold"))
B2
B3 <- SpatialPlot(object = Sftpc_lung2,
                  features = "StromalScore",
                  slot = "counts",
                  crop = FALSE,
                  pt.size.factor = 1.2,
                  stroke = .1,
                  image.alpha = 0) +
  ggtitle(NULL) +
  theme(legend.direction = "vertical", legend.title = element_text(face = "bold"))
B1 + B2 + B3
B3


#### ESTIMATE on SFTPC:KRAS_LUNG_3 #########
setwd("/Users/noahshulkin/Library/CloudStorage/OneDrive-CityofHopeNationalMedicalCenter/Marconett Lab/Visium Files/RUN2/SK019_Lung2")
ESTIMATE_Sftpc_Lung3 <- Read10X_h5(filename = "filtered_feature_bc_matrix.h5", use.names = TRUE, unique.features = TRUE)
class(ESTIMATE_Sftpc_Lung3)
dim(ESTIMATE_Sftpc_Lung3)
### write the matrix to excel csv file to format in excel:
write.csv(ESTIMATE_Sftpc_Lung3, file = "ESTIMATE_Sftpc_Lung3_matrixtoformat1.csv")
### need to manually change format (per Rania) by doing the following:
# re-save the .csv as a .txt (tab delimited text), then close file and manually change to .gct extension
# Add additional column 2 that's blank for "DESCRIPTION" and fill the cells in the column with "NA"
# Add row at the top, and in cell A1 write "#1.2"
# Add another, 2nd row with cell A2 the number of genes (rows) and cell B2 the number of spots (columns) in the SeuratObject
# use Data -> Table Tools -> remove duplicates (rows/lines 1066 and 1067 are Mar-1, Mar-2, correct to gene names with "text" setting) ###
getwd()
filterCommonGenes(input.f = "ESTIMATE_Sftpc_Lung3_matrixtoformat1.gct", output.f = "filtered.ESTIMATE.Sftpc_Lung3-1.gct", id = "GeneSymbol")

estimateScore("filtered.ESTIMATE.Sftpc_Lung3-1.gct", "ESTIMATE_Sftpc_Lung3_estimate_score1.gct", platform="affymetrix")
## Open output file and format
# Remove top 2 lines so X... headers are at the top
# Remove "DESCRIPTION" column
# Remove erroneous First data column that's out of order and 1-base off
# save
estimate.Sftpc.Lung3 <- read.delim("/Users/noahshulkin/Library/CloudStorage/OneDrive-CityofHopeNationalMedicalCenter/Marconett Lab/Visium Files/RUN2/SK019_Lung2/ESTIMATE_Sftpc_Lung3_estimate_score1.gct", row.names=1)
head(estimate.Sftpc.Lung3[,1:5])
dim(estimate.Sftpc.Lung3)

colnames(estimate.Sftpc.Lung3) <- c(Cells(Sftpc_lung3))
dim(estimate.Sftpc.Lung3)
head(estimate.Sftpc.Lung3[,1:5])

#write.csv(estimate.Sftpc.Lung3, file="estimate.Sftpc_Lung3.csv")

estimate.assay.Sftpc.Lung3 <- CreateAssayObject(counts = estimate.Sftpc.Lung3)
head(estimate.assay.Sftpc.Lung3[1:4,1:4])
Sftpc_lung3[["ESTIMATE"]] <- estimate.assay.Sftpc.Lung3
dim(Sftpc_lung3)
dim(estimate.assay.Sftpc.Lung3)
DefaultAssay(Sftpc_lung3) <- "ESTIMATE"


#### Plotting ESTIMATE results for Sftpc:Kras Lung 2 ####
E1 <- SpatialPlot(object = Sftpc_lung3,
                  features = "ImmuneScore",
                  slot = "counts",
                  crop = FALSE,
                  pt.size.factor = 1.2,
                  stroke = .1,
                  image.alpha = 0) +
  ggtitle(NULL) +
  theme(legend.direction = "vertical", legend.title = element_text(face = "bold"))
E1
E2 <- SpatialPlot(object = Sftpc_lung3,
                  features = "TumorPurity",
                  slot = "counts",
                  crop = FALSE,
                  pt.size.factor = 1.2,
                  stroke = .1,
                  image.alpha = 0) +
  ggtitle(NULL) +
  theme(legend.direction = "vertical", legend.title = element_text(face = "bold"))
E2
E3 <- SpatialPlot(object = Sftpc_lung3,
                  features = "StromalScore",
                  slot = "counts",
                  crop = FALSE,
                  pt.size.factor = 1.2,
                  stroke = .1,
                  image.alpha = 0) +
  ggtitle(NULL) +
  theme(legend.direction = "vertical", legend.title = element_text(face = "bold"))
E3
E1 + E2 + E3


#### ESTIMATE on GRAMD2:KRAS_LUNG_1 ########
setwd("/Users/noahshulkin/Library/CloudStorage/OneDrive-CityofHopeNationalMedicalCenter/Marconett Lab/Visium Files/RUN1/Gramd2-KrasG12D_lung1-B")
ESTIMATE_Gramd2_lung1 <- Read10X_h5(filename = "filtered_feature_bc_matrix.h5", use.names = TRUE, unique.features = TRUE)
class(ESTIMATE_Gramd2_lung1)
dim(ESTIMATE_Gramd2_lung1)

### write the matrix to excel csv file to format in excel:
write.csv(ESTIMATE_Gramd2_lung1, file = "ESTIMATE_Gramd2_Lung1_matrixtoformat1.csv")
### need to manually change format to .gct by changing columns and rows in excel and then chancing extension to .gct from .txt per Rania, also use Data -> Table Tools to remove duplicates ###

getwd()
filterCommonGenes(input.f = "ESTIMATE_Gramd2_Lung1_matrixtoformat1.gct", output.f = "filtered.ESTIMATE.Gramd2_lung1-1.gct", id = "GeneSymbol")

estimateScore("filtered.ESTIMATE.Gramd2_lung1-1.gct", "ESTIMATE_Gramd2_lung1_estimate_score1.gct", platform="affymetrix")
### remove header rows/columns in excel as above
estimate.Gramd2Lung1 <- read.delim("/Users/noahshulkin/Library/CloudStorage/OneDrive-CityofHopeNationalMedicalCenter/Marconett Lab/Visium Files/RUN1/Gramd2-KrasG12D_lung1-B/ESTIMATE_Gramd2_lung1_estimate_score1.gct", row.names=1)
head(estimate.Gramd2Lung1[,1:5])
dim(estimate.Gramd2Lung1)
#dim(ESTIMATE_Gramd2_lung1)

colnames(estimate.Gramd2Lung1) <- c(Cells(Gramd2_lung1))
dim(estimate.Gramd2Lung1)
head(estimate.Gramd2Lung1[,1:5])

#write.csv(estimate.Gramd2Lung1, file="estimate.Gramd2Lung1.csv")

estimate.assay.Gramd2_Lung1 <- CreateAssayObject(counts = estimate.Gramd2Lung1)
head(estimate.assay.Gramd2_Lung1[1:4,1:4])
Gramd2_lung1[["ESTIMATE"]] <- estimate.assay.Gramd2_Lung1
dim(Gramd2_lung1)
dim(estimate.assay.Gramd2_Lung1)
DefaultAssay(Gramd2_lung1) <- "ESTIMATE"
#### Plotting ESTIMATE results for Gramd2:Kras Lung 1 ####
C1 <- SpatialPlot(object = Gramd2_lung1,
                  features = "ImmuneScore",
                  slot = "counts",
                  crop = FALSE,
                  pt.size.factor = 1.2,
                  stroke = .1,
                  image.alpha = 0) +
  ggtitle(NULL) +
  theme(legend.direction = "vertical", legend.title = element_text(face = "bold"))
C1
C2 <- SpatialPlot(object = Gramd2_lung1,
                  features = "TumorPurity",
                  slot = "counts",
                  crop = FALSE,
                  pt.size.factor = 1.2,
                  stroke = .1,
                  image.alpha = 0) +
  ggtitle(NULL) +
  theme(legend.direction = "vertical", legend.title = element_text(face = "bold"))
C2
C3 <- SpatialPlot(object = Gramd2_lung1,
                  features = "StromalScore",
                  slot = "counts",
                  crop = FALSE,
                  pt.size.factor = 1.2,
                  stroke = .1,
                  image.alpha = 0) +
  ggtitle(NULL) +
  theme(legend.direction = "vertical", legend.title = element_text(face = "bold"))
C1 + C2 + C3
C3


#### ESTIMATE on GRAMD2:KRAS_LUNG_2 ########
setwd("/Users/noahshulkin/Library/CloudStorage/OneDrive-CityofHopeNationalMedicalCenter/Marconett Lab/Visium Files/RUN1/Gramd2-KrasG12D_lung2-C")
ESTIMATE_Gramd2_lung2 <- Read10X_h5(filename = "filtered_feature_bc_matrix.h5", use.names = TRUE, unique.features = TRUE)
class(ESTIMATE_Gramd2_lung2)
dim(ESTIMATE_Gramd2_lung2)
#19465 (rows/genes/features) #2083 (columns/barcodes/spots)

### write the matrix to excel csv file to format in excel:
write.csv(ESTIMATE_Gramd2_lung2, file = "ESTIMATE_Gramd2_Lung2_matrixtoformat1.csv")
### need to manually change format to .gct by changing columns and rows in excel and then chancing extension to .gct from .txt per Rania, also use Data -> Table Tools to remove duplicates ###

getwd()
filterCommonGenes(input.f = "ESTIMATE_Gramd2_Lung2_matrixtoformat1.gct", output.f = "filtered.ESTIMATE.Gramd2_lung2-1.gct", id = "GeneSymbol")

estimateScore("filtered.ESTIMATE.Gramd2_lung2-1.gct", "ESTIMATE_Gramd2_lung2_estimate_score1.gct", platform="affymetrix")
### remove crap rows (top 2)/columns (second and third) in excel, then import again
estimate.Gramd2Lung2 <- read.delim("/Users/noahshulkin/Library/CloudStorage/OneDrive-CityofHopeNationalMedicalCenter/Marconett Lab/Visium Files/RUN1/Gramd2-KrasG12D_lung2-C/ESTIMATE_Gramd2_lung2_estimate_score1.gct", row.names=1)
head(estimate.Gramd2Lung2[,1:5])
dim(estimate.Gramd2Lung2)
#dim(ESTIMATE_Gramd2_lung2)

colnames(estimate.Gramd2Lung2) <- c(Cells(Gramd2_lung2))
dim(estimate.Gramd2Lung2)
head(estimate.Gramd2Lung2[,1:5])

#write.csv(estimate.Gramd2Lung2, file="estimate.Gramd2Lung2.csv")

estimate.assay.Gramd2_Lung2 <- CreateAssayObject(counts = estimate.Gramd2Lung2)
head(estimate.assay.Gramd2_Lung2[1:4,1:4])
Gramd2_lung2[["ESTIMATE"]] <- estimate.assay.Gramd2_Lung2
dim(Gramd2_lung2)
dim(estimate.assay.Gramd2_Lung2)
DefaultAssay(Gramd2_lung2) <- "ESTIMATE"
#### Plotting ESTIMATE results for Gramd2:Kras Lung 2 ####
D1 <- SpatialPlot(object = Gramd2_lung2,
                  features = "ImmuneScore",
                  slot = "counts",
                  crop = FALSE,
                  pt.size.factor = 1.2,
                  stroke = .1,
                  image.alpha = 0) +
  ggtitle(NULL) +
  theme(legend.direction = "vertical", legend.title = element_text(face = "bold"))
D1
D2 <- SpatialPlot(object = Gramd2_lung2,
                  features = "TumorPurity",
                  slot = "counts",
                  crop = FALSE,
                  pt.size.factor = 1.2,
                  stroke = .1,
                  image.alpha = 0) +
  ggtitle(NULL) +
  theme(legend.direction = "vertical", legend.title = element_text(face = "bold"))
D2
D3 <- SpatialPlot(object = Gramd2_lung2,
                  features = "StromalScore",
                  slot = "counts",
                  crop = FALSE,
                  pt.size.factor = 1.2,
                  stroke = .1,
                  image.alpha = 0) +
  ggtitle(NULL) +
  theme(legend.direction = "vertical", legend.title = element_text(face = "bold"))
D3
D1 + D2 + D3


#### ESTIMATE on GRAMD2:KRAS_LUNG_3 ########
setwd("/Users/noahshulkin/Library/CloudStorage/OneDrive-CityofHopeNationalMedicalCenter/Marconett Lab/Visium Files/RUN2/GK024_Lung2")
ESTIMATE_Gramd2_lung3 <- Read10X_h5(filename = "filtered_feature_bc_matrix.h5", use.names = TRUE, unique.features = TRUE)
class(ESTIMATE_Gramd2_lung3)
dim(ESTIMATE_Gramd2_lung3)
### write the matrix to excel csv file to format in excel:
write.csv(ESTIMATE_Gramd2_lung3, file = "ESTIMATE_Gramd2_lung3_matrixtoformat1.csv")
### need to manually change format (per Rania) by doing the following:
# re-save the .csv as a .txt (tab delimited text), then close file and manually change to .gct extension
# Add additional column 2 that's blank for "DESCRIPTION" and fill the cells in the column with "NA"
# Add row at the top, and in cell A1 write "#1.2"
# Add another, 2nd row with cell A2 the number of genes (rows) and cell B2 the number of spots (columns) in the SeuratObject
# use Data -> Table Tools -> remove duplicates ###
getwd()
filterCommonGenes(input.f = "ESTIMATE_Gramd2_lung3_matrixtoformat1.gct", output.f = "filtered.ESTIMATE.Gramd2_lung3-1.gct", id = "GeneSymbol")

estimateScore("filtered.ESTIMATE.Gramd2_lung3-1.gct", "ESTIMATE_Gramd2_lung3_estimate_score1.gct", platform="affymetrix")
## Open output file and format
# Remove top 2 lines so X... headers are at the top
# Remove "DESCRIPTION" column
# Remove erroneous First data column that's out of order and 1-base off
# save
estimate.Gramd2.lung3 <- read.delim("/Users/noahshulkin/Library/CloudStorage/OneDrive-CityofHopeNationalMedicalCenter/Marconett Lab/Visium Files/RUN2/GK024_Lung2/ESTIMATE_Gramd2_lung3_estimate_score1.gct", row.names=1)
head(estimate.Gramd2.lung3[,1:5])
dim(estimate.Gramd2.lung3)

colnames(estimate.Gramd2.lung3) <- c(Cells(Gramd2_lung3))
dim(estimate.Gramd2.lung3)
head(estimate.Gramd2.lung3[,1:5])

#write.csv(estimate.Gramd2.lung3, file="estimate.Gramd2.lung3.csv")

estimate.assay.Gramd2.lung3 <- CreateAssayObject(counts = estimate.Gramd2.lung3)
head(estimate.assay.Gramd2.lung3[1:4,1:4])
Gramd2_lung3[["ESTIMATE"]] <- estimate.assay.Gramd2.lung3
dim(Gramd2_lung3)
dim(estimate.assay.Gramd2.lung3)
DefaultAssay(Gramd2_lung3) <- "ESTIMATE"


#### Plotting ESTIMATE results for Gramd2:Kras Lung 3 ####
E1 <- SpatialPlot(object = Gramd2_lung3,
                  features = "ImmuneScore",
                  slot = "counts",
                  crop = FALSE,
                  pt.size.factor = 1.2,
                  stroke = .1,
                  image.alpha = 0) +
  ggtitle(NULL) +
  theme(legend.direction = "vertical", legend.title = element_text(face = "bold"))
E1
E2 <- SpatialPlot(object = Gramd2_lung3,
                  features = "TumorPurity",
                  slot = "counts",
                  crop = FALSE,
                  pt.size.factor = 1.2,
                  stroke = .1,
                  image.alpha = 0) +
  ggtitle(NULL) +
  theme(legend.direction = "vertical", legend.title = element_text(face = "bold"))
E2
E3 <- SpatialPlot(object = Gramd2_lung3,
                  features = "StromalScore",
                  slot = "counts",
                  crop = FALSE,
                  pt.size.factor = 1.2,
                  stroke = .1,
                  image.alpha = 0) +
  ggtitle(NULL) +
  theme(legend.direction = "vertical", legend.title = element_text(face = "bold"))
E3
E1 + E2 + E3


#### SAVING/LOADING INDIVIDUAL SEURAT OBJECTS POST-ESTIMATE ####
save(Sftpc_Cre, Sftpc_lung1, Gramd2_Cre, Sftpc_lung2, Sftpc_lung3, Gramd2_lung1, Gramd2_lung2, Gramd2_lung3, file = "/Users/noahshulkin/Library/CloudStorage/OneDrive-CityofHopeNationalMedicalCenter/Marconett Lab/Visium Files/ESTIMATE.RData")
load("/Users/noahshulkin/Library/CloudStorage/OneDrive-CityofHopeNationalMedicalCenter/Marconett Lab/Visium Files/ESTIMATE.RData")


### RE-GENERATE MERGED SEURAT WITH ESTIMATE ANNOTATION ####
ESTIMATE.merge <- merge(Sftpc_Cre, y = list(Sftpc_lung1, Sftpc_lung2, Sftpc_lung3, Gramd2_Cre, Gramd2_lung1, Gramd2_lung2, Gramd2_lung3), add.cell.ids = c('Sftpc_Cre', 'Sftpc_Kras_1', 'Sftpc_Kras_2', 'Sftpc_Kras_3', 'Gramd2_Cre', 'Gramd2_Kras_1', 'Gramd2_Kras_2', 'Gramd2_Kras_3'))
DefaultAssay(ESTIMATE.merge) <- "SCT"
VariableFeatures(ESTIMATE.merge) <- c(VariableFeatures(LUAD.merge))
View(ESTIMATE.merge@meta.data)

#### SPATIALLY PLOTTING METRICS FROM ESTIMATE ON SAME SCALE FOR ALL SAMPLES ####
SpatialPlot(ESTIMATE.merge, features = "ImmuneScore", ncol = 4)
SpatialPlot(ESTIMATE.merge, features = "TumorPurity", ncol = 4)
SpatialPlot(ESTIMATE.merge, features = "StromalScore", ncol = 4)

#### MAPPING "IC" NUMBERS FROM LUAD.merge OBJECT ONTO NEW ESTIMATE.merge OBJECT ####
ESTIMATE.merge@meta.data <- merge(ESTIMATE.merge@meta.data, LUAD.merge$ICs, by = 0)
row.names(ESTIMATE.merge@meta.data) <- ESTIMATE.merge@meta.data[["Row.names"]]
ESTIMATE.merge@meta.data[["Row.names"]] <- NULL

#### REORDERING ESTIMATE.merge OBJECT TO MATCH ORDER OF LUAD.merge OBJECT ####
matchedorder <- match(rownames(LUAD.merge@meta.data), rownames(ESTIMATE.merge@meta.data))
ESTIMATE.merge@meta.data <- ESTIMATE.merge@meta.data[matchedorder,, drop=F]


#### SAVING MERGED SEURAT OBJECT AFTER ESTIMATE WORKFLOW ####
saveRDS(ESTIMATE.merge, file = "/Users/noahshulkin/Library/CloudStorage/OneDrive-CityofHopeNationalMedicalCenter/Marconett Lab/Visium Files/ESTIMATE.RDS")
ESTIMATE.merge <- readRDS("/Users/noahshulkin/Library/CloudStorage/OneDrive-CityofHopeNationalMedicalCenter/Marconett Lab/Visium Files/ESTIMATE.RDS")


#### ADDING ESTIMATE ASSAY INTO LUAD.merge ASSAYS ####
estimate.merged.assay <- GetAssayData(object =  ESTIMATE.merge[['ESTIMATE']], layer = "data")
LUAD.merge[["ESTIMATE"]] <- CreateAssayObject(data = estimate.merged.assay)
#### SAVING LUAD.merge OBJECT AS A SEPARATE RDS FILE ####
saveRDS(LUAD.merge, file = "/Users/noahshulkin/Library/CloudStorage/OneDrive-CityofHopeNationalMedicalCenter/Marconett Lab/Visium Files/LUADwithESTIMATE.RDS")
LUAD.merge <- readRDS("/Users/noahshulkin/Library/CloudStorage/OneDrive-CityofHopeNationalMedicalCenter/Marconett Lab/Visium Files/LUADwithESTIMATE.RDS")

#### PLOTTING METRICS FROM ESTIMATE FOR ALL SAMPLES ####
DefaultAssay(LUAD.merge) <- "ESTIMATE"
VlnPlot(LUAD.merge, feature = "ImmuneScore")
VlnPlot(LUAD.merge, feature = "TumorPurity")
VlnPlot(LUAD.merge, feature = "StromalScore")
LUAD.nocontrols <- subset(LUAD.merge, subset = (cancer != "normal"))

FeaturePlot(LUAD.merge, reduction = "umap", label = F, features = "ImmuneScore")
FeaturePlot(LUAD.merge, reduction = "umap", label = F, features = "TumorPurity", cols = c("gray", "red"))
FeaturePlot(LUAD.merge, reduction = "umap", label = F, features = "StromalScore", cols = c("gray", "green"))

VlnPlot(LUAD.nocontrols, feature = "ImmuneScore", group.by = "orig.ident") + NoLegend()
VlnPlot(LUAD.nocontrols, feature = "TumorPurity", group.by = "orig.ident") + NoLegend()
VlnPlot(LUAD.nocontrols, feature = "StromalScore", group.by = "orig.ident") + NoLegend()

VlnPlot(LUAD.nocontrols, feature = "ImmuneScore", group.by = "cell.of.origin") + NoLegend()
VlnPlot(LUAD.nocontrols, feature = "TumorPurity", group.by = "cell.of.origin") + NoLegend()
VlnPlot(LUAD.nocontrols, feature = "StromalScore", group.by = "cell.of.origin") + NoLegend()

#### BIOSTATISTICS FOR IMMUNE SCORE ####
View(LUAD.nocontrols@meta.data)
DefaultAssay(LUAD.nocontrols) <- "ESTIMATE"

est_AT2 <- data.matrix(GetAssayData(subset(LUAD.nocontrols, subset = (cell.of.origin == "Sftpc"))[["ESTIMATE"]]))
est_AT2 <- t(est_AT2)
AT2_immune <- as.vector(est_AT2[, 2])

est_AT1 <- data.frame(GetAssayData(subset(LUAD.nocontrols, subset = (cell.of.origin == "Gramd2"))[["ESTIMATE"]]))
est_AT1 <- t(est_AT1)
AT1_immune <- as.vector(est_AT1[,2])

t.test(AT2_immune, AT1_immune) #p-value < 2.2e-16

#### PREPPING FOR CIBERSORTx ####
CSx <- LUAD.merge
DefaultAssay(CSx) <- "SCT"
IC <- data.frame(CSx$ICs)
txt <- GetAssayData(CSx[["SCT"]], layer = "counts")
txt <- data.matrix(txt)
txt <- t(txt)
txt <- merge(IC, txt, by = 0)
row.names(txt) <- txt[["Row.names"]]
txt[["Row.names"]] <- NULL

#### CREATING THE INPUT FOR CIBERSORTx ####
##### ACCORDING TO JING, I MUST FIND THE SUM OF THE "COUNT" LAYER FOR EVERY IC ####
integratedclusters <- unique(txt[,"CSx.ICs"])
for (i in integratedclusters) {
  if (i == "IC1"){
  singleIC <- txt[txt$CSx.ICs == i,]
  singleIC[["CSx.ICs"]] <- NULL
  ICtotal <- colSums(singleIC)
  ICtotal <- data.matrix(ICtotal)
  colnames(ICtotal) <- i
  CSx.txt <- ICtotal
  View(CSx.txt)
  } else{
  singleIC <- txt[txt$CSx.ICs == i,]
  singleIC[["CSx.ICs"]] <- NULL
  ICtotal <- colSums(singleIC)
  ICtotal <- data.matrix(ICtotal)
  colnames(ICtotal) <- i
  CSx.txt <- cbind(CSx.txt, ICtotal)
  }
}


#### REORDERING THE CIBERSORTx MATRIX AND SAVING TO CSV ####
colorder <- c("IC1", "IC2", "IC3", "IC4", "IC5", "IC6", "IC7", "IC8", "IC9", "IC10", "IC11", "IC12", "IC13", "IC14", "IC15", "IC16", "IC17")
CSx.txt <- CSx.txt[, colorder]
write.csv(CSx.txt, file = "/Users/noahshulkin/Library/CloudStorage/OneDrive-CityofHopeNationalMedicalCenter/Marconett Lab/Visium Files/CIBERSORTxPrep.csv")

#### UPLOADING SINGLE CELL REFERENCE FROM MINXIAO YANG####
scref <- readRDS("/Users/noahshulkin/Library/CloudStorage/OneDrive-CityofHopeNationalMedicalCenter/Marconett Lab/Visium Files/sc_highFidelity.RDS")
Idents(object = scref) <- "predicted.celltype_level2"
scref <- subset(scref, downsample = 25)
table(Idents(scref))
table(scref$predicted.celltype_level2)
sctypes <- data.frame(scref$predicted.celltype_level2)
screftxt <- GetAssayData(scref[["RNA"]], layer = "counts")
screftxt <- data.matrix(screftxt)
screftxt <- t(screftxt)
screftxt <- merge(sctypes, screftxt, by = 0)
row.names(screftxt) <- screftxt[["Row.names"]]
screftxt[["Row.names"]] <- NULL
screftxt <- t(screftxt)
screftxt <- data.frame(screftxt)
screftxt <- row_to_names(screftxt, 1)
write.csv(screftxt, file= "screftxt.csv")

#### USING ORTHOGENE TO CHANGE MOUSE GENES TO HUMAN ORTHOLOGS ####
if (!requireNamespace("BiocManager", quietly = TRUE))
  install.packages("BiocManager")
# orthogene is only available on Bioconductor>=3.14
if(BiocManager::version()<"3.14") BiocManager::install(version = "3.14")

BiocManager::install("orthogene")
library(orthogene)

LM22 <- read.csv("/Users/noahshulkin/Library/CloudStorage/OneDrive-CityofHopeNationalMedicalCenter/Marconett Lab/Visium Files/CIBERSORTx/LM22.csv")
rownames(LM22) <- LM22[["Gene.symbol"]]
LM22[["Gene.symbol"]] <- NULL

LM22M <- orthogene::convert_orthologs(gene_df = LM22,
                                        gene_input = "rownames", 
                                        gene_output = "rownames", 
                                        input_species = "human",
                                        output_species = "mouse",
                                        non121_strategy = "drop_output_species",
                                        method = "gprofiler") 
LM22M <- data.matrix(LM22M)
write.csv(LM22M, "/Users/noahshulkin/Library/CloudStorage/OneDrive-CityofHopeNationalMedicalCenter/Marconett Lab/Visium Files/CIBERSORTx/LM22M.csv")

#### REPEATING ORTHOLOG CONVERSION WITH LM22 GEP FILE ####
LM22_GEP <- read.csv("/Users/noahshulkin/Library/CloudStorage/OneDrive-CityofHopeNationalMedicalCenter/Marconett Lab/Visium Files/CIBERSORTx/LM22_source_GEPs.csv")
rownames(LM22_GEP) <- LM22_GEP[["genesinput"]]
LM22_GEP[["genesinput"]] <- NULL

LM22M_GEP <- orthogene::convert_orthologs(gene_df = LM22_GEP,
                                      gene_input = "rownames", 
                                      gene_output = "rownames", 
                                      input_species = "human",
                                      output_species = "mouse",
                                      non121_strategy = "drop_output_species",
                                      method = "gprofiler") 
LM22M_GEP <- data.matrix(LM22M_GEP)
write.csv(LM22M_GEP, "/Users/noahshulkin/Library/CloudStorage/OneDrive-CityofHopeNationalMedicalCenter/Marconett Lab/Visium Files/CIBERSORTx/LM22M_GEP.csv")

#### PERFORMING SINGLE CELL INTEGRATION FROM SEURAT ####
#### FYI, THESE STEPS WILL REQUIRE A HIGH-PERFORMANCE COMPUTING ####
myel <- readRDS("/Users/noahshulkin/Library/CloudStorage/OneDrive-CityofHopeNationalMedicalCenter/Marconett Lab/Visium Files/Myeloid_Annotated.RDS")
scref <- readRDS("/Users/noahshulkin/Library/CloudStorage/OneDrive-CityofHopeNationalMedicalCenter/Marconett Lab/Visium Files/sc_highFidelity.RDS")
rownames(myel@meta.data) <- sub(".*?_", "", rownames(myel@meta.data))

table(scref$predicted.celltype_level2)
View(scref@meta.data)
myel_ann <- data.frame(myel$Celltypes)
myel_ann <- myel_ann[c("myel.Celltypes")]

scref_ann <- data.frame(scref$predicted.celltype_level2)
scref_ann <- scref_ann[c("scref.predicted.celltype_level2")]
colnames(scref_ann)[1] <- "myel.Celltypes"

library(tibble)
scref_ann <- scref_ann %>%
  rownames_to_column() %>%
  rows_update(myel_ann %>% rownames_to_column(), by = "rowname", unmatched = "ignore") %>%
  column_to_rownames()

View(scref@meta.data)
scref@meta.data <- merge(scref@meta.data, scref_ann, by = 0)
rownames(scref@meta.data) <- scref$Row.names
scref@meta.data$Row.names <- NULL
View(scref@meta.data)
saveRDS(scref, "/Users/noahshulkin/Library/CloudStorage/OneDrive-CityofHopeNationalMedicalCenter/Marconett Lab/Visium Files/new_scref.rds")

scref <- SCTransform(scref, ncells = 3000, verbose = F)
DimPlot(scref, group.by = "myel.Celltypes", label = F)

DefaultAssay(LUAD.merge) <- "SCT"
anchors <- FindTransferAnchors(reference = scref, reference.assay = "SCT", query = LUAD.merge, normalization.method = "SCT")
integration.assay <- TransferData(anchorset = anchors, refdata = scref$myel.Celltypes, prediction.assay = T, weight.reduction =  LUAD.merge[["pca"]], dims = 1:30)
LUAD.merge[["Integration"]] <- integration.assay
DefaultAssay(LUAD.merge) <- "Integration"
View(LUAD.merge@meta.data)
saveRDS(LUAD.merge, "/Volumes/NO NAME/LUADwithESTIMATE.RDS")

#### VISUALIZING SCRNASEQ INTEGRATION ####
SpatialPlot(LUAD.merge, features = "AT1/AT2", ncol = 4)
SpatialPlot(LUAD.merge, features = "Macrophages", ncol = 4, max.cutoff = 0.1)

FeaturePlot(LUAD.merge, features = c("Macrophages"))
FeaturePlot(LUAD.merge, features = c("AT1/AT2"))

VlnPlot(LUAD.merge, features = "AM", group.by = ("cell.of.origin"))


#### WRANGLING INTEGRATION DATA TO MAKE STACKED BAR CHART ####
integration <- GetAssayData(LUAD.merge[["Integration"]])
integration <- t(integration)
integration <- integration[, colnames(integration) != "max"]
integration <- data.frame(integration)
integration <- merge(LUAD.merge@meta.data, integration, by = 0)
canceronly <- subset(integration, subset = (integration$cancer != "normal"))
canceronly$iMONs <- rowSums(canceronly[, c("iMONs", "iMON")])
canceronly$iMON <- NULL
canceronly <- pivot_longer(canceronly,
                           cols = "AT1.AT2":"Basophil",
                           names_to = "Variable",
                           values_to = "Value")

ggplot(canceronly, aes(fill=Variable, y=Value, x=cell.of.origin)) +
  geom_bar(position="stack", stat="identity")


#### PUTTING GRANULAR CELL TYPES INTO BROADER CATEGORIES AND VISUALIZING ####
Classes <- canceronly %>% mutate(class = case_when(
  Variable == "B" | Variable == "CD4.T" | Variable == "CD8.T"  | Variable == "DCs" | Variable == "HSPCs" | Variable == "Granulocytes" | Variable == "Macrophages" | Variable == "MDSCs" | Variable == "Megakaryocyte.Platelet" | Variable == "NK" | Variable == "Monocytes" | Variable == "Proliferating.Myeloid.Cells" | Variable == "Treg" | Variable == "AM" | Variable == "ILC" | Variable == "cDC1" | Variable == "cDC2" | Variable == "IM" | Variable == "iMONs" | Variable == "Basophil" | Variable == "maDC" | Variable == "TAMs" ~ "Immune",
  Variable == "AT1" | Variable == "AEC" | Variable == "AT1.AT2"  | Variable == "AT2" | Variable == "PNEC" | Variable == "Deuterosomal" | Variable == "Ciliated" | Variable == "Basal" | Variable == "Secretory" | Variable == "Sox9.Epi" ~ "Epithelial",
  Variable == "VEC" | Variable == "CAP1.EPC" | Variable == "CAP2"  | Variable == "LEC" | Variable == "EPC"  ~ "Endothelial",
  Variable == "ASMC" | Variable == "VSMC" | Variable == "SCMF" | Variable == "Pericyte" | Variable == "PMP" | Variable == "AF1" | Variable == "AF2" | Variable == "Mesothelial" ~ "Mesenchymal"
)
)

ggplot(Classes, aes(fill=class, y=Value, x=cell.of.origin)) +
  geom_bar(position="stack", stat="identity") +
  theme(
    panel.background = element_rect(fill = "transparent", colour = NA), # Remove background
    plot.background = element_rect(fill = "transparent", colour = NA)   # Remove background from the plot area
  ) 

ggplot(Classes, aes(fill=class, y=Value, x=cell.of.origin)) +
  geom_bar(position="fill", stat="identity") +
  theme(
    panel.background = element_rect(fill = "transparent", colour = NA), # Remove background
    plot.background = element_rect(fill = "transparent", colour = NA)   # Remove background from the plot area
  ) 

#### SUBSETTING ON IMMUNE CELLS ONLY AND VISUALIZING WITH STACKED BAR CHART ####
immune_only <- subset(Classes, subset = (class == "Immune"))

ggplot(immune_only, aes(fill=Variable, y=Value, x=cell.of.origin)) +
  geom_bar(position="stack", stat="identity") +
  ggtitle("Immune") +
  theme(
    panel.background = element_rect(fill = "transparent", colour = NA), # Remove background
    plot.background = element_rect(fill = "transparent", colour = NA)   # Remove background from the plot area
  ) 

ggplot(immune_only, aes(fill=Variable, y=Value, x=cell.of.origin)) +
  geom_bar(position="fill", stat="identity") +
  ggtitle("Immune") +
  theme(
    panel.background = element_rect(fill = "transparent", colour = NA), # Remove background
    plot.background = element_rect(fill = "transparent", colour = NA)   # Remove background from the plot area
  ) 

aggregate(Value ~ Variable, data = immune_only, sum)


#### LOOKING AT MYELOID COMPARTMENT WITHIN IMMUNE POPULATION ####
myel_variable = c("IM", "iMONs", "AM", "TAMs", "Granulocytes", "Monocytes", "Macrophages", "HSPCs", "Immature.Macrophages", "MDSCs", "Proliferating.Myeloid.Cells")
myeloid_only <- subset(immune_only, subset = (Variable %in% myel_variable))
table(myeloid_only$Variable)

myelpocket <- myeloid_only %>%
  group_by(cell.of.origin, Variable) %>%
  summarise(mean_value = mean(Value),
          sd_value = sd(Value),
          .groups = 'drop')

myel_error = myelpocket %>%
  arrange(cell.of.origin, desc(Variable)) %>%
  group_by(cell.of.origin) %>%
  mutate(mean_new = cumsum(mean_value)) %>%
  ungroup()

ggplot(myelpocket, aes(x = cell.of.origin, y = mean_value)) +
  geom_bar(stat = 'identity', aes (fill = Variable)) +
  geom_errorbar(data = myel_error, 
                aes(x = cell.of.origin, ymax = mean_new + sd_value, ymin = mean_new - sd_value),
                width =  0.2)

ggplot(myeloid_only, aes(fill=Variable, y=Value, x=cell.of.origin)) +
  geom_bar(position="stack", stat="identity") +
  ggtitle("Myeloid") +
  theme(
    panel.background = element_rect(fill = "transparent", colour = NA), # Remove background
    plot.background = element_rect(fill = "transparent", colour = NA)   # Remove background from the plot area
  ) + scale_fill_manual(values = c("HSPCs" = "#F8766D", 
                                   "iMONs" = "#ABA300",
                                   "MDSCs" = "#7CAE00",
                                   "Proliferating.Myeloid.Cells" = "#00BE67",
                                   "Macrophages" = "#00BFC4",
                                   "TAMs" = "#00A9FF",
                                   "DCs" = "#C77CFF",
                                   "Granulocytes" = "#FF61CC"))

ggplot(myeloid_only, aes(fill=Variable, y=Value, x=cell.of.origin)) +
  geom_bar(position="fill", stat="identity") +
  ggtitle("Myeloid") +
  theme(
    panel.background = element_rect(fill = "transparent", colour = NA), # Remove background
    plot.background = element_rect(fill = "transparent", colour = NA)   # Remove background from the plot area
  ) + scale_fill_manual(values = c("HSPCs" = "#F8766D", 
                                   "iMONs" = "#ABA300",
                                   "MDSCs" = "#7CAE00",
                                   "Proliferating.Myeloid.Cells" = "#00BE67",
                                   "Macrophages" = "#00BFC4",
                                   "TAMs" = "#00A9FF",
                                   "DCs" = "#C77CFF",
                                   "Granulocytes" = "#FF61CC"))

aggregate(Value ~ Variable, data = myeloid_only, sum)
# Results from above function 
#Variable      Value
#1           AM   1.371978
#2 Granulocytes   0.000000
#3        HSPCs   1.812683
#4           IM   0.000000
#5  Macrophages  69.438628
#6        MDSCs 103.924353
#7         TAMs   0.000000
#### LOOKING AT MACROPHAGES AND MDSCS ONLY BETWEEN LUAD MODELS ####
supp <- subset(immune_only, subset = (Variable == "MDSCs" | Variable == "Macrophages"))

ggplot(supp, aes(fill=Variable, y=Value, x=cell.of.origin)) +
  geom_bar(position="dodge", stat="identity") +
  ggtitle("Suppression")

ggplot(supp, aes(fill=Variable, y=Value, x=cell.of.origin)) +
  geom_bar(position="fill", stat="identity") +
  ggtitle("Suppression")

#### BIOSTATISTICS FOR MP/MDSC COMPARISONS BETWEEN LUAD MODELS ####
AT2_MP <- subset(supp, subset = (cell.of.origin == "Sftpc" & Variable == "Macrophages"))
AT2_MP <- AT2_MP$Value
AT2_MP_nonzero <- subset(AT2_MP, subset = (AT2_MP > 0))
AT2_MDSC <- subset(supp, subset = (cell.of.origin == "Sftpc" & Variable == "MDSCs"))
AT2_MDSC <- AT2_MDSC$Value
AT2_MDSC_nonzero <- subset(AT2_MDSC, subset = (AT2_MDSC > 0))
AT1_MP <- subset(supp, subset = (cell.of.origin == "Gramd2" & Variable == "Macrophages"))
AT1_MP <- AT1_MP$Value
AT1_MP_nonzero <- subset(AT1_MP, subset = (AT1_MP > 0))
AT1_MDSC <- subset(supp, subset = (cell.of.origin == "Gramd2" & Variable == "MDSCs"))
AT1_MDSC <-AT1_MDSC$Value
AT1_MDSC_nonzero <- subset(AT1_MDSC, subset = (AT1_MDSC > 0))

library(nortest)
ad.test(AT2_MP) #p-value < 2.2e-16
ad.test(AT1_MP) #p-value < 2.2e-16
ad.test(AT2_MDSC) #p-value < 2.2e-16
ad.test(AT1_MDSC) #p-value < 2.2e-16

wilcox.test(AT2_MP, AT1_MP) #p-value = 5.301e-05
wilcox.test(AT2_MDSC, AT1_MDSC) #p-value < 2.2e-16

MP <- subset(supp, subset = (Variable == "Macrophages"))
MP <- MP[, c("Row.names", "cell.of.origin", "Value")]
colnames(MP)[3] <- "MP"

MDSC <- subset(supp, subset = (Variable == "MDSCs"))
MDSC <- MDSC[, c("Row.names", "cell.of.origin", "Value")]
colnames(MDSC)[3] <- "MDSC"

supp_df <- merge(MP, MDSC, by = "Row.names")

data_summary <- function(data, varname, groupnames){
  require(plyr)
  summary_func <- function(x, col){
    c(mean = mean(x[[col]], na.rm=TRUE),
      sd = sd(x[[col]], na.rm=TRUE))
  }
  data_sum<-ddply(data, groupnames, .fun=summary_func,
                  varname)
  data_sum <- rename(data_sum, c("mean" = varname))
  return(data_sum)
}

MDSC_summary <- data_summary(supp_df, varname = "MDSC", groupnames = "cell.of.origin.x")

ggplot(MDSC_summary, aes(x=cell.of.origin.x, y=MDSC)) + 
  geom_bar(stat="identity", position=position_dodge(), fill= c('red','purple')) +
  geom_errorbar(aes(ymin=MDSC-sd, ymax=MDSC+sd), width=.2,
                position=position_dodge(.9)) +
  labs(title = "% MDSCs Within Visium Spot", x = "Model", y = "Percentage") + theme_classic() +
  geom_signif(y_position = c(0.075), xmin = c(1), 
              xmax = c(2), annotation = c("***"),
              tip_length = 0)

MP_summary <- data_summary(supp_df, varname = "MP", groupnames = "cell.of.origin.x")

ggplot(MP_summary, aes(x=cell.of.origin.x, y=MP)) + 
  geom_bar(stat="identity", position=position_dodge(), fill= c('red','purple')) +
  geom_errorbar(aes(ymin=MP-sd, ymax=MP+sd), width=.2,
                position=position_dodge(.9)) +
  labs(title = "% MPs Within Visium Spot", x = "Model", y = "Percentage") + theme_classic() +
  geom_signif(y_position = c(0.025), xmin = c(1), 
              xmax = c(2), annotation = c("***"),
              tip_length = 0)

MP_positive <- subset(LUAD.merge, subset = ((cancer != "normal") & (Macrophages > 0)))
shapiro.test(AT2_MP_nonzero) #p-value < 2.2e-16
shapiro.test(AT1_MP_nonzero) #p-value < 2.2e-16

VlnPlot(MP_positive, group.by = "cell.of.origin", features = "Macrophages", cols = c("#e31e24", "#00a0e3")) + stat_compare_means(method = "wilcox.test", label.x = 1.5)
table(MP_positive$cell.of.origin)

#### ISOLATING SPOTS WITH TUMOR BASED ON ESTIMATE TUMORPURITY SCORE ####
DefaultAssay(LUAD.merge) <- "ESTIMATE"
SpatialFeaturePlot(LUAD.merge, ncol = 4, features = c("TumorPurity"), min.cutoff = 0.8)

#### VISUALZING HIGH TUMOR PURITY SCORE ####
FeaturePlot(LUAD.merge, reduction = "umap", features = c("TumorPurity"), cols = c("lightgrey", "red2"), order = TRUE, min.cutoff = 0.84)
LUAD.tumor <- subset(LUAD.merge, subset = (TumorPurity > 0.84)) ## TumorPurity score of 0.84 excludes 99% of spots from controls
DimPlot(LUAD.tumor, reduction = "umap") + ggtitle("Tumor Purity > 0.84")
DimPlot(LUAD.tumor, reduction = "umap", group.by = "cell.of.origin") + ggtitle("Tumor Purity > 0.84")
table(LUAD.tumor$ICs)
DefaultAssay(LUAD.tumor) <- "SCT"
FeaturePlot(LUAD.tumor, reduction = "umap", features = c("Foxj1", "Dnah5"), blend = T)
DotPlot(LUAD.tumor, features =  c("Foxj1", "Dnah5")) ### BIA Histology is localized to IC2 and IC16
View(LUAD.tumor@meta.data)

#### FILTERING OUT BIA HISTOLOGY THAT IS LOCALIZED IN IC2 AND IC16 ####
LUAD.noBIA <- subset(LUAD.tumor, subset = (ICs != "IC16"))
LUAD.noBIA <- subset(LUAD.noBIA, subset = (ICs != "IC2"))
DimPlot(LUAD.noBIA, reduction = "umap") + ggtitle("Tumor without BIA")
DimPlot(LUAD.noBIA, reduction = "umap", group.by = "cell.of.origin") + ggtitle("Tumor without BIA")

#### FILTERING SPOTS FOR ONES THAT ONLY INCLUDE MACROPHAGES BASED ON SCRNASEQ INTEGRATION ####
DefaultAssay(LUAD.merge) <- "Integration"
LUAD.MP <- subset(LUAD.merge, subset = (Macrophages > 0))
DefaultAssay(LUAD.MP) <- "Integration"

#### MAKING METADATA COLUMNS FOR MP-POSITIVE AND TUMOR-POSITIVE SPOTS BASED ON PRE-PROCESSING FOR THESE CELL TYPES ####
MP_spots <- rownames(LUAD.MP@meta.data)
Tumor_spots <- rownames(LUAD.noBIA@meta.data)
LUAD.merge@meta.data <- LUAD.merge@meta.data %>% mutate(MP.Tumor = ifelse(rownames(LUAD.merge@meta.data) %in% MP_spots, "Macrophage", ifelse(rownames(LUAD.merge@meta.data) %in% Tumor_spots, "Tumor", "NA")))


cols <- c("gray", "red", "blue")
Idents(LUAD.merge) <- LUAD.merge$MP.Tumor
names(cols) <- unique(Idents(LUAD.merge))
SpatialDimPlot(LUAD.merge, group.by = "MP.Tumor", ncol = 4, cols = cols)


#### SAVING LUAD.merge OBJECT AS A SEPARATE RDS FILE ####
saveRDS(LUAD.merge, file = "/Users/noahshulkin/Library/CloudStorage/OneDrive-CityofHopeNationalMedicalCenter/Marconett Lab/Visium Files/LUADwithMPTumor.RDS")
LUAD.merge <- readRDS("/Users/noahshulkin/Library/CloudStorage/OneDrive-CityofHopeNationalMedicalCenter/Marconett Lab/Visium Files/LUADwithMPTumor.RDS")

#### DETERMINING IMMUNOSUPPRESSION OF MACROPHAGE POPULATION ####
LUAD.MP.canceronly <- subset(LUAD.MP, subset = (cancer != "normal"))
DefaultAssay(LUAD.MP.canceronly) <- "SCT"
MP_supp <- list(c("Spp1", "Apoe", "Gpnmb", "Trem2", "Il10", "Il13", "Il33", "Vegf", "Csf1", "Ccl2", "Ccl5", "Ccl17", "Ccl22", "Tgfb", "Rgf", "Pdgf", "Arg1", "Fibrin", "Wnt", "MMPs", "Vcan", "Cxcl8", "Csf1", "Csf3", "Cd47", "Enpp1", "Cgas", "Sting1"))
MP_suppScore <- AddModuleScore(LUAD.MP.canceronly, MP_supp, ctrl = 100, name = "Macrophage Suppression Score")
VlnPlot(MP_suppScore, feature = "Macrophage Suppression Score1", group.by = "cell.of.origin", cols = c("#e31e24", "#00a0e3")) + stat_compare_means(method = "wilcox.test")

ggviolin(MP_suppScore@meta.data, 
         x="cell.of.origin", y="Macrophage Suppression Score1", 
         width = 0.8, color = "black",
         fill = "cell.of.origin",
         xlab = F, 
         add = 'mean_sd', 
         bxp.errorbar=T,
         bxp.errorbar.width=0.05,
         size=0.5, 
         legend = "right") +
  scale_fill_manual(values = c("Sftpc" = "#00a0e3",
                               "Gramd2" = "#e31e24")) +
  RotatedAxis() #+ stat_compare_means(method = "wilcox.test", label.x = 1.5)

table(MP_suppScore$cancer)
AT1_MPsupp <- subset(MP_suppScore@meta.data, subset = (cell.of.origin == "Gramd2"))
AT2_MPsupp <- subset(MP_suppScore@meta.data, subset = (cell.of.origin == "Sftpc"))

shapiro.test(AT1_MPsupp$`Macrophage Suppression Score1`) #p-value = 0.6814
shapiro.test(AT2_MPsupp$`Macrophage Suppression Score1`) #p-value = 0.01022
ad.test(AT2_MPsupp$`Macrophage Suppression Score1`)

MP_react <- list(c('Csf2', 'Isg15', 'Il1a', 'Il1b', 'Il6', 'Il8', 'Il22', 'Il23a', 'Cxcl3', 'Cxcl9', 'Cxcl10', 'Ccl4', "Cd80", "Cd86", "Il1b", "Il2", "Il18"))
MP_reactScore <- AddModuleScore(LUAD.MP.canceronly, MP_react, ctrl = 100, name = "Macrophage Immunoreactivity Score")
VlnPlot(MP_reactScore, feature = "Macrophage Immunoreactivity Score1", group.by = "cell.of.origin", cols = c("#e31e24", "#00a0e3")) + stat_compare_means(method = "wilcox.test")

AT1_MPreact <- subset(MP_reactScore@meta.data, subset = (cell.of.origin == "Gramd2"))
AT2_MPreact <- subset(MP_reactScore@meta.data, subset = (cell.of.origin == "Sftpc"))

shapiro.test(AT1_MPreact$`Macrophage Immunoreactivity Score1`) #p-value = 8.64e-07
shapiro.test(AT2_MPreact$`Macrophage Immunoreactivity Score1`) #p-value = 2.276e-09

ggviolin(MP_reactScore@meta.data, 
         x="cell.of.origin", y="Macrophage Immunoreactivity Score1", 
         width = 0.8, color = "black",
         fill = "cell.of.origin",
         xlab = F, 
         add = 'mean_sd', 
         bxp.errorbar=T,
         bxp.errorbar.width=0.05,
         size=0.5, 
         legend = "right") +
  scale_fill_manual(values = c("Sftpc" = "#00a0e3",
                               "Gramd2" = "#e31e24")) +
  RotatedAxis() + stat_compare_means(method = "wilcox.test")

### MAKING TERTIARY LYMPHOID STRUCTURE MODULE SCORE ####
#### genes used for TLS signature is from https://www.cell.com/immunity/pdf/S1074-7613(22)00081-4.pdf ####
DefaultAssay(LUAD.merge) <- "SCT"
TLS <- list(c("Igha", "Ighg1", "Ighg3", "Ighm", "Igkc", "Iglc1", "Iglc2", "Iglc3", "Jchain", "Cd79a", "Fcrl5", "Mzb1", "Ssr4", "Xbp1", "Trbc2", "Il7r", "Cxcl12", "Lum", "C1qa", "C7", "Cd52", "Apoe", "Pim2", "Derl3", "Timd4", "Cd163", "Folr2"))
TLS_Score <- AddModuleScore(LUAD.merge, TLS, ctrl = 100, name = "TLS Score")
View(TLS_Score@meta.data)
FeaturePlot(TLS_Score, features = "TLS Score1", reduction = "umap", min.cutoff = 0.2)
SpatialFeaturePlot(TLS_Score, features = "TLS Score1", ncol = 4)
TLS_Score_canceronly <- subset(TLS_Score, subset = (cancer != "normal"))
VlnPlot(TLS_Score_canceronly, feature = "TLS Score1", group.by = "cell.of.origin", cols = c("#e31e24", "#00a0e3")) + stat_compare_means(method = "wilcox.test", label.x = 1.5)

AT1_TLS <- subset(TLS_Score_canceronly, subset = (cell.of.origin == "Gramd2"))
AT2_TLS <- subset(TLS_Score_canceronly, subset = (cell.of.origin == "Sftpc"))

ad.test(AT1_TLS$`TLS Score1`) #p-value < 2.2e-16
ad.test(AT2_TLS$`TLS Score1`) #p-value < 2.2e-16

ggviolin(TLS_Score_canceronly@meta.data, 
         x="cell.of.origin", y="TLS Score1", 
         width = 0.8, color = "black",
         fill = "cell.of.origin",
         xlab = F, 
         add = 'mean_sd', 
         bxp.errorbar=T,
         bxp.errorbar.width=0.05,
         size=0.5, 
         legend = "right") +
         scale_fill_manual(values = c("Sftpc" = "#00a0e3",
                                      "Gramd2" = "#e31e24")) +
         RotatedAxis() + stat_compare_means(method = "wilcox.test")

##### STJOINCOUNT FOR EVERY SAMPLE ####
load("/Users/noahshulkin/Library/CloudStorage/OneDrive-CityofHopeNationalMedicalCenter/Marconett Lab/Visium Files/ESTIMATE.RData")
samplelist <- c(Sftpc_Cre, Sftpc_lung1, Sftpc_lung2, Sftpc_lung3, Gramd2_Cre, Gramd2_lung1, Gramd2_lung2, Gramd2_lung3)

library(stringr)
library(stJoincount)
library(pheatmap)

mapped_samplelist <- c()
for (i in samplelist) {
  print(i)
  DefaultAssay(i) <- "Spatial"
  mptumorlist <- data.frame(LUAD.merge$orig.ident, LUAD.merge$MP.Tumor)
  string <- i@meta.data$orig.ident[1]
  print(string)
  if (grepl("Cre", string)) {
    rows <- paste(i@meta.data$orig.ident, row.names(i@meta.data), sep = "_")
    row.names(i@meta.data) <- rows
  } else {
    rows <- paste(paste(i@meta.data$cell.of.origin, "_Kras_", str_sub(string, -1, -1), sep = ""), row.names(i@meta.data), sep = "_")
    row.names(i@meta.data) <- rows
  }
  i@meta.data <- merge(i@meta.data, mptumorlist, by = 0)
  row.names(i@meta.data) <- i@meta.data[["Row.names"]]
  i@meta.data[["Row.names"]] <- NULL
  mapped_samplelist[[length(mapped_samplelist)+1]] = i
  
  if (grepl("Cre", string)) next
  metadata <- i@meta.data
  names(metadata)[names(metadata) == 'LUAD.merge.MP.Tumor'] <- "Cluster"
  coord <- GetTissueCoordinates(i)
  row.names(coord) <- paste(paste(i@meta.data$cell.of.origin, "_Kras_", str_sub(string, -1, -1), sep = ""), row.names(coord), sep = "_")
  merged <-  merge(metadata, coord, by = 0)
  stLUAD <- merged[c("Row.names", "imagecol", "imagerow", "Cluster")]
  rownames(stLUAD) <- stLUAD$Row.names
  stLUAD <- stLUAD[c("imagecol", "imagerow", "Cluster")]
  
  mosaicIntegration <- rasterizeEachCluster(stLUAD)
  joincount.result <- joincountAnalysis(mosaicIntegration)
  matrix <- zscoreMatrix(stLUAD, joincount.result)
  zscorePlot(matrix)
}

for (i in mapped_samplelist){
  string <- i@meta.data$orig.ident[1]
  if (grepl("Cre", string)) next
  DefaultAssay(i) <- "Spatial"
  metadata <- i@meta.data
  names(metadata)[names(metadata) == 'LUAD.merge.MP.Tumor'] <- "Cluster"
  coord <- GetTissueCoordinates(i)
  row.names(coord) <- paste(paste(i@meta.data$cell.of.origin, "_Kras_", str_sub(string, -1, -1), sep = ""), row.names(coord), sep = "_")
  merged <-  merge(metadata, coord, by = 0)
  final <- merged[c("imagecol", "imagerow", "LUAD.merge.orig.ident", "cell.of.origin", "Cluster")]
  write.csv(final, paste("/Users/noahshulkin/Library/CloudStorage/OneDrive-CityofHopeNationalMedicalCenter/Marconett Lab/Visium Files/squidpy/", i@meta.data$orig.ident[1], ".csv", sep = ""))
}

#### REMAKING MYELOID BAR CHARTS TO INCLUDE CONTROL SAMPLES ####
integration <- GetAssayData(LUAD.merge[["Integration"]])
integration <- t(integration)
integration <- integration[, colnames(integration) != "max"]
integration <- data.frame(integration)
integration <- merge(LUAD.merge@meta.data, integration, by = 0)
integration$iMONs <- rowSums(integration[, c("iMONs", "iMON")])
integration$iMON <- NULL
integration <- pivot_longer(integration,
                           cols = "AT1.AT2":"Basophil",
                           names_to = "Variable",
                           values_to = "Value")

Classes <- integration %>% mutate(class = case_when(
  Variable == "B" | Variable == "CD4.T" | Variable == "CD8.T"  | Variable == "DCs" | Variable == "HSPCs" | Variable == "Granulocytes" | Variable == "Macrophages" | Variable == "MDSCs" | Variable == "Megakaryocyte.Platelet" | Variable == "NK" | Variable == "Monocytes" | Variable == "Proliferating.Myeloid.Cells" | Variable == "Treg" | Variable == "AM" | Variable == "ILC" | Variable == "cDC1" | Variable == "cDC2" | Variable == "IM" | Variable == "iMONs" | Variable == "Basophil" | Variable == "maDC" | Variable == "TAMs" ~ "Immune",
  Variable == "AT1" | Variable == "AEC" | Variable == "AT1.AT2"  | Variable == "AT2" | Variable == "PNEC" | Variable == "Deuterosomal" | Variable == "Ciliated" | Variable == "Basal" | Variable == "Secretory" | Variable == "Sox9.Epi" ~ "Epithelial",
  Variable == "VEC" | Variable == "CAP1.EPC" | Variable == "CAP2"  | Variable == "LEC" | Variable == "EPC"  ~ "Endothelial",
  Variable == "ASMC" | Variable == "VSMC" | Variable == "SCMF" | Variable == "Pericyte" | Variable == "PMP" | Variable == "AF1" | Variable == "AF2" | Variable == "Mesothelial" ~ "Mesenchymal"
)
)

immune_only <- subset(Classes, subset = (class == "Immune"))

myel_variable = c("IM", "iMONs", "AM", "TAMs", "Granulocytes", "Monocytes", "Macrophages", "HSPCs", "Immature.Macrophages", "MDSCs", "Proliferating.Myeloid.Cells")
myeloid_only <- subset(immune_only, subset = (Variable %in% myel_variable))

myeloid_only$cell.of.origin[myeloid_only$cancer == "normal"] <- "Control"
table(myeloid_only$cell.of.origin)

ggplot(myeloid_only, aes(fill=Variable, y=Value, x=cell.of.origin)) +
  geom_bar(position="stack", stat="identity") +
  ggtitle("Myeloid") +
  theme(
    panel.background = element_rect(fill = "transparent", colour = NA), # Remove background
    plot.background = element_rect(fill = "transparent", colour = NA)   # Remove background from the plot area
  ) + scale_fill_manual(values = c("HSPCs" = "#F8766D", 
                                   "iMONs" = "#ABA300",
                                   "MDSCs" = "#7CAE00",
                                   "Proliferating.Myeloid.Cells" = "#00BE67",
                                   "Macrophages" = "#00BFC4",
                                   "TAMs" = "#00A9FF",
                                   "DCs" = "#C77CFF",
                                   "Granulocytes" = "#FF61CC"))

ggplot(myeloid_only, aes(fill=Variable, y=Value, x=cell.of.origin)) +
  geom_bar(position="fill", stat="identity") +
  ggtitle("Myeloid") +
  theme(
    panel.background = element_rect(fill = "transparent", colour = NA), # Remove background
    plot.background = element_rect(fill = "transparent", colour = NA)   # Remove background from the plot area
  ) + scale_fill_manual(values = c("HSPCs" = "#F8766D", 
                                   "iMONs" = "#ABA300",
                                   "MDSCs" = "#7CAE00",
                                   "Proliferating.Myeloid.Cells" = "#00BE67",
                                   "Macrophages" = "#00BFC4",
                                   "TAMs" = "#00A9FF",
                                   "DCs" = "#C77CFF",
                                   "Granulocytes" = "#FF61CC"))
```
